# Can Prediction Error Explain Predictability Effects on the N1 during Picture-Word Verification?

**DOI:** 10.1101/2023.08.07.552265

**Authors:** Jack E. Taylor, Guillaume A. Rousselet, Sara C. Sereno

## Abstract

Do early effects of predictability in visual word recognition reflect prediction error? Electrophysiological research investigating word processing has demonstrated predictability effects in the N1, or first negative component of the event-related potential (ERP). However, findings regarding the magnitude of effects and potential interactions of predictability with lexical variables have been inconsistent. Moreover, past studies have typically used categorical designs with relatively small samples and relied on by-participant analyses. Nevertheless, reports have generally shown that predicted words elicit less negative-going (i.e., lower amplitude) N1s, a pattern consistent with a simple predictive coding account. In our preregistered study, we tested this account via the interaction between prediction magnitude and certainty. A picture-word verification paradigm was implemented in which pictures were followed by tightly matched picture-congruent or picture-incongruent written nouns. The predictability of target (picture-congruent) nouns was manipulated continuously based on norms of association between a picture and its name. ERPs from 68 participants revealed a pattern of effects opposite to that expected under a simple predictive coding framework.

## Introduction

Readers and listeners routinely use context to predict upcoming semantic and lexical content. Evidence for such predictive processes arises from both behavioural and neural correlates of language comprehension (Kuperberg & Jaeger, 2016; Luke & Christianson, 2016; Pickering & Gambi, 2018; Rayner et al., 2011; Van Petten & Luka, 2012), with demonstrated facilitation for the processing of predicted information (Federmeier, 2007; Pickering & Garrod, 2013).

A key question in this area is, how early in the processing stream are predictive processes able to modulate visual word recognition. One early stage in visual word recognition, which may be sensitive to prediction, involves the processing of visual word forms. A word form can be defined as the visual pattern of a single written word, comprised of smaller orthographic components (e.g., letters, letter bigrams, graphemes, strokes). While some electrophysiological evidence suggests sensitivity to orthographic variables in an earlier posterior P1 component peaking at around 100 ms after word presentation (e.g., Nobre et al., 1994; Segalowitz & Zheng, 2009; Sereno et al., 1998), the event-related potential (ERP) component most identified as an index of orthographic processing across different scripts is the first posterior negative-going wave, the N1 (Bentin et al., 1999; Lin et al., 2011; Ling et al., 2019; Maurer, Brandeis, & McCandliss, 2005; Maurer et al., 2008; Pleisch et al., 2019). The N1 is also sometimes referred to as the N170 due to the timing of its peak in some studies, at around 170 ms. This typically occipitotemporal, negative-going component shows reliable differences between orthographic and non-orthographic stimuli (e.g., words elicit more negative-going N1s than false-font strings do; Appelbaum et al., 2009; Bentin et al., 1999; Eberhard-Moscicka et al., 2016; Maurer, Brandeis, & McCandliss, 2005; Maurer, Brem, et al., 2005; Pleisch et al., 2019; Zhao et al., 2014).

Accounts of orthographic processing often stress the importance of top-down predictions, and their interactions with bottom-up sensory input. For instance, the interactive account of the ventral occipito-temporal cortex (vOT), a region which is a likely generator of the N1 ERP component (Allison et al., 1994; Brem et al., 2009; Cohen et al., 2000; Dale et al., 2000; Maurer, Brem, et al., 2005; Nobre et al., 1994; Taha et al., 2013; Woolnough et al., 2021), suggests that sensitivity to orthography arises through the synthesis of bottom-up visuospatial information and top-down predictions informed by prior experience and knowledge (Price & Devlin, 2011). Such accounts exist within a predictive coding framework, according to which the brain utilises higher-level information to build, maintain, and continually update hierarchical series of estimators that form generative models of sensory information (Friston, 2010; Rao & Ballard, 1999; Rauss et al., 2011). Predictive coding accounts have been employed to explain prediction effects observed in early evoked responses across a range of domains, such as the mismatch negativity (Garrido et al., 2009) and sensory attenuation of self-generated percepts (Knolle et al., 2013). A key feature of such accounts is that higher-level predictions cause lower-level features to be preactivated, and that the difference between the bottom-up sensory input and top-down predictions corresponds to a prediction error, which the brain attempts to minimise (Clark, 2013; Walsh et al., 2020).

In a predictive coding framework, prediction errors are commonly determined by two key attributes: the magnitude of the error, and the precision or certainty of the error (Feldman & Friston, 2010; Kanai et al., 2015). Such variants of predictive coding models are commonly referred to as *precision-weighted*. Feldman and Friston (2010) likened the error signal to the calculation of the *t* statistic, where magnitude of an observation (i.e., mean, or mean difference) is divided by the inverse of its precision (i.e., standard error). Prediction errors, weighted by precision in this manner, can be conceptualised as representing the degree of “surprise” associated with a set of observations under a specified hypothesis.

Firstly, the magnitude of the error should determine the size of the error signal, with larger prediction errors resulting from greater mismatch between descending (top-down) predictions and ascending (bottom-up) sensory input. In neutral (non-biasing) contexts, a predictive coding account that includes learning of statistical regularities over extended periods would assert that error signals should vary as a function of stimulus regularity. More specifically, a predictive coding account of orthographic processing would expect error signals to vary as a function of the size of the difference between a general orthographic prior (e.g., an average word form) and a presented word form. Some recent findings appear to support the notion that the N1 reflects a neutral-context error signal, with greater distance from an orthographic prior eliciting greater amplitude (Gagl et al., 2020), while the profile of the N1’s sensitivity to word form regularity over experience matches that expected under a predictive coding account (Huang et al., 2022; Zhao et al., 2019).

Secondly, the precision or certainty of the prediction error should influence the response, with more certain descending predictions, and more certain ascending sensory input, eliciting greater error signals when predictions are violated. In neutral contexts, predictions, and certainty about them, may not be expected to vary much from a context-general prior. Indeed, it is easier to envisage the expected role of prediction precision for orthographic processing in biasing contexts, where precision is more variable than it is in neutral contexts. A predictive coding model of orthographic processing that allows for online, context-informed updating of orthographic priors would expect that the predictability of word forms should influence error responses, with more predictable contexts eliciting stronger prediction error effects. For instance, a sentential context that elicits a clear and reliable prediction for an upcoming word (i.e., that has high Cloze probability) should show a larger prediction error difference, between succeeding prediction-congruent and -incongruent word forms, than should a more neutral sentential context that is consistent with a large number of low-probability candidate words.

In this paper, we examine whether a simple predictive coding account that includes online updating of context-biased predictions and expectations can explain neural activity, captured in the N1, elicited by a word in context. Specifically, we examine whether sensitivity to prediction error in the N1 is dependent on contextual predictability in the manner that a predictive coding account would expect. This question is prompted by (1) the emerging evidence that the N1 in neutral contexts is consistent with an orthographic prediction error signal (Gagl et al., 2020; Huang et al., 2022; Zhao et al., 2019), and (2) existing evidence that biasing semantic contexts can modulate the N1 ERP (outlined below). To address our question, we employ a paradigm informed directly by predictive coding models, manipulating prediction congruency and precision independently, to examine whether the N1 shows the pattern of amplitudes expected under such a model, in biasing contexts. Moreover, we maximise our sensitivity to an orthographic prediction error by presenting prediction-congruent and -incongruent words that are carefully matched item-wise on possible confounders, with maximal orthographic distance from one another. Importantly, evidence for a context-informed prediction error signal at an early, likely orthographic, stage of processing, would not preclude the existence of similar prediction error signals at later stages. Indeed, the hierarchically composed generative model posited by a predictive coding account is fully compatible with the production of prediction errors spanning a hierarchy of linguistic representations.

We hypothesise that according to a simple predictive coding model, the N1 should be larger for prediction-incongruent than prediction-congruent word forms (i.e., prediction error), in a manner dependent on the level of predictability (i.e., precision). We hypothesise that as predictability increases, so too should the prediction error effect.

We begin by reviewing findings from prior studies. We make a distinction between those studies that have biased expectations via *linguistic cues* (text preceding the target word), and those that have employed *non-linguistic cues* (e.g., cross-modal contexts and manipulation of task demands).

### Biasing Word Form Predictions via Linguistic Cues

Readers’ predictions of upcoming word forms are generally manipulated via linguistic cues. In these studies, a target word’s predictability is typically determined in a pre-experiment norming study, operationalised via Cloze probability (i.e., the probability that the target is correctly guessed given its preceding context). Such a measure of word form predictability aligns closely with the concept of prediction precision or certainty in a predictive coding account.

Recent ERP investigations that have manipulated sentential context have also often varied word frequency, with the assumption that an interaction of predictability with word frequency would provide evidence for top-down influences on lexical access. In **Table** 1, we summarise N1 results reported from studies using sentential paradigms that have employed such *Predictability × Frequency* designs. While effects often extend to earlier and later components, we limit our summary to those involving predictability within the N1 window. In Figure 1 we visualise the timing of N1 windows applied in these studies and others cited in this introduction. Sentential studies using a Predictability × Frequency design have demonstrated effects in the N1, although the pattern of effects observed across studies is varied (for a review, see Sereno et al., 2019). We also note that studies using average reference showed more posterior effects, while effects reported from studies using mastoid reference showed more centroparietal topography.

**Figure 1.**
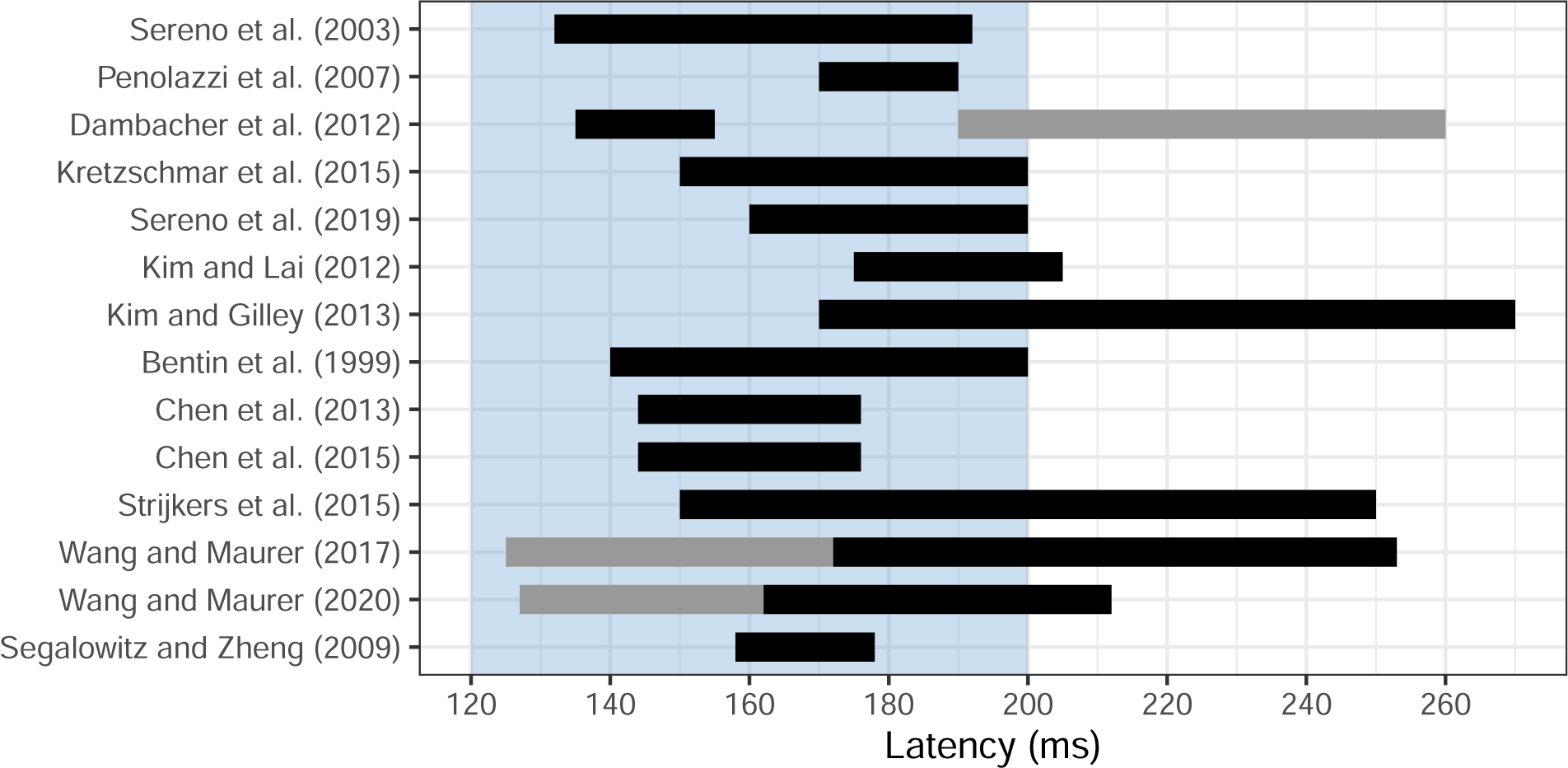
*N1 windows in predictability studies.* Some studies analysed two N1 windows (e.g., onset and offset). N1 windows reported to show a predictability effect are highlighted in black, while N1 windows that failed to show a predictability effect are highlighted in grey. Studies are listed in order of their mention in our review. For reference, the blue region displays the N1 period that we pre-registered.

In addition to such studies that focused on the N1, some studies designed to focus on N400 effects of predictability may also provide insight into early prediction effects. For instance, Brothers et al. (2015) examined correlates of prediction accuracy in the N400, in a sentential design with cloze probability of either medium (.5) or low (<.01) cloze probability. Although they did not report effects in the N1, Brothers et al. did show that accurate predictions of upcoming words were associated with more positive amplitudes after 200 ms, in a P2 component immediately following the N1. In another N400 study using a design related to that employed in the present study, Lau et al. (2016) presented adjective-noun pairs to participants in which the effects of both congruency and predictability were examined, showing small congruency, and large predictability, effects in the N400. As with Brothers et al., Lau et al. report ERPs with no robust differences prior to a P2 component.

**Table 1.** Summary of N1 effects reported in studies that biased word form predictions in sentential paradigms, using a Predictability (low, high) × Frequency (low, high) factorial design.

| Study and Results Summary | SOA | Window | Effect Sites | Reference |
| --- | --- | --- | --- | --- |
| Sereno et al. (2003)<br>More negative amplitudes at lower predictability, but only for low frequency words. | 450 | 132-192 | Posterior & Anterior <sup>a</sup> | Average |
| Penolazzi et al. (2007) <sup>b</sup><br>More negative amplitudes at higher predictability, and no interaction with frequency. | 700 | 170-190 | Centroparietal | Mastoids |
| Dambacher et al. (2012)<br>More negative amplitudes for low than high frequency words, but only at higher predictability. | 280 <sup>c</sup> | 135-155 | Posterior | Average |
| More negative amplitudes for low than high frequency words, and no interaction with predictability. | 280 <sup>c</sup> | 190-260 | Posterior | Average |
| Kretzschmar et al. (2015)<br>More negative amplitudes at higher predictability, and no interaction with frequency. | - <sup>d</sup> | 150-200 | Centroparietal | Mastoids |
| Sereno et al. (2019)<br>More negative amplitudes at higher predictability, but only for high frequency words. | 300 | 160-200 | Left | Average |
| More negative amplitudes at lower predictability, but only for high frequency words. | 300 | 160-200 | Right | Average |

SOA: Stimulus Onset Asynchrony (ms).
Window: Analysed ERP Window (ms).
<sup>a</sup> This topography describes the first factor in a spatial factor analysis.
<sup>b</sup> This study additionally manipulated word length, finding no interaction in the N1.
<sup>c</sup> SOA varied (280, 490, & 700 ms), with N1 effects only observed at the SOA of 280 ms.
<sup>d</sup> This study analysed Fixation-Related Potentials in self-paced reading.

Instead of manipulating error precision or certainty, as the above studies have by varying predictability, Kim and Lai (2012) manipulated the orthographic error *magnitude*. Using a 550 ms SOA, the target word or alternative orthographic versions of it were presented in contexts that were acutely predictive of the target (*M_Cloze_*=.90). Contexts were followed by the predictable target word (e.g., *cake*), an orthographically similar pseudoword (e.g., *ceke*), an orthographically dissimilar pseudoword (e.g., *tont*), or a consonant-string nonword (e.g., *srdt*). Consistent with an orthographic explanation for prediction effects in the N1, relative to targets, N1 (175-205 ms) amplitude was more negative-going for orthographically dissimilar pseudowords and nonwords (i.e., when orthographic prediction error magnitude was greater). Orthographically similar pseudowords, while significantly different from all other conditions in the earlier P1, elicited N1 components more similar in amplitude to target words.

Another linguistic cue that has been manipulated is grammaticality. Kim and Gilley (2013) demonstrated effects of syntactic anomaly on the N1. Sentences leading to a strong prediction for the determiner, *the*, were presented unchanged or with the determiner replaced with an agrammatic preposition (e.g., *The thief was caught by the/for police*). The left-lateralised occipitotemporal N1 (170-270 ms) was more negative-going with the syntactically anomalous preposition than with the determiner. As the authors point out, the N1 effect is unlikely to be evidence for sensitivity to syntax per se. Rather, given evidence of the N1’s sensitivity to orthographic features, it is probably more accurate to posit that the high predictability of the determiner’s orthographic features elicited a less negative-going N1 when these predictions were confirmed.

Kim and Gilley’s simultaneous manipulation of orthography and syntax highlights a prevalent issue within the literature: namely, altering the visual word form necessitates alteration of the semantics, syntax, and/or plausibility of the sentence or wider discourse. Another limitation shared by studies using word-by-word presentation of sentences is that ERPs elicited by the target word can become difficult to disentangle from ERPs elicited by preceding or succeeding words, especially if the SOA is short or unjittered.

While fast presentation times of sentential contexts and targets are useful for demonstrating that early modulation by predictive processes extends to realistic reading rates, their application may not be necessary to demonstrate that such modulation can occur. It is also of note that in a recent review of ERP studies using sentence- and discourse-level contexts to examine early neural correlates of word form prediction, Nieuwland (2019) concluded that findings thus far have been weak, inconsistent, and in need of more replication attempts. Moreover, most studies to date were not pre-registered and often used inappropriate analysis models that did not account for measurement variability, raising questions about false positives in that literature.

### Biasing Word Forms via Non-Linguistic Cues

Effects of prediction and expectation may alternatively be investigated using paradigms that modulate non-linguistic features of tasks and stimuli. In one approach, identical or suitably matched stimuli are presented under different task instructions (e.g., Compton et al., 1991). In that context, tasks requiring more explicit lexical and semantic processing cause words to elicit more negative-going N1s (144-176 ms; Chen et al., 2013). Tasks requiring more in-depth lexicosemantic processing may also increase sensitivity to lexical variables such as word frequency in the N1 (144-176 ms, Chen et al., 2015; 150-250 ms, Strijkers et al., 2015), and may increase the size of script familiarity effects (more negative amplitudes for familiar scripts) (F. Wang & Maurer, 2017). This may be especially in the N1’s offset period (172-253 ms F. Wang & Maurer, 2017), where onsets and offsets are defined respectively as the periods in the component’s time window which precede and succeed its peak. F. Wang and Maurer (2020) further showed that biasing participants’ word form predictions towards expecting a familiar script increased the size of the script familiarity effect in the N1 offset (162-212 ms).

In addition to task manipulations, non-sentential semantic contexts, leading to predictions for specific words or categories of words, have also been used to investigate predictive processing. Segalowitz and Zheng (2009) reported an interaction between stimulus type (word vs. pseudoword) and expectation (one vs. five categories) in the N1 (158-178 ms), wherein expectation affected N1 amplitudes for words but not for pseudowords. Their finding suggested that the N1 was sensitive to the greater predictive strength of a single semantic category. Using a similar paradigm, Hauk et al. (2012) compared ERPs in lexical and semantic decision tasks, showing that effects of category relevance were observed in the semantic decision task as early as 166 ms (data were analysed continuously, with no N1 window definition). This finding suggests, consistent with the findings of Segalowitz and Zheng, an early sensitivity to category relevance during the N1 which, given the N1’s robust sensitivity to orthography, is likely to reflect an influence of semantic-level predictions on orthographic processing.

In another attempt to modulate top-down expectancy without linguistic context, Dikker and Pylkkänen (2011) implemented a picture-noun phrase verification task. An image of a target object alone or an image of objects related to the target object was followed by a written noun phrase (article + noun) denoting the target object. They manipulated congruency and predictability. For congruent trials, the noun phrase referred to a food/drink or animal (e.g., *the apple* or *the monkey*) that matched the prior image of the object presented on its own or ‘contained’ in a stylized image (e.g., a grocery bag or Noah’s Ark, respectively). In the incongruent condition, the noun phrase did not match the prior image (single object or collection of objects). Predictability was considered high when the target object appeared on its own, and was considered low when the target object could be inferred to exist within the stylized images. Example conditions for the noun phrase, *the apple*, are determined by its preceding image as follows: an apple (congruent, high predictability), a banana (incongruent, high predictability), a bag of groceries (congruent, low predictability), or Noah’s Ark (incongruent, low predictability). Noun phrases (40 food/drink, 40 animal) were repeated four times across conditions.

Although Dikker and Pylkkanen did not examine effects in the MEG equivalent of an N1 window, they did find effects of congruency only in the high predictive condition (i.e., the apple preceded by an apple vs. a banana image) in temporal windows preceding (*∼*100 ms) and succeeding (250-400 ms) the N1. Their stimuli were designed to minimise orthographic similarity between congruent and incongruent pairs of noun phrases (i.e., maximising the magnitude of orthographic errors), suggesting that the authors anticipated that any early sensory effect of predictability may be related to orthographic processing. With only 7 participants, the study likely lacked the sample size necessary to identify such an effect in an N1-like window. In a study using the same stimuli as Dikker and Pylkkänen, Cheimariou et al. (2019) examined the effects of aging on lexical prediction indexed by the N400 component. Cheimariou et al. also did not analyse an N1 window, and used Mastoid reference sites, but did show that an early, though wide, window from 125-348 ms showed topographically broad prediction effects in younger adults, with more negative amplitudes for predictive content.

We note that related paradigms using fMRI often show orthography-semantics interactions in the likely N1 generator, the left vOT (e.g., Branzi et al., 2022; Kherif et al., 2011; J. Wang et al., 2019). However, fMRI prevents the interpretation of the timing of such effects - its coarse temporal resolution means that mapping of semantic content to representations in vOT could occur so late after word presentation as to be irrelevant to initial orthographic word recognition processes.

One advantage of paradigms like picture-word verification tasks is that the researcher can control and manipulate variables like predictability and specificity of the picture-word relation. This was demonstrated in the design used by Dikker and Pylkkänen (2011), where the picture preceding the target word unambiguously biased participants’ expectations to a single word form (with an image of one clearly identifiable object), or instead biased a set of semantically related possible word forms (with an image inducing multiple object candidates). Such a manipulation is comparable to the use of Cloze probability in sentential contexts or single versus multiple category priming, and similarly aligns with the concept of error precision or certainty.

### The Present Study

In the present study, we adapted the picture-word verification paradigm to examine the Congruency-Predictability interaction in the N1. We presented participants with PICTURE-word pairs that were congruent (e.g., ONION-onion) or incongruent (e.g., ONION-torch). Predictability of the congruent word, given the picture that precedes it, was operationalised via a continuous variable drawn from picture naming norms (Brodeur et al., 2014) reflecting the probability of the congruent word being given as a name for the picture (**Figure 2**). Picture-congruent words with very low predictability were always semantically appropriate names for their associated image, though they were difficult to predict, often because several acceptable names exist. For example, the image for *spear* in **Figure 2** could also be plausibly named with words like *lance*, *javelin*, or *pole*. Incongruent words, meanwhile, were specifically selected to be semantically incongruent with congruent words, but matched on relevant psycholinguistic dimensions. By manipulating both Congruency and Predictability of word forms, we were able to examine whether the effect of Congruency on the N1 (sensitivity to prediction error) is contingent on Predictability (certainty or precision of prediction errors), in the manner expected according to a simple predictive coding account of the N1 in which observed N1 magnitude indexes prediction error.

**Figure 2.**
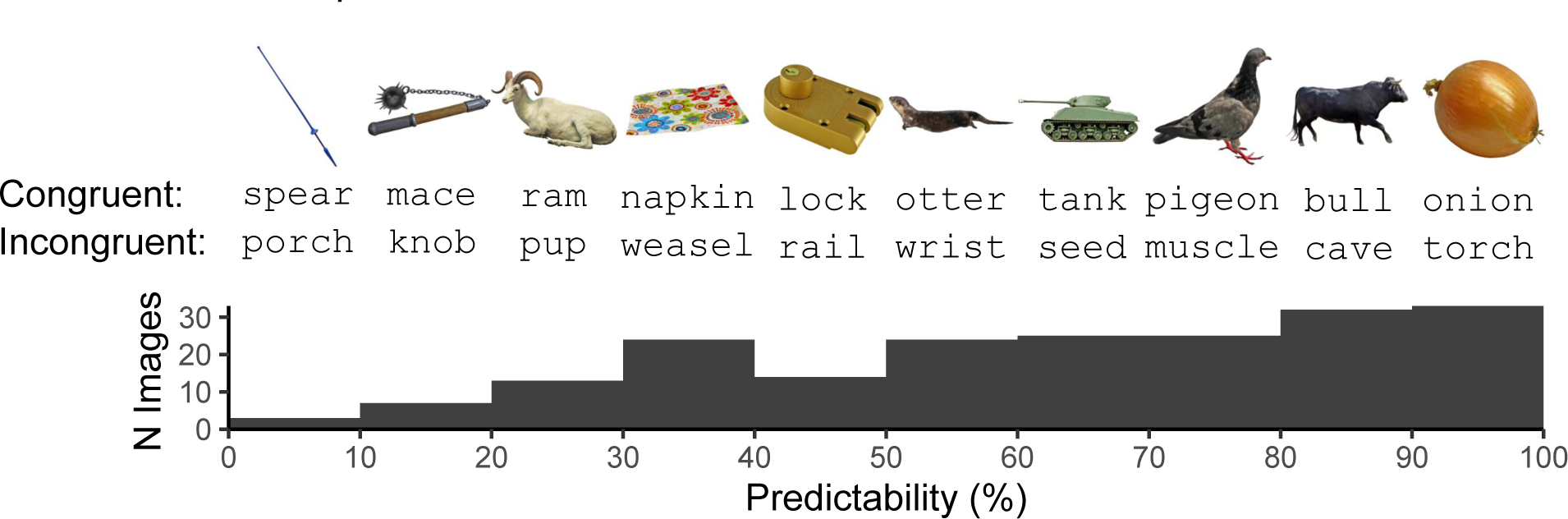
*Illustration of the experimental stimuli*. PICTURE-word pairs were either congruent (e.g., NAPKIN-*napkin*) or incongruent (e.g., NAPKIN-*weasel*), while predictability of congruent picture-word pairs varied continuously. Ten example picture-congruent and -incongruent pairs are presented, with their predictability corresponding to the histogram bin they appear above.

We hypothesised, consistent with such a predictive coding account, that that there would be a Congruency-Predictability interaction in which at the highest levels of Predictability, N1s elicited by picture-incongruent words would be more negative-going than those elicited by picture-congruent words, while at the lowest level of Predictability picture-congruent and -incongruent words should elicit N1s of similar magnitude. We anticipated three patterns of results that would have been consistent with this hypothesis:

1. higher levels of Predictability lead to a reduction in N1 magnitude only for picture-congruent words, with no such effect for picture-incongruent words (**Figure 3a**);
2. higher levels of Predictability lead to an increase in N1 magnitude only for picture-incongruent words, with no such effect for picture-congruent words (**Figure 3b**); or
3. higher levels of Predictability lead to both a reduction in N1 magnitude for picture-congruent words and an increase in N1 magnitude for picture-incongruent words (**Figure 3c**).

**Figure 3.**
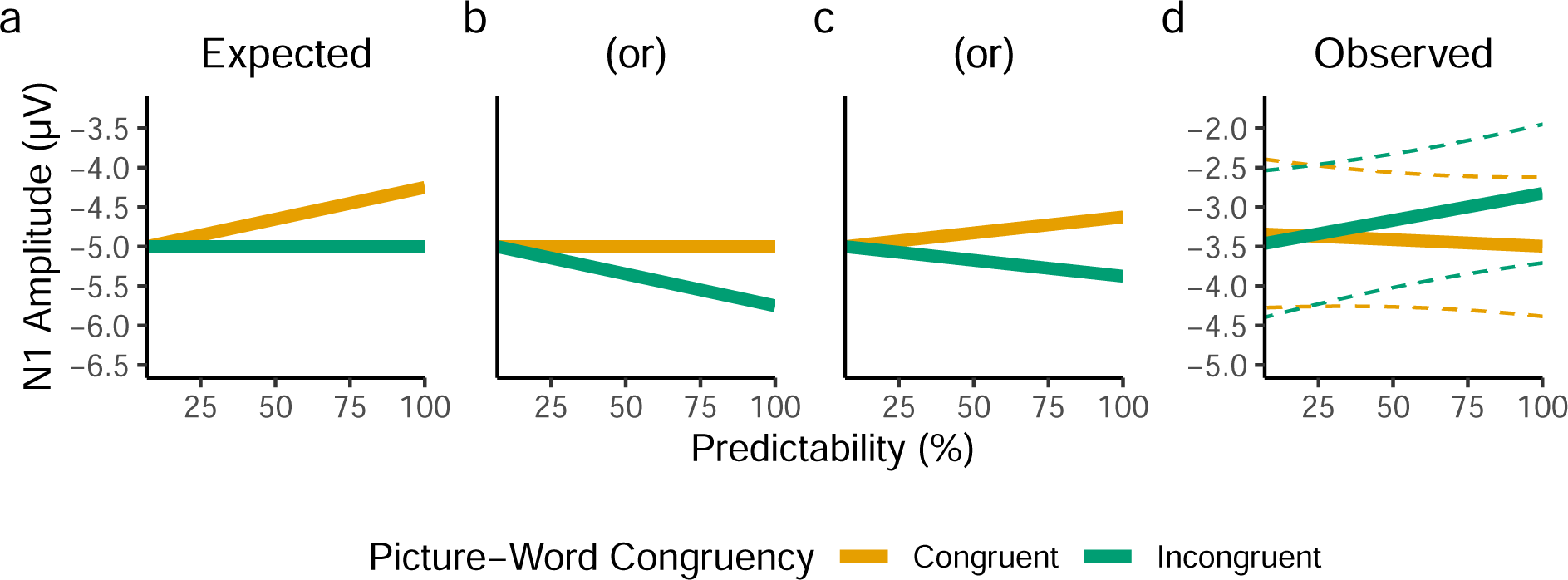
*A comparison between the predicted (**a**,**b**,**c**) and observed (**d**) patterns of results*. The predicted pattern of results was based on a predictive coding interpretation of the N1, according to which the magnitude of the N1 should be smaller for picture-congruent words relative to picture-incongruent words, and to a greater extent as Predictability increases. The observed pattern of results depicts the fixed effect predictions from the pre-registered linear mixed-effects model, with dashed lines depicting 95% bootstrapped prediction intervals (estimated from 5,000 bootstrap samples).

In our power analysis, we focused on the first of these possible patterns of results, but importantly, the Congruency-Predictability interaction term that we pre-registered to test our hypothesis would capture any of these patterns, as the interaction term’s coefficient would be in the same direction in all cases.

In our analysis, we found a pattern of effects counter to our pre-registered hypothesis (**Figure 3d**), with a Congruency-Predictability interaction in the opposite direction. An exploratory Bayesian analysis revealed that the observed interaction was 16.61 times more likely than our hypothesis. Based on these findings, we argue our results suggest that such a simplistic predictive coding account is, at least on its own, insufficient to explain the pattern of prediction effects observed in the N1 during a picture-word verification task.

This study was pre-registered at https://osf.io/jk3r4 and the reported methodology and planned analysis conform to that specified in the pre-registration, except for two changes: an accidental change to timing of stimuli, and a lowering of the EEG high-pass filter cut-off. We explain these changes in the relevant sections, and demonstrate in **Supplementary Materials F** that the change to the high-pass filter cut-off had minimal effect on the results and conclusions. All data and code are available at https://osf.io/389ce/.

## Method

The experiment included two separate tasks: The principal picture-word task was preceded by a localiser task to account for between-participant variability in the N1’s timing and location. The details of stimulus selection and control as well as presentation timing are provided in the following sections. For clarity, we first introduce the overall Congruency-Predictability design of the picture-word task. In this task, pictures of single objects are presented, followed by a noun, and participants decide whether the noun corresponds to the object. The level of Predictability of the noun was determined from norms of possible terms used to label a set of individual pictures (Brodeur et al., 2014). The most frequent, modal name agreement varied across pictures. Thus, level of noun Predictability was continuous and varied between 7% and 100%. The Congruency of the noun was either congruent (matching the modal name of the picture) or incongruent (a semantically unrelated noun matched across several lexical variables).

### Materials: Picture-Word Task

A total of 400 words (200 per Congruency condition) were selected with LexOPS (Taylor et al., 2020), a package for the generation and control of lexical variables in the R programming language (R Core Team, 2021). Picture-congruent and -incongruent words were matched precisely in an item-wise manner on a range of relevant psycholinguistic variables, comprising word length, frequency, concreteness, OLD20, and character bigram probability. To ensure that picture-incongruent words were not inadvertent possible descriptors for images, we minimised the semantic relatedness between pairs of words. Additionally, counterbalanced sets of stimuli were matched on distributions in these variables using a measure of distributional similarity (Pastore & Calcagnì, 2019). A full description of the method by which stimuli were selected, and a full list of stimuli, is available in **Supplementary Materials A**.

Before embarking on the electrophysiological picture-word experiment, we first ran a proof-of-concept behavioural experiment using a different stimulus set generated from a very similar pipeline. We anticipated that increased Predictability should cause faster response time (RT) for congruent trials and have either no effect or a minimal effect on performance for incongruent trials. The results from this behavioural validation are presented in **Supplementary Materials B**. In short, we observed the pattern of results consistent with our expectations, with Predictability leading to faster RTs for congruent trials, but having almost no effect on incongruent trials.

### Materials: Localiser Task

The precise location of the N1, and timing of its peak amplitude, is known to vary across studies and among participants. As such, we did not specify a common N1 electrode or timepoint shared among all participants before data collection. Instead, we employed a localiser task to identify, within an appropriate region and time period of interest, the electrode and timepoint at which each participant’s maximal sensitivity to orthography emerges (i.e., more extreme amplitudes for words than false-font stimuli).

This data could then be used to extract N1 amplitudes in the picture-word task, while accounting for variability among participants in timing and topography of orthographic processes.

For the localiser task, three categories of stimuli were presented for 100 trials each (**Figure 4**). These consisted of matched triplets of words (Courier New font), false-font strings (BACS2serif font; Vidal et al., 2017), and phase-shuffled words. The comparison between words and false-font strings is a standard measure of N1 sensitivity to orthography, with previous evidence suggesting a more robust difference than exists between nonwords and words (Brem et al., 2018; Maurer, Brandeis, & McCandliss, 2005; Pleisch et al., 2019). However, phase-shuffled words were employed as an alternative comparison for exploratory analyses, with equal spatial-frequency amplitude and permuted spatial-frequency phase. Similar phase-shuffled word stimuli have shown robust differences to word forms in fMRI investigations of vOT activity (Rauschecker et al., 2012; Rodrigues et al., 2019; White et al., 2019; Yeatman et al., 2013).

**Figure 4.**
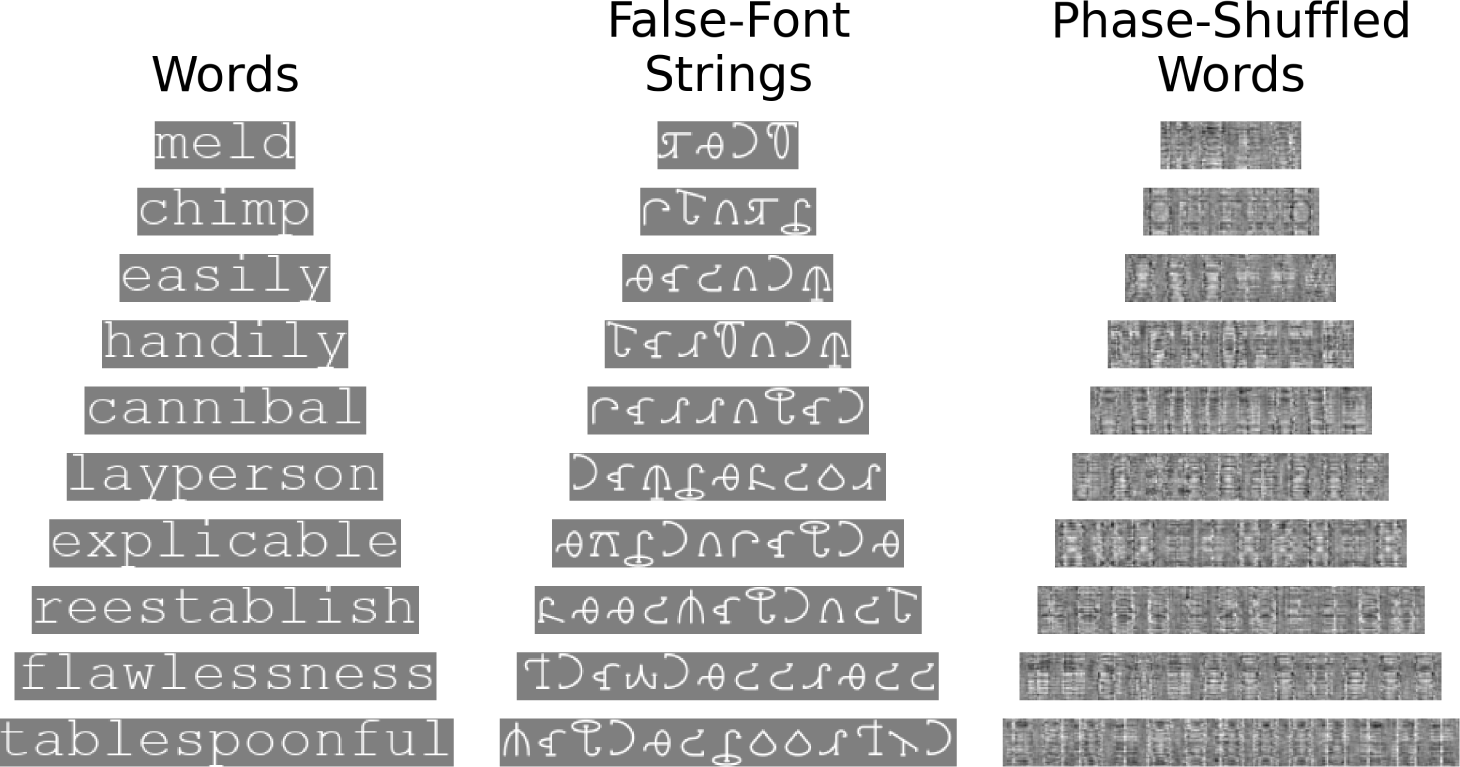
*Ten example stimuli for each stimulus type in the localiser task*. Each row represents a matched triplet of word, false-font string, and phase-shuffled word stimuli. The phase-shuffled word images were generated uniquely for each trial.

The word stimuli were selected to be widely known by participants (>90% proportion known), and to be representative on a range of psycholinguistic variables including length, frequency, part of speech, and prevalence. A full description of how the Localiser Stimuli were selected, and a list of all word stimuli, is presented in Supplementary Materials C.

### Participants

The sample size of 68 participants was decided via a power analysis using Monte-Carlo simulations of a realistic effect size (**Supplementary Materials D**). This revealed that with *≥*68 participants we could expect >80% statistical power in the long run (**Figure 5**). All 68 participants (40 female, 27 male, 1 non-binary) were monolingual native English speakers. Participants were randomly allocated into one of the four combinations of stimulus set (Set 1, Set 2) and response group (i.e., the left-right mapping of the two response buttons for affirmative and negative responses), such that each combination of stimulus set and response group comprised 17 participants. No participants reported diagnosis of any reading disorder. Ages ranged from 18 to 37 years (M=22.69, SD=4.9), and all participants reported having normal or corrected-to-normal vision. Participants’ handedness was assessed via the revised short form of the Edinburgh Handedness Inventory (Veale, 2014), with participants only permitted to take part if they scored a laterality quotient of +40 indicating right handedness. Exclusion criteria for participants were determined prior to data collection as follows: (1) if 10 or more channels showed an offset more extreme than ±25 mV (as measured on the BioSemi acquisition software, ActiView), or (2) if more than 5% of the trials were lost due to technical issues with the EEG system. As no participants satisfied these criteria, no participants were excluded after data collection. Data collection was approved by the Ethics Committee of the University of Glasgow College of Science and Engineering (application number: 300200117), and all participants provided informed consent.

**Figure 5.**
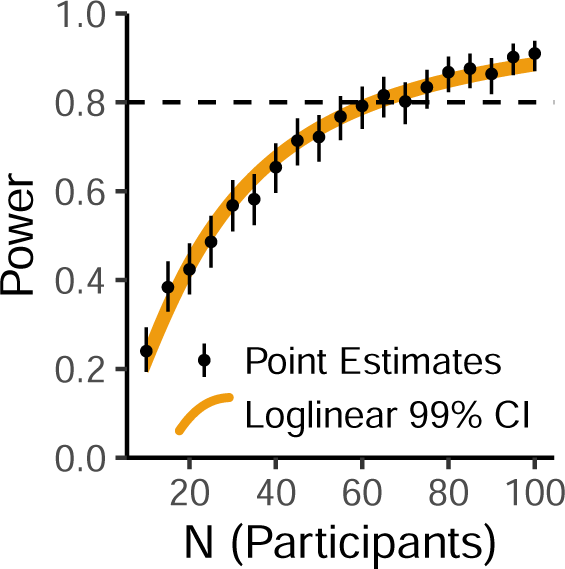
*Estimated relationship between number of participants and statistical power*. *Black* points and error bars depict point estimates ±99% Binomial confidence intervals, each from 500 simulations. As 500 simulations provides a noisy estimate, we interpolated the relationship between N and power via a loglinear, logit-link Binomial model. The *orange* region depicts the 99% confidence intervals of this loglinear model.

### Procedure

Stimuli were presented on a VPixx Technologies VIEWPixx screen (resolution 1920×1080 pixels, diagonal length 23”, model VPX-VPX-2004A). Participants completed the experiment on a chin rest positioned 48 cm from the centre of the screen. Stimuli were presented on a grey background equal to 50% of the maximum intensity in each colour channel, roughly 12.3 cd/m^2^. The experiment was written using the Python library PsychoPy (Peirce, 2007), and all code and materials are available in the repository associated with the study. All stimuli were presented centrally (horizontally and vertically). All trials in both tasks were presented in a pseudo-randomised order, such that no more than five consecutive trials required the same response from the participant. Trials were randomised across blocks, with the exception of the practice block, for which trials were randomised within the one block.

A mistake in the lab setup, which we discovered after data collection, meant that the display screen was running at 120 Hz rather than an expected 60 Hz. As we were controlling stimulus presentation by screen refreshes, this meant that all our stimuli were presented for half the expected durations. For this reason, the veridical stimulus durations described here differ from those described in the pre-registration.

Participants started with the localiser task, in the form of a lexical decision task (**Figure 6a**). The localiser task began with 30 practice trials, and was then followed by 300 trials split into 5 blocks of 60 trials. Each trial began with the bullseye fixation target recommended by Thaler et al. (2013) (outer and inner circle diameters were 0.6° and 0.2° of visual angle), presented for 150 ms. This was followed by a jittered interval of between 150 and 650 ms, during which the screen was blank. The stimulus (word, false-font string, or phase-shuffled word image) was then presented at a height of 1.5° (width of 1.07° for one character). Words and false-font strings were presented in white (80 cd/m^2^), in the respective fonts of non-proportional Courier New and BACS2serif font. The stimulus was visible for 250 ms, after which the font colour changed to green to signal participants to respond. Participants were requested to respond once after the stimulus changed colour, quickly and accurately, to indicate whether the stimulus they saw in each trial was either a word or not a word. The stimulus remained on screen until the participant responded. Responses were given with the right and left control (‘Ctrl’) keys of a QWERTY keyboard, with the mapping of affirmative and negative responses counterbalanced across participants. After the participant had responded, there was a delay of around 100 ms (variable as data was saved to disk during this interval), and then the next trial began.

**Figure 6.**
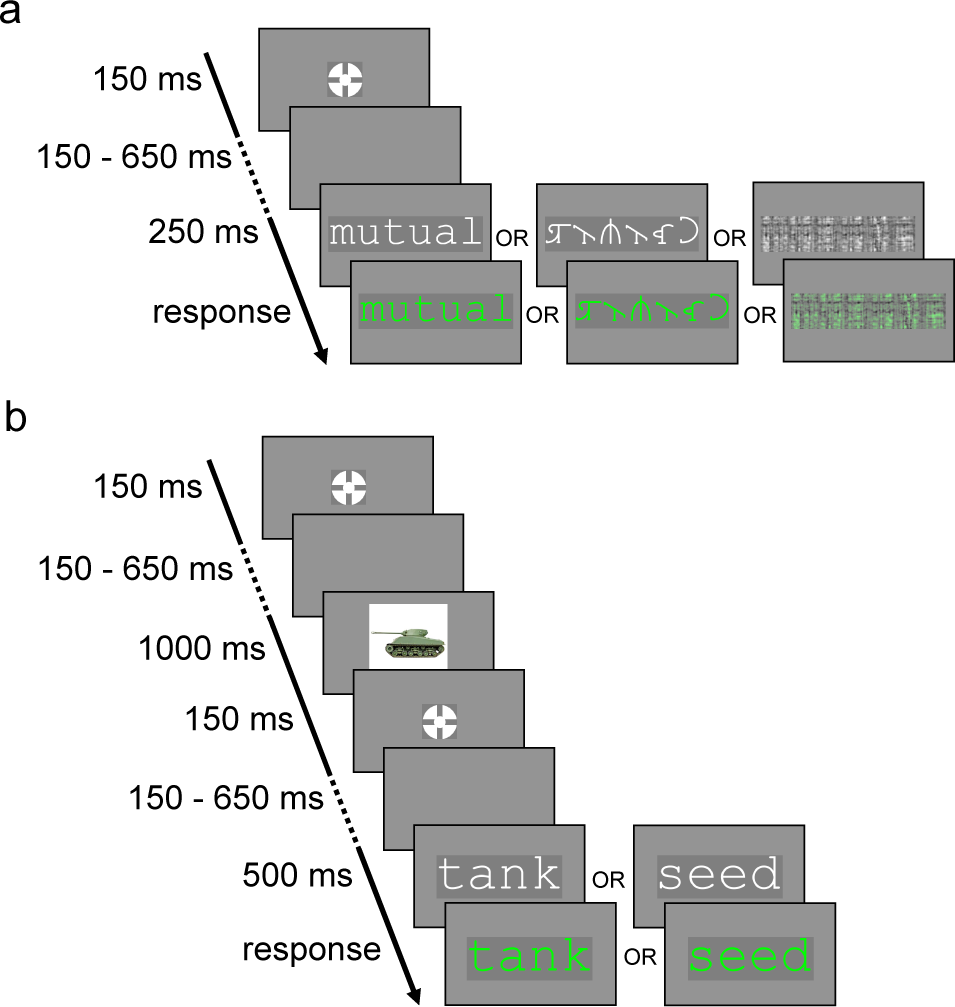
*Trial structure of the (**a**) localiser task and (**b**) picture-word task*. This figure is illustrative and the sizes are not to scale; in the experiment, images were in fact presented at a much larger scale than words.

After the localiser task, participants completed the picture-word task (**Figure 6b**), comprising an initial practice block of 20 trials, followed by 200 trials split into 5 blocks of 40 trials. As in the localiser task, each trial in the picture-word task began with the bullseye fixation point, presented for 150 ms, after which there was a blank screen for a jittered interval of between 150 and 650 ms. An image was then presented for 1000 ms, at a size of 10x10°. The bullseye fixation point was then presented again for 150 ms, followed by another interval jittered between 150 and 650 ms. The word was then presented in white Courier New font, at a height of 1.5° (width 1.07° for one character). After 500 ms, the word turned green, and participants could provide their response to indicate whether the word described the image they saw. The word remained on screen until the participant responded. As in the localiser task, responses were given with the right and left control (‘Ctrl’) keys of a QWERTY keyboard, with the mapping of affirmative and negative responses counterbalanced across participants, but kept consistent within participants across the two tasks. After participants had responded, there was a delay of around 100 ms (again, variable as data was saved to disk during this interval), and then the next trial began. There was no deadline for participants to respond. The instructions given to participants for the picture-word task are presented in **Supplementary Materials E**.

The first blocks of both tasks consisted of practice trials with 10 exemplars for each stimulus type (word or false-font string or phase-shifted image, and congruent or incongruent noun for the localiser and picture-word tasks, respectively), during which participants were additionally given immediate feedback on their accuracy for each trial. These practice trials were followed by green text reading ”CORRECT!” if the participant responded correctly, or else by red text reading ”INCORRECT!”, presented in Courier New font with a height of 1.5°, for 1000 ms. Participants had self-paced breaks between blocks for each task. Before the practice trials and at the start of every experimental block, participants were presented with instructions for the task (available in **Supplementary Materials E**), summarising what would occur in each trial, and specifying that they should respond as quickly and accurately as possible once the stimulus turned green. These instructions also specified which keys participants should press to indicate their decision. After each experimental block, including the practice trials, participants were presented with their average accuracy and median response time. After the practice trials, participants were additionally given the option to run the practice trials again. In the experimental blocks, no trial-level feedback was provided.

### Recording

EEG data were recorded using a 64-channel BioSemi ActiveTwo system, sampling at 512 Hz, with an online anti-aliasing low-pass filter cutoff at one fifth of the sample rate (i.e., 102.4 Hz). Electrodes were positioned in the standard 10-20 system locations. Four electro-oculography (EOG) electrodes were placed to record eye movements and blinks: 2 were placed to the sides of eyes (on the right and left outer canthi), and 2 below the eyes (on the infraorbital foramen). Electrode offset was kept stable and low through the recording, within ±25 mV, as measured by the BioSemi ActiView EEG acquisition tool. Electrodes whose activity exceeded this threshold were recorded but were removed (and interpolated) in data preprocessing.

### Preprocessing

The following section details the procedure applied to EEG data from each individual session, with the same pipeline being applied to both the localisation task and picture-word task unless otherwise specified. EEG preprocessing was achieved using functions from the EEGLAB (Delorme & Makeig, 2004) toolbox for MATLAB (MATLAB, 2022) or OCTAVE (Eaton et al., 2020). For both tasks, trials were excluded if responded to incorrectly (total of *N_trials_*=368, or .02%, in localiser task, and *N_trials_*=226, or .02%, in picture-word). Further trials were excluded if responded to later than 1500 ms after the word (or nonword) changed colour (total of *N_trials_*=41, or .002%, in localiser task, *N_trials_*=42, or .003%, in picture-word).

Channels recorded as having offsets ±25 mV during data acquisition were removed from the data (in both tasks, 56 channels, or 1.27%, were removed across all participants), with their activity to be later interpolated. In addition, we found that even when not identified as problematic during recording, the channel PO4 was consistently noisy, and so we interpolated this channel for all participants. PO4 was not part of our left occipitotemporal region of interest, but was interpolated for exploratory analyses of the whole scalp, and to avoid affecting other preprocessing steps. Interpolating electrode PO4 was not a preregistered step. However, we note that this change did not alter the direction of any results, rather, only reducing the size of effects. After interpolation, the EEG data were then re-referenced to the average activity across all electrodes and filtered with a 4th order Butterworth filter between .1 and 40 Hz. To counteract the distortion in signals’ timing (phase) that is inherent to causal filters, the filter was applied in both directions (i.e., two-pass), with the MATLAB function *filtfilt()*. In our pre-registration, we specified that we would apply a Butterworth filter with a bandpass of .5-40 Hz. However, after the pre-registration, we considered that, consistent with research into the effects of high-pass filters (Rousselet, 2012; Tanner et al., 2015; VanRullen, 2011), this could produce artefactually early effects. As a result, we lowered the high-pass filter to a less problematic .1 Hz. For comparison, demonstrating that our change to the pre-registered pipeline had minimal effect on the results or our conclusions, the results using the original filter are presented in **Supplementary Materials F**.

Segments of data outside of experimental blocks (i.e., in break periods) were identified and removed so they did not impact the independent components analysis (ICA) applied later in the pipeline. Blocks were identified as beginning 500 ms before stimulus presentation in the first trial of each block, ending 500 ms after the end of the last trial’s epoch. To reduce the impact of occasional non-stationary artefacts with high amplitude (such as infrequent muscle movements), artefact subspace reconstruction (ASR; Chang et al., 2020) was used with a standard deviation cutoff of 20 to remove non-stationary artefacts. Following this, an ICA was run on the data to identify more stationary artefacts. The ICA was run using the FastICA algorithm (Hyvärinen & Oja, 1997), with a recorded random seed for reproducibility. The ICA was run on a copy of the data with channel offsets removed to allow for better sensitivity to electro-oculogram (EOG) artefacts (Groppe et al., 2009). The ICLabel classifier (Pion-Tonachini et al., 2019) was used to automatically identify artefacts which were eye- or muscle-related.

Components classified by ICLabel as eye-related or muscle-related with a probability of *≥*85% were removed from the data. Following eye movement artefact removal, activity from channels which were removed was interpolated via spherical splines (Localiser: *M* =1.14 per participant, *SD*=1.58; Picture-Word: *M* =1.68, *SD*=2.03), as implemented in EEGLAB. Trials were then epoched and baseline-corrected to the 200 ms preceding stimulus presentation. For the localiser task, stimulus presentation refers to the time point at which words, false-font strings, or phase-shuffled images were presented; in the picture-word task, stimulus presentation refers to the target word.

For the planned analysis, we pre-registered an approach to maximise sensitivity to effects of Congruency and Predictability on the N1. To encompass the typical topography and timing of the posterior left-lateralised N1, we selected eight occipitotemporal electrodes (**Figure 7**; electrodes O1, PO3, PO7, P5, P7, P9, CP5, and TP7) and a 120-200 ms window. In contrast to some previous studies whose N1 windows extended beyond 200 ms, we set 200 ms as an upper bound for the possible maximal timepoint in the main analysis, to ensure effects were indeed restricted to the N1, and not later components like the N400. For each participant, we identified the electrode that showed maximal sensitivity to orthographic information in the N1 during the localisation task. Specifically, each participant’s “maximal electrode” (within the region of interest and selected time window) was the one which showed the largest mean amplitude difference, in the expected direction, across all localiser trials between word and false-font string stimuli. The expected direction was a more negative-going N1 for words than for false-font strings, a pattern based on previous findings (Appelbaum et al., 2009; Bentin et al., 1999; Eberhard-Moscicka et al., 2016; Pleisch et al., 2019; Zhao et al., 2014). Each participant’s “maximal timepoint” was the timepoint at which the maximal electrode showed the greatest sensitivity to the word-versus-false-font difference in the expected direction. Each participant’s maximal electrode and maximal timepoint were then used to extract their trial-level N1 amplitudes from the picture-word task. To reduce the influence of noise on trial-level data, the trial-level N1 amplitudes in the picture-word task were calculated as the maximal electrode’s mean amplitude across 3 timepoints: the participant’s maximal timepoint, and the timepoints immediately preceding and following it. At the recorded sample rate of 512 Hz, this is equivalent to a window of 5.85 ms (i.e., 1/512*3) centred on the maximal timepoint.

**Figure 7.**
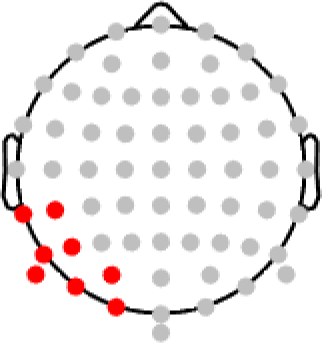
*The left-lateralised occipitotemporal region of interest selected for the N1 (highlighted in red)*.

### Planned Analysis

Our planned analysis tested the pre-registered hypothesis of a Congruency-Predictability interaction in which N1 amplitudes are reduced (i.e., less negative going) for picture-congruent trials than for picture-incongruent trials, and in which this difference is greatest at the highest levels of predictability, and smallest at the lowest levels of predictability. This was based on the notion that the N1 indexes prediction error in biasing contexts.

The trial-level N1 amplitudes from the picture-word task were modelled using a linear mixed-effects model fit with the R package *lme4* (Bates et al., 2015), estimating the maximal random effects structure justified by the experiment’s design (Barr et al., 2013) as detailed in the section on the power analysis. The model was fit using the *bobyqa* optimiser (Powell, 2009). In *lme4* syntax, the formula for the mixed-effect model was specified as:

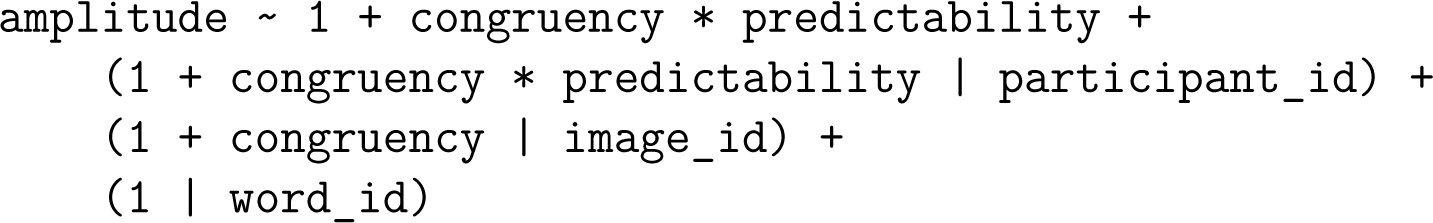

In this formula, *amplitude* is the trial-level N1 amplitude in microvolts, while *congruency* is a deviation-coded categorical variable indicating whether a given trial’s word was picture-congruent or -incongruent, and *predictability* refers to the proportion of name agreement in the BOSS norms, normalised between 0 and 1. A consequence of this coding method is that the model’s intercept reflects the predicted amplitude at the lowest level of Predictability, averaged across both levels of Congruency, while the slopes’ coefficients are standardised and directly comparable in their magnitude. The variables of *participant_id*, *image_id*, and *word_id*, in the formula, identify each trial’s participant, image, and word, respectively.

## Results

The planned, pre-registered analysis examined whether the hypothesised effect of a Predictability-dependent reduction of N1 amplitudes for picture-congruent words was observed at the electrode/timepoint in which each participant showed maximal sensitivity to orthography. We then present exploratory analyses, which respectively examine the Bayesian probability that our data are consistent with the hypothesis, and delineate the time-course of the Congruency-Predictability interaction. We also conducted further exploratory analyses, which we report in the supplementary materials, examining behavioural results in the picture-word study (**Supplementary Materials G**), and EEG and behavioural results from the localiser task (**Supplementary Materials H**)

### Planned Analysis

The fixed effect relationships estimated in the planned analysis are presented in **Figure 8**. The model intercept, reflecting the average N1 amplitude at the lowest level of Predictability, was estimated to be *β*=-3.4 µV (*SE*=.48). The fixed effect of Congruency from this model was estimated as *β*=-.12 µV (*SE*=.34), which captures that, at the lowest level of Predictability (7%), N1 components for picture-congruent and -incongruent words were estimated to be quite similar (.12 µV difference). The main effect of Predictability was estimated as *β*=.25 µV (*SE*=.27), meaning that N1 amplitudes, averaged across congruent and incongruent conditions, were .25 µV less negative-going at the highest level (100%) than at the lowest level of Predictability (7%). The effect of interest, the interaction between Congruency and Predictability, was in the opposite direction from that hypothesised, estimated as *β*=-.79 µV (*SE*=.52). As our hypothesis was directional, with a prediction in the opposite direction, we interpret these results as a failure to find evidence in favour of the hypothesis.

**Figure 8.**
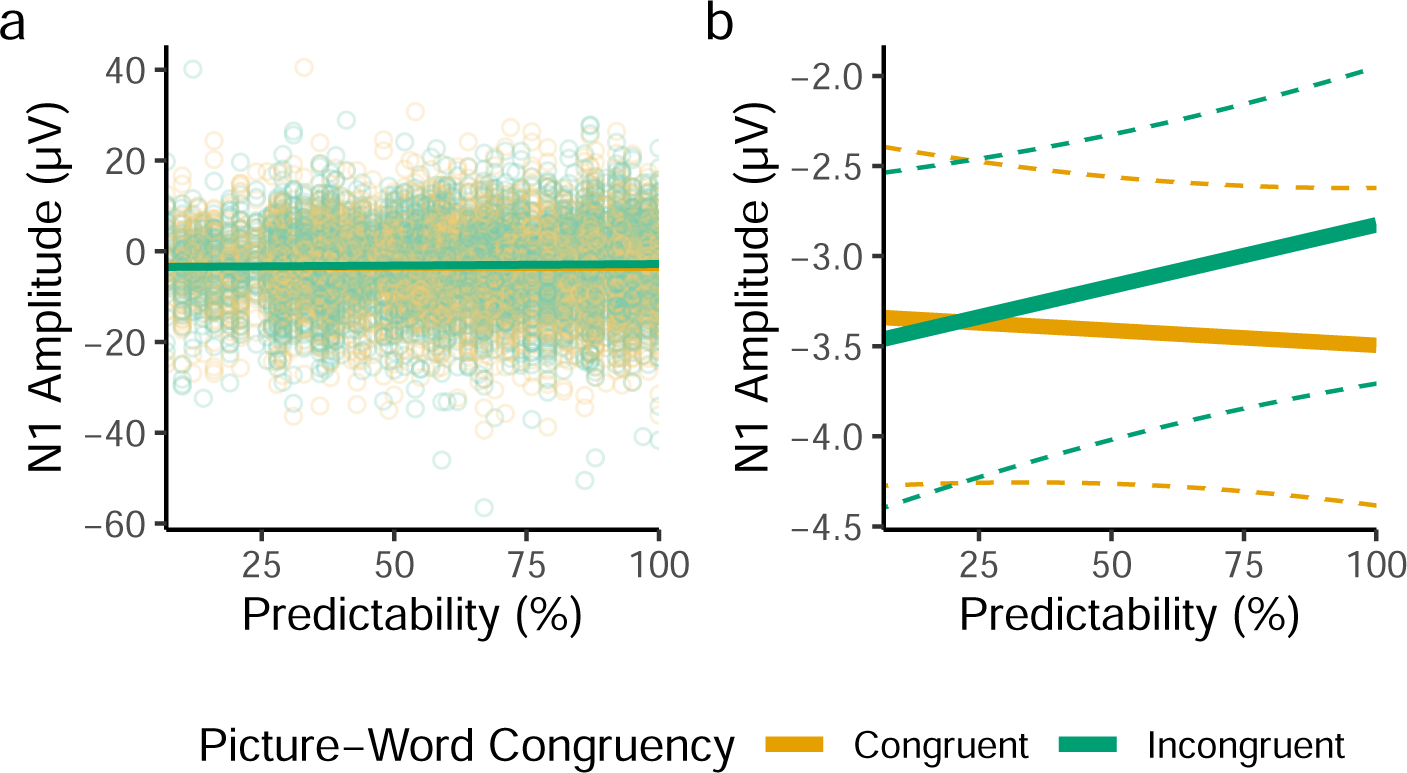
*Fixed effect predictions from the planned analysis of the picture-word task*. (**a**) Model-derived fixed-effect predictions, visualised over results from all trials (individual points). (**b**) Fixed-effect predictions visualised alone for visibility, with dashed lines depicting the bounds of 95% bootstrapped prediction intervals (estimated from 5,000 samples), where bootstrapped predictions were generated using the *bootMer()* function of *lme4*. For feasibility, bootstrapped predictions were generated from a version of the model that lacked random slopes.

To describe the estimated interaction, for picture-incongruent words, the effect of Predictability was estimated to be *β*=.63 µV (*SE*=.36), while for picture-congruent words, the effect of Predictability was estimated to be *β*=-.15 µV (*SE*=.4). As such, the slopes for the effect of Predictability in both Congruency conditions were of different magnitudes, and were both in directions inconsistent with our predictive coding hypothesis.

For comparison, we also analysed the data altering aspects of our planned analysis method: first using the maximal electrodes that would be identified from the comparison between words and phase-shuffled words, and second using averages within the occipitotemporal region of interest (**Supplementary Materials I**). These exploratory analyses revealed very similar patterns of effects, with estimates of the Congruency-Predictability interaction similarly inconsistent with our hypothesis, which we derived from a simple predictive coding account of the N1.

### Exploratory Bayesian Analysis

We observed a Congruency-Predictability interaction in the opposite direction (i.e., negative) to what we expected under our predictive coding hypothesis (i.e., positive). To explicitly quantify the probability of our predictive coding hypothesis, we fit a Bayesian implementation of the model described in the planned analysis, in STAN (STAN Development Team, 2023) via *brms* (Bürkner, 2017). This model was fit to the same data, and estimated the same hierarchical formula, with the same Gaussian link function as that described above, but was specified with weakly informative priors for the fixed effects. Specifically, the prior for the fixed effect intercept was specified as a normal distribution of mean -5, and *SD* 10, while all fixed effect slopes’ priors were specified as normal distributions centred on 0, with *SD*s of 5. Covariance matrices were assigned flat priors, and default priors for brms were used for random effect *SD*s and the sigma parameter of the normal distribution. The model was fit with 5 chains and 5000 iterations per chain (split equally between warmup and sampling) such that there were a total of 12,500 posterior samples. Consistent with the linear mixed-effects model we fit via *lme4*, this analysis revealed a median posterior estimate for the Congruency-Predictability interaction of *β*=-.79 µV (89% highest density interval = [-1.59, .013]; **Figure 9**). We calculated, given this posterior distribution, that the Congruency-Predictability interaction is 16.61 times more likely to be less than 0, than it is to be greater than zero (that is, *BF*_01_), which we consider to be strong evidence against our hypothesis.

**Figure 9.**
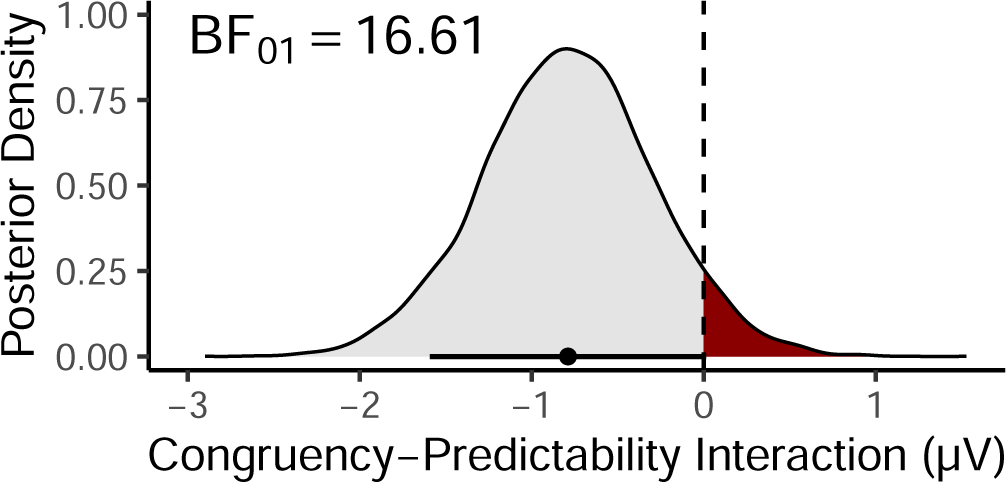
*Posterior density for the Congruency-Predictability interaction*. The region of the posterior distribution consistent with the predictive coding hypothesis (where *β*>0) is highlighted in *red*. The point and horizontal line below the density plot depict respectively the median estimate and 89% highest density interval of the posterior distribution.

We considered that our use of a localiser task may have been an inappropriate approach for identifying electrodes sensitive to orthographic information. Indeed, our pre-registered approach for identifying maximal electrodes specified the direction of the difference that should be used, with more negative N1s for words than for false-font strings. However, exploratory ERP analyses of the localiser task showed that left-lateralised occipitotemporal electrodes showed a more negative N1 peak overall for false-font strings than for words (**Supplementary Materials H**). Our approach may therefore have systematically selected electrodes that are not representative of the ROI. As a result, we re-ran the Bayesian analysis as described above, but modelling average amplitudes from all electrodes in the left occipitotemporal ROI (**Supplementary Materials I**). This revealed even stronger evidence against the hypothesis, with a Congruency-Predictability interaction for the average amplitude in the ROI of *β*=-1.03 µV (89% highest density interval = [-1.52, -.058], estimated to be 2082 times more likely to be less than 0, than it is to be greater than zero.

### Exploratory Time-Course Analysis

To examine the time-course of effects, we fit separate linear mixed-effects models to sample level data for the left-lateralised occipitotemporal region of interest, with variables coded as described for the planned analysis. For feasibility, data were down-sampled to 256 Hz, and the models did not estimate random slopes. To account for variability between electrodes, and for per-participant differences in topography, random intercepts were estimated for each combination of participant and electrode. In *lme4* syntax, the model formula was specified as follows:

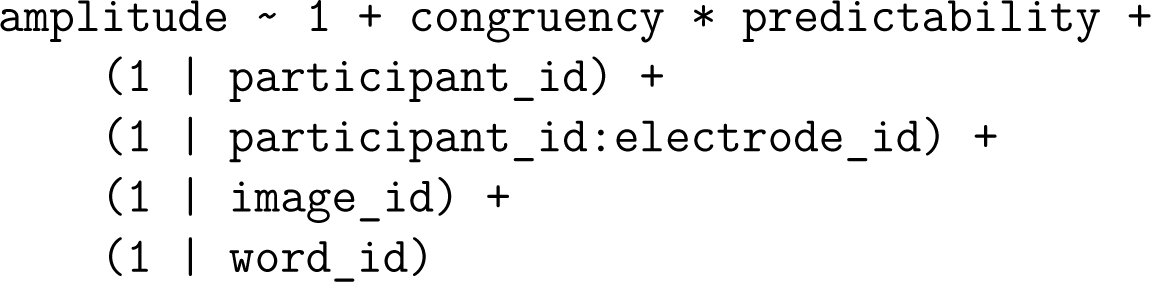

The results (**Figure 10**) reproduced findings from the planned analysis, with increases in Predictability associated with more negative (larger) N1 amplitudes for picture-congruent words, and with less negative (smaller) N1 amplitudes for picture-incongruent words. The Congruency-Predictability interaction of interest remained negative, and thus in the opposite direction to that hypothesised, throughout the N1.

**Figure 10.**
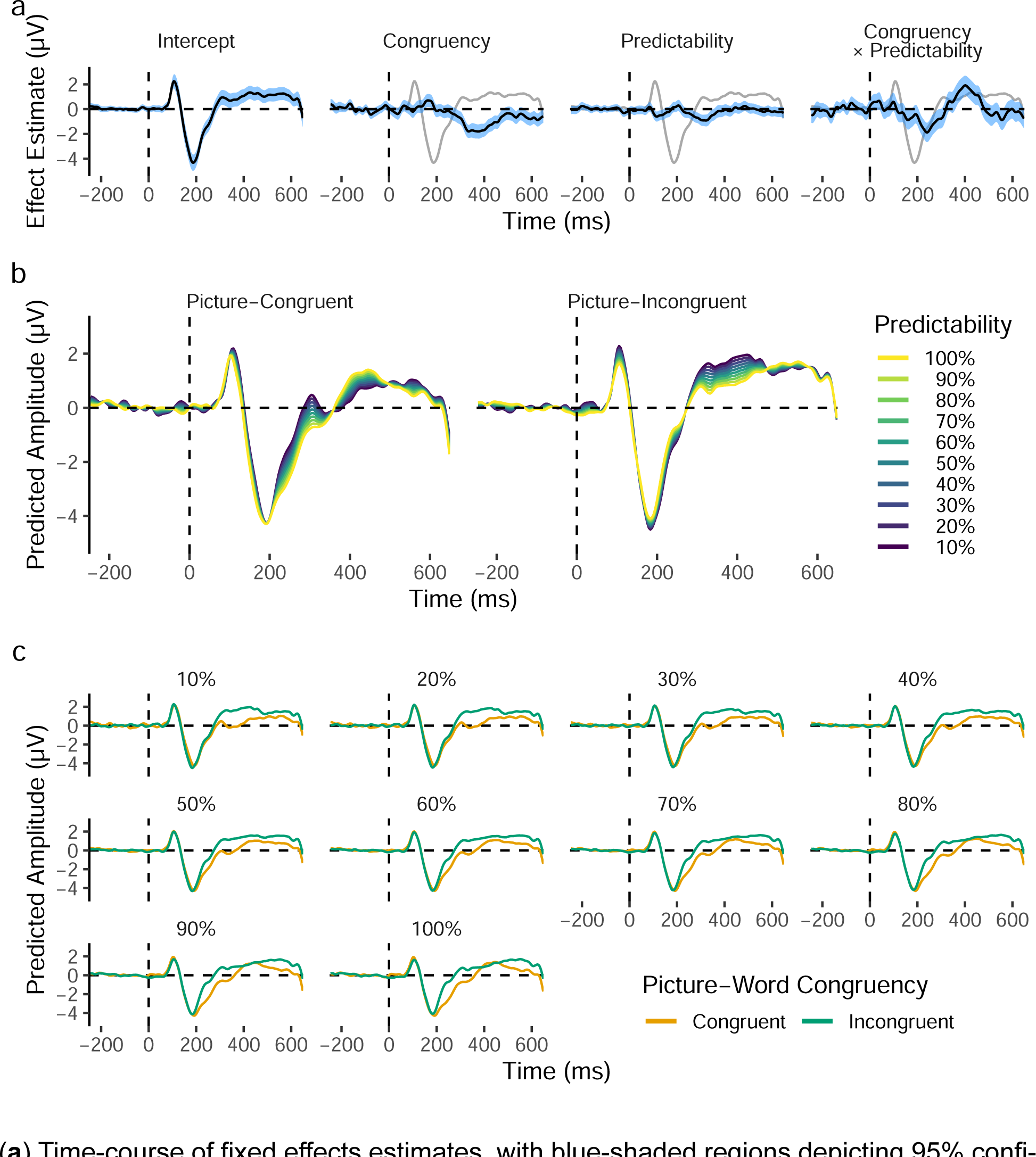
*Time-course of fixed effects from the sample-level analysis of the left-lateralised occipitotemporal region of interest*. (**a**) Time-course of fixed effects estimates, with blue-shaded regions depicting 95% confidence intervals. The model intercept (reflecting average amplitudes at the lowest level of Predictability) is depicted as a grey line on each panel to provide a reference for timing and magnitude of effects. (**b**) Fixed-effect predictions for picture-congruent and -incongruent words at levels of Predictability from 10 to 100%, in steps of 10%. (**c**) Same data as (**b**), but split by Predictability rather than Congruency. The rapid change in amplitude after 650 ms was likely elicited by the stimulus colour change at 500 ms, as shown more clearly in Figure 12.

The sample-level analysis additionally suggested that the difference was largest in the N1’s offset period (succeeding the peak). A later Congruency-Predictability interaction was also observed, peaking at around 400 ms (possibly resulting from effects in the N400 component) in the opposite direction to that observed for the N1’s offset. To better understand the time-course of the Congruency-Predictability interaction, we examined the time-course of the effect of Predictability for picture-congruent and -incongruent words separately (i.e., simple effects; **Figure 11**). This showed more clearly that Predictability reduced amplitudes in the N1 for picture-incongruent words, but increased amplitudes for picture-congruent words. This difference peaked around 225 ms, but reversed in direction after 300 ms. It is of note that the peak of the observed effects in the N1 was later than originally anticipated (the planned analysis was limited to *≤*200 ms). Nevertheless, the model intercept (**Figure 10a**) clearly shows that these effects peaked during the N1’s offset period.

**Figure 11.**
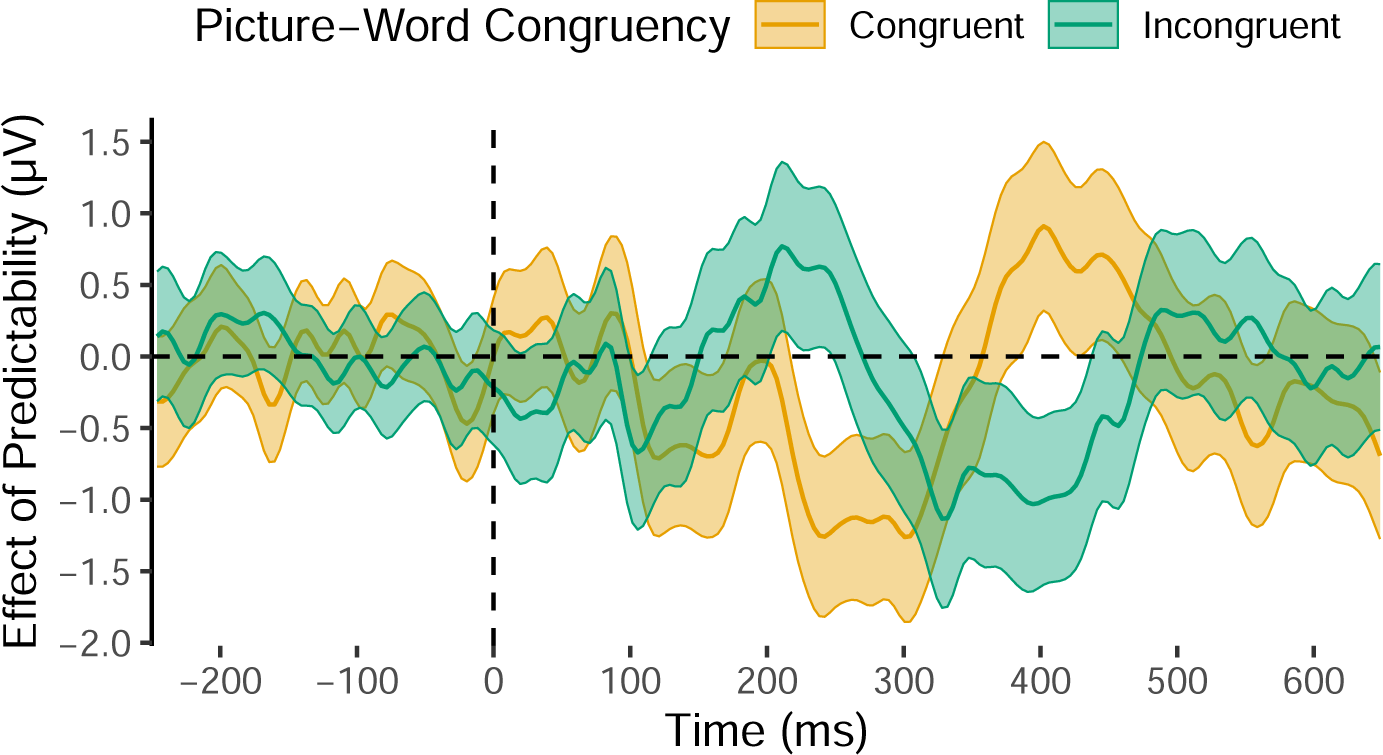
*Time-course of the effect of Predictability for picture-congruent and -incongruent words*. Central lines depict effect estimates, derived from sample-level models that were coded such that the model intercept lay at the respective levels of picture-word Congruency. Estimates reflect occipitotemporal ERPs for words at the maximum level of Predictability, minus those at the minimum level of Predictability. Shaded areas depict 95% confidence intervals of model estimates.

### Exploratory Scalp-Wide Analysis

Finally, we examined how the full topography of effects changed over time (**Figure 12**). Specifically, we fit a linear mixed-effects model to data from each time-point and electrode separately, with variables coded as described in the section on the time-course analysis of the region of interest. As in that analysis, we again excluded random slopes for feasibility:

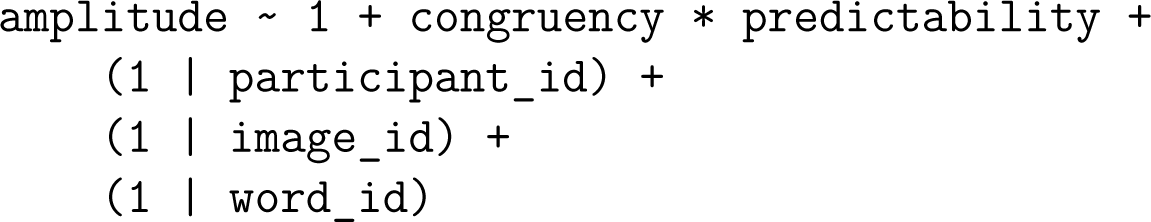

**Figure 12.**
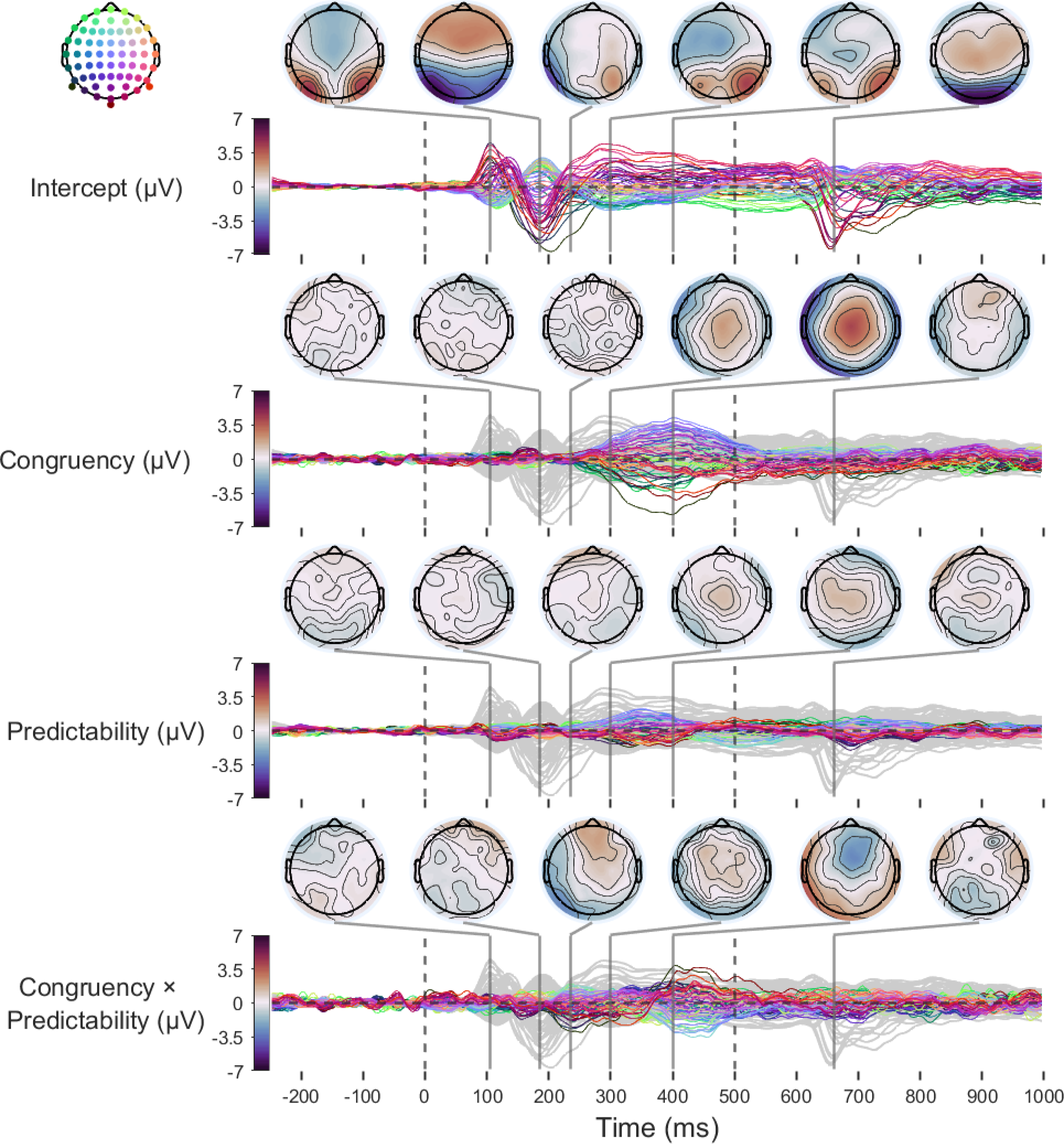
*Time-course of scalp-wide fixed-effects estimates.* The first dashed vertical line (0 ms) indicates stimulus (word) onset. The second dashed vertical line (500 ms) indicates the time-point at which the word changed colour to green. Topographic plots of fixed effects are highlighted at key time-points. Model intercepts (reflecting average amplitudes at the lowest level of Predictability) are depicted as grey lines on each panel to provide a reference for timing and magnitude of effects.

Results confirmed that the Congruency-Predictability interaction at left occipitotemporal sites was the earliest fixed effect to emerge, and that the effect was small relative to that observed at later time points. It additionally showed that the switch in direction of the Predictability-Congruency interaction shown to peak at around 400 ms in **Figure 11** exhibits a frontocentral topography. This effect was sustained until around 475 ms. Interestingly, if this effect captures changes in an N400 component, then the direction of the N400 modulation was, as was the case for the effect on the N1, arguably inconsistent with a simple predictive coding account. This is because the direction of effects suggests that prediction-congruent words elicited the most negative-going N400 amplitudes at the lowest level of predictability. As predictability increased, N400 amplitudes elicited by picture-congruent words became less extreme, increasingly approaching the N400 amplitudes elicited by picture-incongruent words (**Supplementary Materials J**). The modulation observed in the opposite direction at occipitotemporal sites in **Figure 10** at around 400 ms likely results from the use of average reference.

The scalp-wide analysis also revealed that the main effect of picture-word Congruency shown in **Figure 10** indeed peaks at around 400 ms, with a centroparietal topography. Interpreting this as a modulation of the N400 would mean that, at the lowest level of predictability, picture-congruent words elicited more negative-going N400s overall than picture-incongruent words did.

Finally, this analysis covered an extended period of time, which revealed a clear negative-going posterior component at around 650 ms. This component peaked around 150 ms after the stimulus changed colour to indicate that participants could respond.

Given the timing and topography of this component, this is likely to reflect an N1 response to the colour change.

## Discussion

In the present study, we tested whether a simple predictive coding account could explain online prediction effects on the amplitude of N1 ERP components elicited by words in biasing contexts. We biased expectations for upcoming words via images of varying predictability. Based on a predictive coding framework, we hypothesised that there would be an interaction between picture-word Predictability and Congruency in which N1 amplitude scales with prediction error. Planned analyses failed to find evidence for this hypothesis, and exploratory analyses revealed, despite strong evidence for prediction effects in the N1, that the direction of the interaction was opposite to that expected under the hypothesis. Specifically, increases in Predictability were associated with greater-amplitude N1s for picture-congruent words, and smaller-amplitude N1s for picture-incongruent words. On this basis, we conclude that a simple predictive coding explanation of the N1 cannot explain predictability effects observed in the picture-word verification task used here.

In recent years, predictive coding models have been increasingly applied to explain neural phenomena observed during language processing. This includes predictive coding perspectives on the N1 specifically (e.g., Gagl et al., 2020; Huang et al., 2022; Zhao et al., 2019), or its likely generator, vOT (Price & Devlin, 2011), and other areas of language processing. For example, consider the well-researched N400 ERP component, generally recognised since its initial identification as capturing activity related to semantic processes (Kutas & Federmeier, 2011; Kutas & Hillyard, 1980). The N400 shows sensitivity to word- and sentence-level surprise or predictability (Delaney-Busch et al., 2019; Lau et al., 2013; Lindborg et al., 2023; Mantegna et al., 2019; Van Petten & Kutas, 1990), in a manner that may be consistent with predictive coding (Bornkessel-Schlesewsky & Schlesewsky, 2019; Eddine et al., 2023; Rabovsky & McRae, 2014). Similar interpretations have been made of other signals, as capturing prediction errors for phonological, semantic, or syntactic representations (Fitz & Chang, 2019; Gagnepain et al., 2012; Van Petten & Luka, 2012; Ylinen et al., 2016, 2017).

Indeed, emerging evidence is beginning to support the broader contention that naturalistic language comprehension utilises a predictive coding hierarchy spanning the language network (Caucheteux et al., 2023; Schuster et al., 2021; Shain et al., 2020; L. Wang et al., 2023). In this way, evidence for predictive coding in language reflects the growing, although not definitive, empirical evidence for predictive coding models in perception more generally (Clark, 2013; Heilbron & Chait, 2018; Hodson et al., 2024; Walsh et al., 2020).

We do not believe our findings refute the existence of predictive coding mechanisms during the N1. This is informed by our review of the literature outlined in the Introduction, in which we found evidence broadly consistent with a predictive coding interpretation of the N1. Instead, we argue that a simple predictive coding account of the N1, in which the component’s amplitude straightforwardly indexes prediction error in a manner dependent on prediction certainty, is insufficient to explain the pattern of effects we observed in the picture-word verification task we used here. For a predictive coding model to better account for these data, it would require elaboration. One feature that may be relevant is the nature of the task. We elected to use a picture-word verification task as it encourages explicit prediction of word forms from non-linguistic contexts.

However, this task paradigm may alter predictive processing of word forms in two key ways. First, participants will have soon learned that the observed word form only matches its preceding image 50% of the time, which could have interacted with the effect of Predictability (prediction certainty) in unexpected ways. Second, the requirement for explicit verification of prediction congruency may have encouraged artificial processing strategies that are not representative of naturalistic word recognition and reading processes. To better understand whether and how such factors influence any possible predictive coding effects on the N1, we could manipulate prediction error magnitude and precision while the participant’s task instructions do not explicitly require processing of the cue. For instance, we could use a picture-word priming design (Sperber et al., 1979; Vanderwart, 1984), presenting picture-word pairs, as in the current study, but ask participants to respond with lexical decisions. Here, prediction error magnitude could be operationalised as the orthographic distance between the string (whether word or non-word), and precision as the predictability of a word given its picture. We believe that such an approach could provide insight into whether, and which, features of the paradigm we used could have resulted in the unexpected pattern of results. Finally, it is possible that dynamics of predictive processing were influenced by the slow presentation rate employed in the present study, relative to more naturalistic reading paradigms.

Indeed, previous research has highlighted the importance of presentation rate in prediction effects during reading (e.g., Dambacher et al., 2012), and recent findings have shown that unpredictability in stimulus presentation timing (e.g., with jittered inter-stimulus intervals) may interfere with predictive processes, as indexed by the mismatch negativity component (Tsogli et al., 2022). This explanation of our results could be tested by study designs examining how the congruency-predictability interaction varies over stimulus onset asynchronies of different durations. In sum, while predictive coding mechanisms may ultimately underlie the pattern of effects we observed, the simple account we have tested requires elaboration, informed by insights from other paradigms, for it to explain why our current pattern of effects is opposite to that expected.

Our study is not the first to identify patterns in evoked responses that seemingly run counter to a simple predictive coding model (e.g., Bowman et al., 2013; Eisenhauer et al., 2022; Mangun & Hillyard, 1991; Vidal-Gran et al., 2020). One suggested elaboration to a simple predictive coding model that could allow expected stimuli to elicit greater evoked responses than unexpected stimuli supposes that, in such cases, expected stimuli may benefit from greater precision than is allotted to deviant stimuli (Bowman et al., 2023; Heilbron & Chait, 2018; Kok et al., 2012). In our experiment, matched picture-congruent and -incongruent words followed the same pictures, such that predictability, which we used as a measure of top-down prediction precision, should have been identical prior to word presentation. However, if top-down effects can penetrate early stages of visual processing that precede the N1, it is conceivable that processing after word presentation, but prior to the N1, could have up-weighted the precision of information in picture-congruent words’ representations, resulting in the observed pattern of effects. In simulations, Bowman et al. (2023) recently demonstrated that precision-modulated predictive coding models can indeed produce ”contra-vanilla” patterns in prediction errors if prediction-congruent stimuli benefit from higher precision, but that this should be expected to affect the evoked response non-linearly. Specifically, the latency of the evoked response should be shorter for the prediction-congruent stimulus. Our findings did indeed reveal a latency difference in the N1 offset period, between congruent and incongruent words at the highest level of predictability (**Figure 10**), but the direction of this difference was opposite to that predicted by Bowman et al. (2023), with a shorter offset period for picture-incongruent words. As such, it is unclear how an elaboration based on differential precision between picture-congruent and-incongruent words may relate to our findings.

We acknowledge the possibility that the insufficiency of predictive coding accounts to explain the data we observed may reflect a more fundamental shortcoming. Indeed, an enduring criticism of predictive coding models is that some evidence for them may also be explained by alternative models (de Lange et al., 2018; Hodson et al., 2024). To speculate, predictive coding models may account for activity in the N1 in previously tested paradigms without accurately describing the underlying neural processes. For instance, Luthra et al. (2021) showed that, in spoken word recognition, interactive activation models may provide an alternative account of the ERP amplitude reduction observed in response to prediction violations, without invoking key features of predictive coding models. Indeed, effects indicative of predictive processing may emerge in a system that that lacks any representations of, or mechanisms implementing, predictions or prediction errors, instead only implementing ”pattern completion” (Falandays et al., 2021). It is tentatively possible that the picture-word verification paradigm we applied here may be a scenario that employs the same neurocognitive processes in the N1 as those employed in other paradigms, but elicits cognitive dynamics whose corresponding neural activity reveals differences from a predictive coding model. It is possible that processing indexed by the N1 can only be explained by a model distinct from the predictive coding framework, even though predictive coding models may correlate with patterns of activity seen in most paradigms. Justifying the development of such a model, distinct from predictive coding, would require much more evidence for the shortcomings of a predictive coding account, and we do not believe our study provides the insights necessary to speculate on the form such a model could take.

Such further insights may be provided by an approach that examines patterns in the representational content of neural activity, rather than univariate patterns of overall activity. Such an approach has been exploited previously as a way of comparing prediction error models, in which neural signals represent unexplained content, with sharpening models of language processing, in which neural signals contain sharper representations of predicted content (Desimone, 1996; Grill-Spector et al., 2006). While these models can account for similar patterns in overall neural activity, they predict dissociable patterns in corresponding representational content (Blank & Davis, 2016).

For instance, Blank and Davis (2016) employed a Congruency (matching, neutral) × Precision (signal quality: 4 or 12 vocoder channels) design in an fMRI experiment on speech perception. Representational similarity analyses of fMRI activity from the posterior superior temporal sulcus showed a pattern consistent with the representational content expected under a prediction error account, and not a sharpening account.

Analyses of EEG signals in a similar paradigm by Sohoglu and Davis (2020) also show evidence for patterns of representational enhancement and suppression that match a prediction error account, from 100 ms after stimulus presentation. Further evidence for prediction error accounts of early speech perception processes is seen in fMRI and MEG analyses of two-syllable words, where precision is quantified as the predictability of syllable two, given syllable one (Sohoglu et al., 2023). In contrast, however, an MEG study on the representation of lexical-semantic information during *visual* word recognition found evidence more consistent with a sharpening account (Eisenhauer et al., 2022). Although their use was motivated by a need to disentangle two explanations of evoked-response patterns that are both consistent with predictive coding, we believe that such analyses, focusing on representational content, may also provide an avenue to further investigate the pattern we observed that was seemingly inconsistent with predictive coding. This could reveal whether the N1’s modulation is accompanied by the representation of more or less stimulus-relevant information, and may more clearly point to the underlying mechanisms.

Representational content is also of particular importance when testing predictive coding accounts because it determines the depth in the hierarchy to which top-down predictions can be conveyed and effectively implemented. This is because in a hierarchical model of predictive coding, where levels of the hierarchy utilise different representational formats, the interaction between ascending input and descending predictions must involve some mapping of higher-level onto lower-level representations. For instance, if semantic context can influence processing that is closer to sensory input and indexed by early ERP components (e.g., Enge et al., 2023; Getz & Toscano, 2019; Segalowitz & Zheng, 2009), then higher-level semantic information must be translated into predictions of upcoming lower-level sensory signals. In the case of our study’s modulation of the N1, if the N1 is implicated in visual-orthographic processing (Bentin et al., 1999; Brem et al., 2018; Ling et al., 2019; Maurer, Brandeis, & McCandliss, 2005), then predictions of upcoming words must be translated into a visual-orthographic code. Such a mapping could be expected to be very computationally lossy; predictions for visual-orthographic features of a single word should be expected to also confer facilitation for words that are orthographically similar, yet picture-incongruent (Kim & Lai, 2012). In contrast, a later ERP more directly implicated in semantic processing, like the N400, may be expected to be less limited by such mappings.

From one perspective, mapping of predictions to lower-level representations may be considered a requisite for a phenomenon to be considered top-down modulation (Rauss et al., 2011). This relates to a long-standing debate on whether prediction effects at the lexical level of language processing necessitate top-down input informed by higher-level semantic processes, or could instead result from perhaps more parsimonious intralexical effects (Fodor, 1983; Forster, 1979). A similar argument could be made that context effects on the N1 could be interpreted as intra-*orthographic*, resulting from local interactions in a possible *orthographic module*. As an example, the orthographic features of the word form *fish* may preactivate features of the word form *chips* simply through learned co-occurrence rather than top-down modulation, entirely within an orthographic processing module that possesses nothing approaching a semantic representation. Such facilitation could be implemented via an extension to classic interactive activation models (e.g., McClelland & Rumelhart, 1981) in which there are excitatory lateral connections between word-level units whose strength is determined by co-occurrence frequency. We consider this point to highlight an advantage of paradigms such as ours, that use non-linguistic contexts (e.g., task instructions, images, etc.) to cue upcoming words and word forms. Effects of context that map across representations in this way necessitate transfer of information across levels of the processing hierarchy, and may thus be considered stronger evidence for an influence of top-down predictions.

An aspect of the predictive coding account that our design did not fully test also relates to this idea of representational mapping. We dichotomised the variable of congruency (prediction error magnitude), with orthographic Levenshtein distance maximised between picture-congruent and -incongruent word forms. However, prediction error magnitude should also be expected to vary continuously, from unpredicted word forms that are less to more orthographically similar to the predicted word form. This is comparable to Gagl et al.’s (2020) use of a pixel distance metric to calculate the continuous distance between a presented word form and a context-neutral prior. Such an approach could be applied to biasing contexts by instead calculating the orthographic distance between a presented word form and a context-informed prior, where the probability of observing certain pixels (or orthographic features) could be up-weighted proportional to prediction certainty. We believe such an approach could provide useful insights in elucidating the pattern of effects we observed.

We note that exploratory analyses at the typical latency of the N400 revealed a pattern which also appears to run counter to a simple predictive coding account of predictability effects. This is seemingly inconsistent with interpretations of the N400 as indexing prediction error (Bornkessel-Schlesewsky & Schlesewsky, 2019; Eddine et al., 2023; Rabovsky & McRae, 2014). At the lowest level of predictability, we observed greater N400 amplitudes for picture-congruent words, than for picture-incongruent words. As predictability increased, meanwhile, N400 amplitudes became less extreme for picture-congruent words, rather than becoming more extreme for picture-incongruent words. We caution against over-interpreting these results. In addition to these results being entirely exploratory, we used an average EEG reference, rather than the more standard mastoid reference for later centroparietal components like the N400.

Furthermore, when elicited by words in context, such as a sentence or picture, evidence suggests that the N400 indexes both prediction and integration processes (Nieuwland et al., 2018). Nevertheless, that our design elicited effects on the N400 that are seemingly inconsistent with existing findings from more traditional experimental designs may point to our specific experimental design playing a key role in the pattern of effects we observed in the N1.

In sum, we tested a simple predictive coding account of the word-elicited N1, but failed to find evidence in favour of it. Exploratory analyses suggest that the pattern of effects in the Congruency-Predictability interaction were in the opposite direction to that expected under a simple predictive coding model. We argue that such a model is insufficient to explain the pattern of effects we observed, and we have identified avenues of future research that could better delineate how predictive processes interact with processing during the N1.

