## Supplementary Materials for "Can Prediction Error Explain Predictability Effects on the N1 during Picture-Word Verification?"

Contents

**A Picture-Word Stimuli** 3

**B Behavioural Validation Results** 18

**C Localiser Task Stimuli** 22

**D Statistical Power Analysis** 30

**E Instructions given to Participants** 37

**F Change to the High-Pass Filter Cut-Off** 39

**J Model Estimates for the N400** 60

**K Checking ERPs Time-Locked to Pictures** **61**

**L Checking ERPs Time-Locked to Responses** **64**

#### A Picture-Word Stimuli

There were 200 words per Congruency condition, with one congruent and one incongruent word per image. The experimental stimuli are summarised in **Figure 1**. First, stimuli were subset according to norms collected by Brysbaert et al. (2019), such that at least 90% of participants knew each word. Stimuli were additionally subset such that all words were nouns according to the dominant part of speech data from SUBTLEX-UK (van Heuven et al., 2014), and had a mean concreteness rating above 4 (on a Likert scale from 1, least concrete, to 5, most concrete) according to Brysbaert et al. (2014). Images were taken from the Bank of Online Standardised Stimuli (BOSS) norms (Brodeur et al., 2014), a large database of images with normed statistics, including percentage of name agreement, which, critically, we used as a measure of Predictability. Words were identified as possible picture-congruent words if they were listed as the most frequent (i.e., modal) name for any image in the BOSS norms, and were identified as possible picture-incongruent words if they were not.

Picture-congruent and -incongruent words were matched item-wise across five lexical variables, with specific tolerance ranges, as follows: (1) word length (number of characters), exactly; (2) concreteness according to Brysbaert et al. (2014), within $\pm.25$; (3) Zipf frequency (a logarithmic scale of word frequency) according to SUBTLEX-UK, within $\pm.125$; (4) character bigram probability (calculated from SUBTLEX-UK), within $\pm.0025$; and (5) OLD20 (the average Orthographic Levenshtein Distance of the 20 closest neighbours to a given word; Yarkoni et al., 2008) calculated from the LexOPS inbuilt dataset, within $\pm.75$. To ensure that picture-incongruent words were not inadvertent possible descriptors for images, the cosine positive pointwise mutual information (PPMI) measure of associative semantic similarity calculated from the Small World of Words (SWOW) word association norms (De Deyne et al., 2019) was minimised to be $\leq.01$ between each image's matched picture-congruent and picture-incongruent words. To ensure picture-incongruent words did not share orthographic features with their respective

**Figure 1**

*Summary of the picture-word stimuli.*

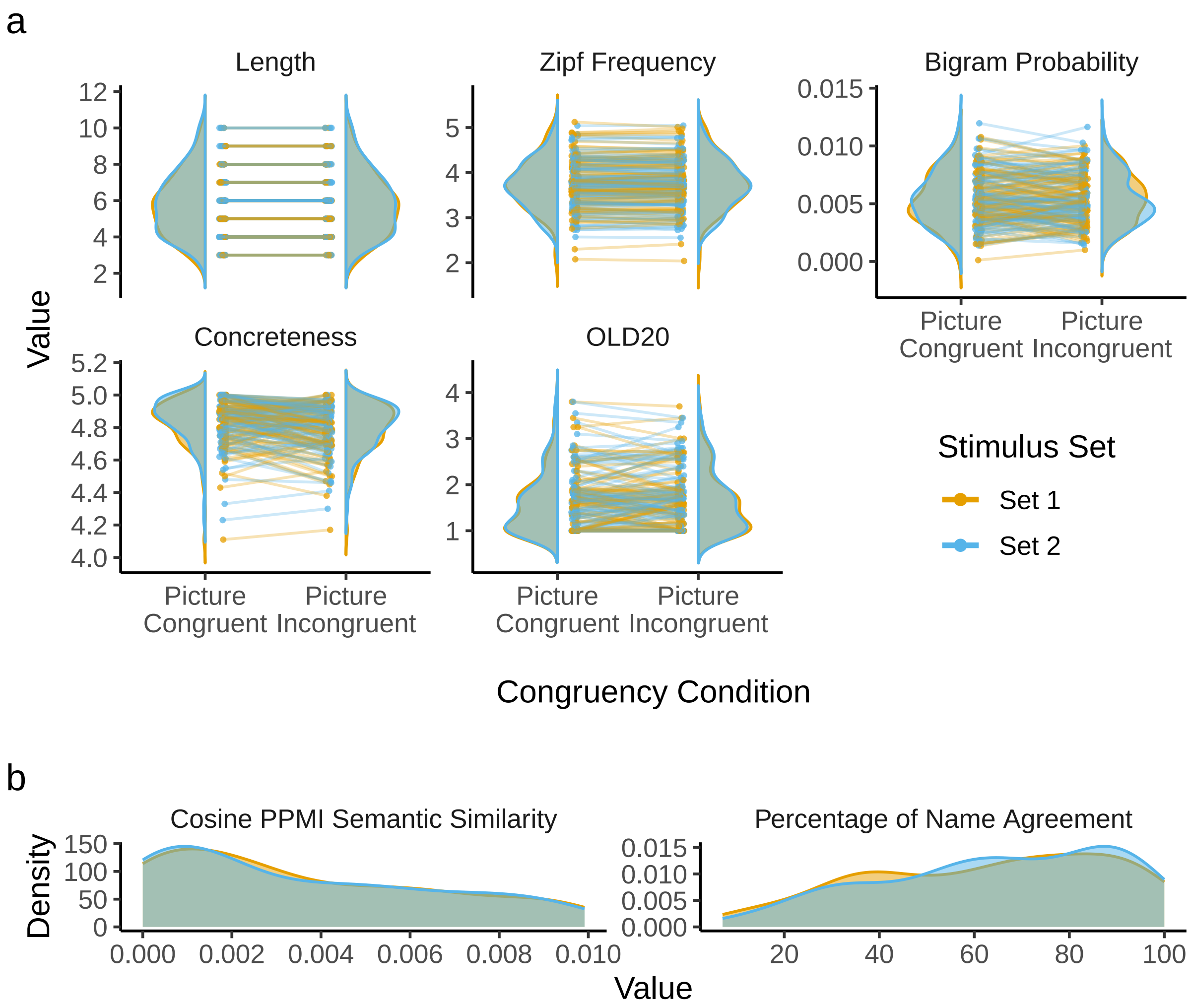

Each panel depicts how a single variable was controlled. (**a**) Probability densities for variables which were matched item-wise between picture-congruent and picture-incongruent conditions, and distribution-wise between counterbalanced stimulus Sets 1 (in *yellow*) and 2 (in *blue*). Points representing pairs of words which are matched item-wise are joined by lines. Points' positions are jittered slightly along the x-axis for visibility. (**b**) Probability densities for two variables matched only in a distribution-wise manner between the counterbalanced stimulus sets: Cosine PPMI (Positive Pointwise Mutual Information) Semantic Similarity from SWOW (Small World of Words; De Deyne et al., 2019), and modal name agreement from the BOSS norms. These variables cannot be matched between Congruency conditions because only a single value describes each matched congruent-incongruent word pair.

picture-congruent words, orthographic Levenshtein distance between matched items was maximised. As items were also matched in word length, this meant all matched pairs of words had a Levenshtein distance equal to their number of characters. The variable used to index the Predictability of picture-congruent words was percentage of modal name agreement, which was sampled pseudo-randomly (picture-congruent words were not selected if no incongruent match could be identified fitting the constraints specified above) from the BOSS norms, and varied continuously in the generated stimuli from 7 to 100%.

As the participants were recruited in the United Kingdom, possible congruent and incongruent picture-word pairs were excluded if we identified the words as less frequent in British English (e.g., *sidewalk*) or if they were modal names for images that the Canadian participants of the BOSS norms are likely to have been more able to name or distinguish (e.g., *buffalo*, *bison*). In addition, picture-word pairs were excluded if words were identified as shortened versions of nouns (e.g., *limo*, *chimp*) or alternate names for the same object (e.g., *motorbike*, *motorcycle*). Candidate picture-incongruent words were additionally excluded if images were not representative of the images in the BOSS (e.g., *waiter* or *church*, as there were no other images of people or entire buildings in the BOSS), or if they were unimageable despite their high concreteness value (e.g., *item*). Plural words (e.g., *sticks*) were excluded, as most images in the BOSS have modal names that are singular. Finally, four images with modal names *nut*, *trumpet*, *spinach*, and *tuba* were excluded, as we judged these names to be incorrect descriptions of their images.

To avoid repetition effects, each image was presented once, with participants viewing either the associated picture-congruent or picture-incongruent word. This was counterbalanced by splitting the stimuli pseudo-randomly into two equally sized stimulus sets, referred to as Set 1 and Set 2. Each participant was presented with only one of these stimulus sets. Pictures followed by congruent words in Set 1 were followed by incongruent words in Set 2, and vice versa. To minimise any systematic difference between the counterbalanced groups, the split of stimuli was selected to maximise the empirical distributional overlap (Pastore & Calcagnì, 2019) between the two stimulus sets in relevant variables. Specifically, the stimulus sets were selected from 50,000 random splits to maximise the overlap between the distributions of the following seven variables: (1) percentage of modal name agreement according to the BOSS norms; (2) cosine PPMI semantic similarity according to the SWOW; (3, 4) Zipf word frequency and character bigram probability according to SUBTLEX-UK; (5) word concreteness (Brysbaert et al., 2014); (6) word length; and (7) OLD20. Variables that were also matched item-wise between the conditions were matched distribution-wise separately within each Congruency condition. This ensured there were minimal systematic differences in distributions between conditions or stimulus sets.

To generate stimuli for practice trials, 20 matched pairs of picture-congruent and -incongruent words were generated using the same pipeline as above, except that word frequency, word concreteness, and character bigram probability were not matched item-wise. The practice stimuli were generated from images and words not used in the experimental stimuli. The same practice trials were presented to all participants.

**Table 1**

*All stimuli for the picture-word task. Column Image IDs are unique file names given to each image in the BOSS, while %Agree reports the percentage of modal name agreement for the image in the BOSS. Set refers to the assigned stimulus sets. Column* Word *contains the matched congruent (*C*) and incongruent (*I*) words associated with each image. The remaining columns are as follows, separating values into those for the congruent (*C*) and incongruent (*I*) words where possible: Length = number of characters;* Zipf *= Zipf frequency in SUBTLEX-UK;* OLD20 *= OLD20 values in the LexOPS dataset;* BG *= mean character bigram probabilities in SUBTLEX-UK;* CNC *= mean concreteness ratings in Brysbaert et al. (2014);* Cosine PPMI *= cosine positive pointwise mutual information values of semantic associative similarity between matched congruent and incongruent words from the Small World of Words. Rows are numbered for ease of reference.*

| | Image ID | %Agree | Set | Word | | Length | | Zipf | | OLD20 | | BG | | CNC | | Cosine PPMI |
|---|---|---|---|---|---|---|---|---|---|---|---|---|---|---|---|---|
| | | | | C | I | C | I | C | I | C | I | C | I | C | I | |
| 1 | joustingspear | 7% | 2 | spear | porch | 5 | 5 | 3.42 | 3.47 | 1.20 | 1.15 | .0062 | .0047 | 5.00 | 4.92 | .0071 |
| 2 | cabasa | 10% | 1 | shaker | trough | 6 | 6 | 3.35 | 3.32 | 1.00 | 1.65 | .0092 | .0067 | 4.11 | 4.17 | .0073 |
| 3 | powerchair | 10% | 1 | scooter | missile | 7 | 7 | 3.63 | 3.66 | 1.60 | 1.70 | .0077 | .0057 | 4.96 | 4.83 | .0065 |
| 4 | pottery | 12% | 1 | pottery | rainbow | 7 | 7 | 4.14 | 4.18 | 1.65 | 2.40 | .0070 | .0069 | 4.72 | 4.57 | .0093 |
| 5 | lbracket01 | 13% | 1 | bracket | tornado | 7 | 7 | 3.40 | 3.49 | 1.75 | 2.10 | .0036 | .0059 | 4.43 | 4.53 | .0003 |
| 6 | flail | 14% | 2 | mace | knob | 4 | 4 | 3.37 | 3.50 | 1.00 | 1.35 | .0042 | .0029 | 4.81 | 4.75 | .0017 |
| 7 | plastictube | 16% | 1 | tube | chip | 4 | 4 | 4.28 | 4.26 | 1.00 | 1.00 | .0031 | .0051 | 4.82 | 4.71 | .0008 |
| 8 | paintscraper | 16% | 2 | scraper | nightie | 7 | 7 | 2.80 | 2.89 | 1.75 | 1.85 | .0052 | .0037 | 4.23 | 4.30 | .0002 |
| 9 | pillar | 19% | 2 | pillar | sewage | 6 | 6 | 3.54 | 3.58 | 1.60 | 1.95 | .0060 | .0041 | 4.77 | 4.52 | .0016 |
| 10 | bazooka | 19% | 1 | bazooka | sunburn | 7 | 7 | 2.76 | 2.86 | 2.55 | 1.85 | .0015 | .0026 | 4.66 | 4.57 | .0037 |
| 11 | chocolatecroissant | 21% | 1 | pastry | weapon | 6 | 6 | 4.37 | 4.29 | 1.55 | 1.90 | .0057 | .0067 | 4.97 | 4.76 | .0011 |
| 12 | solderingwire | 21% | 2 | wire | pond | 4 | 4 | 4.29 | 4.20 | 1.00 | 1.00 | .0089 | .0097 | 4.72 | 4.90 | .0084 |
| 13 | hedgeshears | 24% | 2 | shears | tendon | 6 | 6 | 2.94 | 2.98 | 1.40 | 1.55 | .0120 | .0103 | 4.61 | 4.47 | .0013 |

| | Image ID | %Agree | Set | Word | | Length | | Zipf | | OLD20 | | BG | | CNC | | Cosine PPMI |
|---|---|---|---|---|---|---|---|---|---|---|---|---|---|---|---|---|
| | | | | C | I | C | I | C | I | C | I | C | I | C | I | |
| 14 | pouch01b | 26% | 1 | pouch | ledge | 5 | 5 | 3.36 | 3.38 | 1.05 | 1.10 | .0069 | .0051 | 4.50 | 4.72 | 0 |
| 15 | ram | 27% | 2 | ram | pup | 3 | 3 | 3.65 | 3.74 | 1.00 | 1.00 | .0037 | .0014 | 4.55 | 4.61 | .0083 |
| 16 | oats | 28% | 1 | oats | lice | 4 | 4 | 3.33 | 3.29 | 1.00 | 1.00 | .0041 | .0051 | 4.78 | 4.73 | .0059 |
| 17 | bandage | 28% | 2 | bandage | whisker | 7 | 7 | 3.22 | 3.10 | 1.80 | 1.65 | .0071 | .0087 | 4.85 | 4.70 | .0087 |
| 18 | bastingbrush | 28% | 2 | brush | stamp | 5 | 5 | 4.29 | 4.20 | 1.35 | 1.30 | .0029 | .0050 | 4.54 | 4.70 | .0021 |
| 19 | rug01 | 29% | 1 | rug | soy | 3 | 3 | 3.57 | 3.64 | 1.00 | 1.00 | .0014 | .0028 | 4.79 | 4.70 | 0 |
| 20 | radio01 | 29% | 1 | radio | smile | 5 | 5 | 4.82 | 4.71 | 1.40 | 1.00 | .0036 | .0040 | 4.74 | 4.50 | .0012 |
| 21 | tonfa | 29% | 2 | baton | yeast | 5 | 5 | 3.53 | 3.64 | 1.00 | 1.50 | .0097 | .0076 | 4.64 | 4.72 | .0036 |
| 22 | salsa | 29% | 1 | salsa | trunk | 5 | 5 | 3.82 | 3.93 | 1.20 | 1.15 | .0038 | .0025 | 4.70 | 4.71 | .0010 |
| 23 | smokedsalmon | 29% | 2 | salmon | tunnel | 6 | 6 | 4.34 | 4.23 | 1.25 | 1.65 | .0058 | .0039 | 4.81 | 4.82 | .0007 |
| 24 | handmixer01d | 31% | 2 | mixer | wedge | 5 | 5 | 3.45 | 3.45 | 1.40 | 1.25 | .0056 | .0048 | 4.33 | 4.41 | .0019 |
| 25 | videotape01b | 31% | 1 | cassette | revolver | 8 | 8 | 2.94 | 2.85 | 1.95 | 1.75 | .0058 | .0079 | 4.60 | 4.69 | .0019 |
| 26 | woodboard | 31% | 2 | wood | ship | 4 | 4 | 4.78 | 4.77 | 1.00 | 1.00 | .0034 | .0047 | 4.85 | 4.87 | .0097 |
| 27 | jar03 | 33% | 2 | jar | lip | 3 | 3 | 3.96 | 3.91 | 1.00 | 1.00 | .0055 | .0035 | 5.00 | 4.96 | .0039 |
| 28 | cuttingpliers02 | 33% | 2 | pliers | beanie | 6 | 6 | 2.73 | 2.80 | 1.65 | 1.35 | .0070 | .0082 | 4.93 | 4.74 | .0019 |
| 29 | kalashnikov | 33% | 2 | rifle | altar | 5 | 5 | 3.62 | 3.58 | 1.65 | 1.00 | .0042 | .0063 | 4.85 | 4.85 | .0057 |
| 30 | overalls | 33% | 1 | overalls | mongoose | 8 | 8 | 3.01 | 2.94 | 2.00 | 2.70 | .0079 | .0070 | 4.74 | 4.89 | .0025 |
| 31 | towel01 | 34% | 1 | towel | spine | 5 | 5 | 3.87 | 3.91 | 1.30 | 1.00 | .0077 | .0090 | 4.86 | 4.88 | .0090 |
| 32 | branch02 | 36% | 1 | branch | powder | 6 | 6 | 4.10 | 4.17 | 1.15 | 1.55 | .0064 | .0065 | 4.90 | 4.76 | 0 |
| 33 | ribbon03a | 36% | 2 | lace | beak | 4 | 4 | 3.73 | 3.83 | 1.00 | 1.00 | .0041 | .0059 | 4.85 | 4.96 | .0023 |

| | Image ID | %Agree | Set | Word | | Length | | Zipf | | OLD20 | | BG | | CNC | | Cosine PPMI |
|---|---|---|---|---|---|---|---|---|---|---|---|---|---|---|---|---|
| | | | | C | I | C | I | C | I | C | I | C | I | C | I | |
| 34 | yarn | 36% | 1 | yarn | twig | 4 | 4 | 3.14 | 3.22 | 1.00 | 1.20 | .0041 | .0028 | 4.93 | 4.75 | 0 |
| 35 | napkin | 36% | 1 | napkin | weasel | 6 | 6 | 3.31 | 3.33 | 1.90 | 1.60 | .0062 | .0076 | 4.93 | 4.74 | .0028 |
| 36 | bag | 36% | 1 | bag | oil | 3 | 3 | 4.89 | 4.98 | 1.00 | 1.00 | .0021 | .0033 | 4.90 | 4.93 | .0026 |
| 37 | mussel | 36% | 2 | clam | sash | 4 | 4 | 3.35 | 3.37 | 1.00 | 1.00 | .0029 | .0050 | 4.89 | 4.67 | .0033 |
| 38 | tray | 37% | 1 | tray | sail | 4 | 4 | 4.15 | 4.17 | 1.00 | 1.00 | .0038 | .0035 | 4.74 | 4.59 | .0032 |
| 39 | brainmodel | 38% | 1 | brain | river | 5 | 5 | 4.84 | 4.93 | 1.00 | 1.00 | .0086 | .0094 | 4.69 | 4.89 | .0014 |
| 40 | megaphone | 38% | 1 | megaphone | billiards | 9 | 9 | 2.89 | 2.89 | 2.85 | 2.30 | .0050 | .0046 | 4.76 | 4.61 | .0018 |
| 41 | foodprocessor | 38% | 2 | blender | javelin | 7 | 7 | 3.32 | 3.35 | 1.45 | 1.85 | .0098 | .0087 | 5.00 | 4.90 | .0022 |
| 42 | slide02 | 38% | 2 | slide | trail | 5 | 5 | 4.17 | 4.27 | 1.15 | 1.10 | .0036 | .0038 | 4.48 | 4.46 | .0078 |
| 43 | turnip | 38% | 2 | turnip | nickel | 6 | 6 | 3.36 | 3.27 | 1.70 | 1.35 | .0024 | .0040 | 4.79 | 4.79 | .0013 |
| 44 | oyster02 | 38% | 1 | oyster | canvas | 6 | 6 | 3.82 | 3.95 | 1.55 | 1.80 | .0080 | .0068 | 4.85 | 4.78 | .0017 |
| 45 | giftbow02b | 39% | 1 | bow | jam | 3 | 3 | 4.22 | 4.34 | 1.00 | 1.00 | .0040 | .0018 | 4.61 | 4.71 | .0072 |
| 46 | mask02a | 39% | 1 | mask | pony | 4 | 4 | 4.04 | 3.96 | 1.00 | 1.00 | .0046 | .0059 | 4.96 | 4.90 | .0045 |
| 47 | bulldozer | 40% | 1 | bulldozer | pepperoni | 9 | 9 | 2.91 | 2.95 | 2.50 | 2.70 | .0055 | .0067 | 4.90 | 5.00 | 0 |
| 48 | iceberglettuce | 41% | 1 | lettuce | pyramid | 7 | 7 | 3.81 | 3.71 | 2.40 | 2.50 | .0038 | .0022 | 4.97 | 4.96 | .0031 |
| 49 | leek | 42% | 1 | leek | moat | 4 | 4 | 3.56 | 3.69 | 1.00 | 1.00 | .0047 | .0045 | 4.92 | 4.69 | .0013 |
| 50 | scalpel | 43% | 1 | scalpel | tequila | 7 | 7 | 3.10 | 3.19 | 1.85 | 2.60 | .0043 | .0034 | 4.86 | 4.77 | 0 |
| 51 | pipe | 43% | 2 | pipe | taxi | 4 | 4 | 4.26 | 4.30 | 1.00 | 1.00 | .0019 | .0016 | 4.88 | 4.93 | .0011 |
| 52 | glassescase | 44% | 1 | wallet | brandy | 6 | 6 | 3.81 | 3.83 | 1.20 | 1.25 | .0074 | .0077 | 4.81 | 4.81 | .0038 |
| 53 | coaster | 44% | 2 | tile | mast | 4 | 4 | 3.56 | 3.51 | 1.00 | 1.00 | .0068 | .0080 | 4.68 | 4.92 | .0057 |

| | Image ID | %Agree | Set | Word | | Length | | Zipf | | OLD20 | | BG | | CNC | | Cosine PPMI |
|---|---|---|---|---|---|---|---|---|---|---|---|---|---|---|---|---|
| | | | | C | I | C | I | C | I | C | I | C | I | C | I | |
| 54 | lectern01 | 45% | 2 | podium | liquid | 6 | 6 | 4.17 | 4.25 | 1.85 | 1.75 | .0018 | .0022 | 4.89 | 4.72 | .0032 |
| 55 | doorlock | 46% | 2 | lock | rail | 4 | 4 | 4.42 | 4.40 | 1.00 | 1.00 | .0028 | .0040 | 4.65 | 4.90 | .0045 |
| 56 | puzzle | 48% | 1 | puzzle | sketch | 6 | 6 | 3.93 | 3.85 | 1.65 | 1.70 | .0018 | .0031 | 4.75 | 4.56 | .0069 |
| 57 | rhinoceros02 | 48% | 2 | rhinoceros | aftershave | 10 | 10 | 3.07 | 3.14 | 3.55 | 3.35 | .0084 | .0075 | 4.75 | 4.56 | .0012 |
| 58 | box01a | 49% | 1 | box | sun | 3 | 3 | 5.12 | 5.01 | 1.00 | 1.00 | .0015 | .0028 | 4.90 | 4.83 | .0033 |
| 59 | star | 50% | 2 | star | wall | 4 | 4 | 5.04 | 5.05 | 1.00 | 1.00 | .0086 | .0079 | 4.69 | 4.86 | .0015 |
| 60 | scanner | 50% | 1 | scanner | bedding | 7 | 7 | 3.46 | 3.55 | 1.75 | 1.35 | .0089 | .0093 | 4.79 | 4.61 | 0 |
| 61 | mug05 | 50% | 2 | mug | wax | 3 | 3 | 3.93 | 3.87 | 1.00 | 1.00 | .0014 | .0027 | 4.80 | 4.97 | .0069 |
| 62 | ladle02a | 51% | 1 | ladle | tiara | 5 | 5 | 3.21 | 3.16 | 1.50 | 1.45 | .0041 | .0059 | 4.90 | 4.89 | .0050 |
| 63 | humanskeleton | 52% | 1 | skeleton | tortoise | 8 | 8 | 3.78 | 3.82 | 2.05 | 2.60 | .0073 | .0089 | 4.97 | 4.87 | .0025 |
| 64 | gecko | 52% | 1 | lizard | barley | 6 | 6 | 3.73 | 3.61 | 1.60 | 1.00 | .0038 | .0051 | 4.68 | 4.59 | .0009 |
| 65 | boxtrailer | 52% | 2 | trailer | receipt | 7 | 7 | 3.68 | 3.68 | 1.70 | 2.20 | .0071 | .0049 | 4.79 | 4.86 | .0058 |
| 66 | mechanicalpencil02 | 53% | 1 | pencil | kidney | 6 | 6 | 3.98 | 3.94 | 1.90 | 1.70 | .0047 | .0030 | 4.88 | 4.96 | .0086 |
| 67 | spatula03 | 54% | 2 | spatula | airship | 7 | 7 | 2.95 | 2.83 | 2.05 | 2.35 | .0039 | .0040 | 4.96 | 4.92 | .0002 |
| 68 | fusilli03a | 54% | 1 | pasta | motor | 5 | 5 | 4.19 | 4.25 | 1.00 | 1.60 | .0066 | .0082 | 4.86 | 4.84 | .0022 |
| 69 | bracelet01 | 54% | 2 | bracelet | postcard | 8 | 8 | 3.79 | 3.72 | 2.60 | 2.60 | .0047 | .0048 | 4.96 | 4.93 | .0047 |
| 70 | riverotter | 55% | 2 | otter | wrist | 5 | 5 | 3.80 | 3.84 | 1.00 | 1.15 | .0090 | .0073 | 4.86 | 4.93 | .0042 |
| 71 | grandpiano | 55% | 2 | piano | salad | 5 | 5 | 4.36 | 4.38 | 1.10 | 1.00 | .0072 | .0049 | 4.90 | 4.97 | 0 |
| 72 | canoepaddle02 | 55% | 2 | paddle | buzzer | 6 | 6 | 3.73 | 3.73 | 1.30 | 1.70 | .0029 | .0046 | 4.80 | 4.66 | .0039 |
| 73 | suitcase | 56% | 2 | suitcase | pavement | 8 | 8 | 3.78 | 3.68 | 2.85 | 2.10 | .0057 | .0070 | 4.97 | 4.72 | .0014 |

| | Image ID | %Agree | Set | Word | | Length | | Zipf | | OLD20 | | BG | | CNC | | Cosine PPMI |
|---|---|---|---|---|---|---|---|---|---|---|---|---|---|---|---|---|
| | | | | C | I | C | I | C | I | C | I | C | I | C | I | |
| 74 | aquarium | 57% | 1 | aquarium | textbook | 8 | 8 | 3.33 | 3.38 | 2.45 | 2.75 | .0028 | .0028 | 4.77 | 4.86 | .0026 |
| 75 | trombone | 57% | 2 | trombone | mosquito | 8 | 8 | 3.27 | 3.29 | 2.50 | 2.55 | .0058 | .0051 | 4.90 | 4.88 | 0 |
| 76 | spaghetti01 | 57% | 1 | spaghetti | underwear | 9 | 9 | 3.79 | 3.72 | 3.25 | 2.60 | .0071 | .0084 | 5.00 | 4.96 | .0061 |
| 77 | thimble | 58% | 2 | thimble | oregano | 7 | 7 | 2.98 | 3.00 | 1.80 | 2.15 | .0106 | .0096 | 5.00 | 4.81 | 0 |
| 78 | syringe01 | 58% | 2 | syringe | mascara | 7 | 7 | 3.12 | 3.05 | 1.85 | 1.80 | .0080 | .0058 | 4.81 | 4.93 | 0 |
| 79 | antenna | 59% | 2 | antenna | sirloin | 7 | 7 | 3.01 | 2.95 | 1.95 | 2.70 | .0087 | .0067 | 4.75 | 4.66 | 0 |
| 80 | notebook03a | 59% | 1 | notebook | pendulum | 8 | 8 | 3.32 | 3.30 | 2.75 | 2.70 | .0044 | .0049 | 4.92 | 4.69 | .0001 |
| 81 | cleaver01 | 59% | 2 | knife | album | 5 | 5 | 4.49 | 4.55 | 1.75 | 1.75 | .0021 | .0033 | 4.90 | 4.69 | .0037 |
| 82 | honeydewmelon | 59% | 2 | melon | timer | 5 | 5 | 3.49 | 3.40 | 1.00 | 1.00 | .0084 | .0097 | 4.78 | 4.69 | .0003 |
| 83 | platypus | 60% | 2 | platypus | campfire | 8 | 8 | 2.82 | 2.94 | 2.75 | 2.55 | .0035 | .0050 | 4.83 | 4.79 | .0014 |
| 84 | shelf | 60% | 1 | shelf | trout | 5 | 5 | 4.02 | 3.96 | 1.50 | 1.00 | .0108 | .0087 | 4.96 | 4.72 | <.0001 |
| 85 | macaroni01 | 60% | 2 | macaroni | bookcase | 8 | 8 | 3.08 | 3.02 | 1.95 | 2.70 | .0068 | .0044 | 4.97 | 4.93 | .0014 |
| 86 | apricot | 61% | 1 | peach | valve | 5 | 5 | 3.62 | 3.53 | 1.00 | 1.55 | .0054 | .0050 | 4.90 | 4.83 | 0 |
| 87 | seaturtle | 62% | 1 | turtle | pelvis | 6 | 6 | 3.64 | 3.53 | 1.65 | 1.75 | .0039 | .0048 | 5.00 | 4.93 | .0012 |
| 88 | triangle | 62% | 1 | triangle | lighting | 8 | 8 | 3.93 | 4.00 | 1.85 | 1.35 | .0072 | .0086 | 4.52 | 4.38 | .0037 |
| 89 | vulture | 62% | 1 | vulture | measles | 7 | 7 | 3.20 | 3.12 | 1.80 | 1.90 | .0052 | .0074 | 4.73 | 4.69 | .0024 |
| 90 | balcony02 | 64% | 1 | balcony | seaweed | 7 | 7 | 3.83 | 3.83 | 1.90 | 1.85 | .0056 | .0063 | 4.68 | 4.89 | .0033 |
| 91 | adjustablewrench01b | 64% | 2 | wrench | blouse | 6 | 6 | 3.15 | 3.08 | 1.60 | 1.75 | .0076 | .0078 | 4.93 | 4.96 | .0048 |
| 92 | cane | 64% | 2 | cane | reef | 4 | 4 | 3.75 | 3.79 | 1.00 | 1.00 | .0107 | .0087 | 4.87 | 4.70 | .0053 |
| 93 | shield02 | 64% | 2 | shield | packet | 6 | 6 | 3.80 | 3.90 | 1.70 | 1.45 | .0051 | .0036 | 4.66 | 4.46 | 0 |

| | Image ID | %Agree | Set | Word | | Length | | Zipf | | OLD20 | | BG | | CNC | | Cosine PPMI |
|---|---|---|---|---|---|---|---|---|---|---|---|---|---|---|---|---|
| | | | | C | I | C | I | C | I | C | I | C | I | C | I | |
| 94 | tank | 64% | 1 | tank | seed | 4 | 4 | 4.34 | 4.24 | 1.00 | 1.00 | .0086 | .0071 | 4.80 | 4.71 | .0066 |
| 95 | straw | 66% | 2 | straw | badge | 5 | 5 | 4.17 | 4.06 | 1.00 | 1.00 | .0047 | .0025 | 4.77 | 4.93 | .0020 |
| 96 | pickle01a | 66% | 2 | pickle | magnet | 6 | 6 | 3.66 | 3.70 | 1.10 | 1.70 | .0034 | .0039 | 4.64 | 4.70 | .0081 |
| 97 | axe01 | 67% | 1 | axe | rum | 3 | 3 | 3.85 | 3.91 | 1.00 | 1.00 | .0001 | .0010 | 5.00 | 4.93 | .0023 |
| 98 | boat | 67% | 1 | boat | card | 4 | 4 | 4.89 | 4.89 | 1.00 | 1.00 | .0044 | .0059 | 4.93 | 4.90 | .0093 |
| 99 | bowl01 | 67% | 1 | bowl | neck | 4 | 4 | 4.69 | 4.65 | 1.00 | 1.00 | .0027 | .0044 | 4.87 | 5.00 | .0063 |
| 100 | plunger02 | 67% | 2 | plunger | caribou | 7 | 7 | 3.03 | 2.93 | 1.65 | 1.95 | .0072 | .0073 | 4.96 | 4.92 | .0069 |
| 101 | panda | 67% | 1 | panda | lever | 5 | 5 | 3.73 | 3.66 | 1.00 | 1.00 | .0091 | .0100 | 4.75 | 4.77 | .0008 |
| 102 | toothpick02 | 67% | 2 | toothpick | periscope | 9 | 9 | 2.79 | 2.81 | 3.35 | 2.65 | .0090 | .0069 | 4.93 | 4.78 | .0011 |
| 103 | kettle01 | 67% | 2 | kettle | picnic | 6 | 6 | 4.02 | 4.04 | 1.45 | 1.90 | .0041 | .0027 | 4.75 | 4.83 | 0 |
| 104 | lime | 67% | 2 | lime | swan | 4 | 4 | 4.09 | 3.98 | 1.00 | 1.00 | .0061 | .0083 | 4.96 | 4.96 | .0031 |
| 105 | razor01 | 68% | 1 | razor | strap | 5 | 5 | 3.69 | 3.62 | 1.75 | 1.00 | .0038 | .0049 | 4.90 | 4.79 | .0021 |
| 106 | sailboat | 69% | 1 | sailboat | knapsack | 8 | 8 | 2.08 | 2.04 | 2.50 | 3.00 | .0034 | .0020 | 4.89 | 4.90 | .0089 |
| 107 | ribbon04 | 69% | 1 | ribbon | bunker | 6 | 6 | 3.58 | 3.63 | 1.85 | 1.30 | .0047 | .0065 | 4.89 | 4.79 | .0015 |
| 108 | barn | 69% | 2 | barn | menu | 4 | 4 | 4.32 | 4.36 | 1.00 | 1.00 | .0048 | .0070 | 4.79 | 4.67 | .0009 |
| 109 | moon | 69% | 2 | moon | seat | 4 | 4 | 4.74 | 4.78 | 1.00 | 1.00 | .0072 | .0088 | 4.90 | 4.78 | .0001 |
| 110 | parrot01 | 69% | 2 | parrot | sleeve | 6 | 6 | 3.84 | 3.88 | 1.65 | 1.70 | .0053 | .0054 | 5.00 | 4.84 | .0016 |
| 111 | bacon | 71% | 1 | bacon | photo | 5 | 5 | 4.34 | 4.42 | 1.00 | 1.55 | .0067 | .0063 | 4.90 | 4.93 | 0 |
| 112 | americangoldfinch | 71% | 2 | bird | cake | 4 | 4 | 4.85 | 4.81 | 1.00 | 1.00 | .0021 | .0039 | 5.00 | 4.81 | .0035 |
| 113 | cheetah | 71% | 1 | cheetah | stopper | 7 | 7 | 3.45 | 3.39 | 2.20 | 1.55 | .0090 | .0083 | 4.70 | 4.83 | 0 |

| | Image ID | %Agree | Set | Word | | Length | | Zipf | | OLD20 | | BG | | CNC | | Cosine PPMI |
|---|---|---|---|---|---|---|---|---|---|---|---|---|---|---|---|---|
| | | | | C | I | C | I | C | I | C | I | C | I | C | I | |
| 114 | seagull | 71% | 2 | seagull | apricot | 7 | 7 | 3.30 | 3.29 | 2.50 | 2.40 | .0053 | .0045 | 5.00 | 4.97 | .0008 |
| 115 | nail | 72% | 2 | nail | sofa | 4 | 4 | 4.18 | 4.22 | 1.00 | 1.00 | .0034 | .0047 | 4.93 | 4.90 | .0079 |
| 116 | starfish01 | 72% | 2 | starfish | armchair | 8 | 8 | 3.27 | 3.31 | 2.15 | 2.80 | .0065 | .0056 | 4.90 | 5.00 | .0054 |
| 117 | pill | 72% | 1 | pill | knot | 4 | 4 | 3.81 | 3.74 | 1.00 | 1.05 | .0050 | .0045 | 4.72 | 4.87 | .0006 |
| 118 | acorn | 73% | 1 | acorn | bugle | 5 | 5 | 3.13 | 3.01 | 1.65 | 1.25 | .0056 | .0033 | 4.96 | 4.84 | .0065 |
| 119 | shorts01 | 74% | 1 | shorts | needle | 6 | 6 | 3.82 | 3.93 | 1.35 | 1.55 | .0052 | .0058 | 4.82 | 4.93 | .0018 |
| 120 | tripod01 | 74% | 1 | tripod | seesaw | 6 | 6 | 3.04 | 2.97 | 1.85 | 1.95 | .0029 | .0053 | 4.72 | 4.92 | 0 |
| 121 | cabbage | 74% | 2 | cabbage | uniform | 7 | 7 | 4.07 | 4.16 | 1.65 | 2.00 | .0027 | .0043 | 4.75 | 4.67 | .0053 |
| 122 | raccoon | 74% | 2 | raccoon | notepad | 7 | 7 | 2.57 | 2.55 | 2.45 | 2.80 | .0055 | .0046 | 4.67 | 4.70 | .0004 |
| 123 | dormer | 76% | 1 | window | letter | 6 | 6 | 4.84 | 4.85 | 1.40 | 1.00 | .0106 | .0087 | 4.86 | 4.70 | .0094 |
| 124 | volleyball | 76% | 1 | volleyball | chimpanzee | 10 | 10 | 3.31 | 3.19 | 3.80 | 3.70 | .0050 | .0052 | 4.93 | 4.96 | 0 |
| 125 | cocktailshrimp02 | 76% | 2 | shrimp | tablet | 6 | 6 | 3.63 | 3.54 | 1.80 | 1.65 | .0030 | .0045 | 4.80 | 4.82 | .0042 |
| 126 | bowrake | 76% | 1 | rake | yolk | 4 | 4 | 3.40 | 3.52 | 1.00 | 1.20 | .0035 | .0040 | 4.84 | 4.78 | .0043 |
| 127 | tulip02 | 76% | 1 | tulip | llama | 5 | 5 | 3.21 | 3.12 | 1.70 | 1.60 | .0031 | .0054 | 5.00 | 4.78 | .0022 |
| 128 | tie02 | 79% | 1 | tie | map | 3 | 3 | 4.58 | 4.52 | 1.00 | 1.00 | .0051 | .0031 | 4.81 | 4.93 | .0008 |
| 129 | popcorn | 79% | 1 | popcorn | luggage | 7 | 7 | 3.68 | 3.61 | 2.60 | 2.55 | .0038 | .0017 | 5.00 | 4.83 | 0 |
| 130 | pigeon | 79% | 1 | pigeon | muscle | 6 | 6 | 4.03 | 4.11 | 1.70 | 1.80 | .0048 | .0034 | 4.71 | 4.50 | .0027 |
| 131 | honeybee | 79% | 1 | bee | lid | 3 | 3 | 4.19 | 4.16 | 1.00 | 1.00 | .0063 | .0044 | 4.88 | 4.96 | 0 |
| 132 | callbell | 79% | 2 | bell | oven | 4 | 4 | 4.54 | 4.54 | 1.00 | 1.00 | .0073 | .0079 | 4.96 | 4.97 | .0066 |
| 133 | teapot | 79% | 1 | teapot | mousse | 6 | 6 | 3.78 | 3.76 | 1.90 | 1.35 | .0054 | .0076 | 4.96 | 4.83 | .0039 |

| | Image ID | %Agree | Set | Word | | Length | | Zipf | | OLD20 | | BG | | CNC | | Cosine PPMI |
|---|---|---|---|---|---|---|---|---|---|---|---|---|---|---|---|---|
| | | | | C | I | C | I | C | I | C | I | C | I | C | I | |
| 134 | rope03 | 79% | 1 | rope | text | 4 | 4 | 4.30 | 4.40 | 1.00 | 1.10 | .0040 | .0035 | 4.93 | 4.93 | 0 |
| 135 | marble | 80% | 2 | marble | puppet | 6 | 6 | 3.86 | 3.77 | 1.50 | 1.70 | .0051 | .0027 | 4.85 | 4.64 | .0094 |
| 136 | boot02b | 82% | 2 | boot | page | 4 | 4 | 4.43 | 4.52 | 1.00 | 1.00 | .0043 | .0028 | 4.96 | 4.90 | .0009 |
| 137 | plum01 | 82% | 1 | plum | ramp | 4 | 4 | 3.79 | 3.67 | 1.00 | 1.00 | .0016 | .0030 | 4.85 | 4.69 | .0047 |
| 138 | tampon | 82% | 1 | tampon | poncho | 6 | 6 | 2.30 | 2.41 | 1.80 | 1.65 | .0051 | .0059 | 4.86 | 4.97 | .0076 |
| 139 | slipper01b | 82% | 2 | slipper | warship | 7 | 7 | 3.19 | 3.09 | 1.40 | 1.85 | .0054 | .0056 | 4.86 | 4.86 | 0 |
| 140 | chalk | 82% | 2 | chalk | organ | 5 | 5 | 3.87 | 3.99 | 1.30 | 1.00 | .0077 | .0080 | 4.90 | 4.77 | .0018 |
| 141 | banjo | 83% | 2 | banjo | scalp | 5 | 5 | 3.31 | 3.28 | 1.45 | 1.35 | .0057 | .0040 | 4.90 | 4.82 | .0050 |
| 142 | peanut01 | 83% | 2 | peanut | bumper | 6 | 6 | 3.65 | 3.53 | 1.95 | 1.40 | .0078 | .0058 | 4.89 | 4.96 | .0091 |
| 143 | pillow01a | 84% | 2 | pillow | beetle | 6 | 6 | 3.73 | 3.72 | 1.60 | 1.60 | .0050 | .0053 | 5.00 | 4.83 | .0068 |
| 144 | cigar | 85% | 2 | cigar | stump | 5 | 5 | 3.59 | 3.52 | 1.75 | 1.25 | .0041 | .0037 | 4.93 | 4.78 | 0 |
| 145 | jellyfish | 86% | 2 | jellyfish | sunflower | 9 | 9 | 3.56 | 3.55 | 3.10 | 2.95 | .0049 | .0054 | 4.93 | 4.80 | .0096 |
| 146 | calendar | 86% | 2 | calendar | medicine | 8 | 8 | 4.33 | 4.31 | 2.30 | 2.10 | .0085 | .0084 | 4.62 | 4.79 | .0017 |
| 147 | bull | 86% | 1 | bull | cave | 4 | 4 | 4.28 | 4.19 | 1.00 | 1.00 | .0052 | .0064 | 4.85 | 4.96 | .0058 |
| 148 | daddylonglegs | 86% | 1 | spider | tongue | 6 | 6 | 4.24 | 4.36 | 1.25 | 1.75 | .0059 | .0081 | 4.97 | 4.93 | .0083 |
| 149 | chimney | 86% | 2 | chimney | bicycle | 7 | 7 | 3.90 | 3.92 | 1.85 | 2.40 | .0047 | .0027 | 5.00 | 4.89 | .0098 |
| 150 | ashtray01 | 87% | 2 | ashtray | brownie | 7 | 7 | 3.20 | 3.29 | 2.30 | 1.75 | .0043 | .0033 | 4.97 | 4.82 | .0055 |
| 151 | binoculars01b | 87% | 1 | binoculars | ammunition | 10 | 10 | 3.59 | 3.61 | 3.45 | 3.00 | .0065 | .0057 | 5.00 | 4.88 | .0099 |
| 152 | baseball01a | 87% | 2 | baseball | cinnamon | 8 | 8 | 3.74 | 3.78 | 2.55 | 2.55 | .0057 | .0072 | 4.86 | 4.85 | 0 |
| 153 | broom01 | 87% | 1 | broom | algae | 5 | 5 | 3.56 | 3.44 | 1.15 | 1.60 | .0045 | .0026 | 4.89 | 4.93 | .0051 |

| | Image ID | %Agree | Set | Word | | Length | | Zipf | | OLD20 | | BG | | CNC | | Cosine PPMI |
|---|---|---|---|---|---|---|---|---|---|---|---|---|---|---|---|---|
| | | | | C | I | C | I | C | I | C | I | C | I | C | I | |
| 154 | balloon01b | 87% | 1 | balloon | stomach | 7 | 7 | 4.25 | 4.28 | 1.65 | 1.95 | .0073 | .0072 | 4.92 | 4.89 | .0071 |
| 155 | avocado01 | 87% | 1 | avocado | sparrow | 7 | 7 | 3.25 | 3.38 | 2.55 | 1.80 | .0031 | .0046 | 4.89 | 4.85 | .0025 |
| 156 | sock01a | 87% | 2 | sock | tuna | 4 | 4 | 3.77 | 3.76 | 1.00 | 1.00 | .0029 | .0025 | 4.91 | 4.89 | .0047 |
| 157 | jeans01 | 88% | 1 | jeans | wagon | 5 | 5 | 3.84 | 3.73 | 1.30 | 1.60 | .0078 | .0065 | 5.00 | 4.89 | .0012 |
| 158 | nose | 88% | 2 | nose | mail | 4 | 4 | 4.72 | 4.63 | 1.00 | 1.00 | .0057 | .0042 | 4.89 | 4.69 | .0017 |
| 159 | knee | 88% | 2 | knee | soil | 4 | 4 | 4.26 | 4.35 | 1.35 | 1.00 | .0048 | .0039 | 5.00 | 4.87 | .0075 |
| 160 | stool01 | 88% | 2 | stool | weeds | 5 | 5 | 3.71 | 3.66 | 1.05 | 1.00 | .0078 | .0054 | 4.90 | 4.83 | 0 |
| 161 | jeep | 88% | 1 | jeep | wick | 4 | 4 | 3.17 | 3.16 | 1.00 | 1.00 | .0023 | .0039 | 4.80 | 4.69 | .0004 |
| 162 | cannon | 88% | 2 | cannon | throat | 6 | 6 | 4.08 | 4.16 | 1.15 | 1.70 | .0092 | .0116 | 4.79 | 4.97 | .0022 |
| 163 | ostrich | 88% | 2 | ostrich | shuttle | 7 | 7 | 3.52 | 3.58 | 2.10 | 1.80 | .0053 | .0036 | 4.71 | 4.63 | .0077 |
| 164 | porcupine | 88% | 1 | porcupine | lawnmower | 9 | 9 | 3.06 | 3.11 | 3.25 | 3.45 | .0064 | .0052 | 5.00 | 4.97 | .0023 |
| 165 | arrow02 | 90% | 2 | arrow | jewel | 5 | 5 | 3.78 | 3.76 | 1.00 | 1.75 | .0059 | .0035 | 4.97 | 4.96 | .0006 |
| 166 | tricycle | 90% | 2 | tricycle | songbird | 8 | 8 | 2.73 | 2.75 | 2.60 | 2.80 | .0033 | .0053 | 4.68 | 4.59 | .0062 |
| 167 | sponge01 | 90% | 2 | sponge | timber | 6 | 6 | 4.12 | 4.05 | 1.45 | 1.40 | .0068 | .0075 | 5.00 | 4.90 | .0002 |
| 168 | celery | 92% | 1 | celery | tattoo | 6 | 6 | 3.66 | 3.78 | 1.90 | 1.85 | .0082 | .0067 | 4.80 | 4.71 | .0039 |
| 169 | violin | 92% | 1 | violin | burger | 6 | 6 | 3.82 | 3.90 | 1.75 | 1.15 | .0081 | .0065 | 4.96 | 4.93 | .0014 |
| 170 | iron01b | 92% | 1 | iron | soup | 4 | 4 | 4.52 | 4.41 | 1.00 | 1.00 | .0078 | .0086 | 4.59 | 4.72 | .0060 |
| 171 | lamp04a | 92% | 1 | lamp | wool | 4 | 4 | 4.09 | 4.11 | 1.00 | 1.00 | .0030 | .0036 | 4.97 | 4.86 | .0093 |
| 172 | scarf | 92% | 2 | scarf | patio | 5 | 5 | 3.76 | 3.73 | 1.05 | 1.35 | .0043 | .0058 | 4.97 | 4.89 | .0026 |
| 173 | microscope | 92% | 2 | microscope | spacecraft | 10 | 10 | 3.56 | 3.46 | 2.50 | 3.25 | .0035 | .0025 | 5.00 | 4.80 | .0083 |

| | Image ID | %Agree | Set | Word | | Length | | Zipf | | OLD20 | | BG | | CNC | | Cosine PPMI |
|---|---|---|---|---|---|---|---|---|---|---|---|---|---|---|---|---|
| | | | | C | I | C | I | C | I | C | I | C | I | C | I | |
| 174 | rice | 92% | 1 | rice | bomb | 4 | 4 | 4.42 | 4.49 | 1.00 | 1.00 | .0050 | .0031 | 4.86 | 4.84 | .0075 |
| 175 | rooster | 93% | 1 | rooster | serpent | 7 | 7 | 3.13 | 3.16 | 1.50 | 1.70 | .0087 | .0088 | 4.75 | 4.97 | .0008 |
| 176 | beaver | 93% | 1 | beaver | shrine | 6 | 6 | 3.50 | 3.53 | 1.00 | 1.55 | .0098 | .0088 | 4.68 | 4.47 | .0097 |
| 177 | trophy01 | 93% | 2 | trophy | jacket | 6 | 6 | 4.37 | 4.29 | 1.90 | 1.40 | .0025 | .0032 | 4.89 | 4.86 | .0032 |
| 178 | cactus | 93% | 2 | cactus | poodle | 6 | 6 | 3.35 | 3.27 | 1.70 | 1.45 | .0037 | .0035 | 5.00 | 4.89 | 0 |
| 179 | snowboard | 95% | 2 | snowboard | amplifier | 9 | 9 | 2.84 | 2.73 | 2.65 | 2.65 | .0035 | .0051 | 4.86 | 4.79 | .0076 |
| 180 | potato02b | 95% | 1 | potato | ticket | 6 | 6 | 4.44 | 4.51 | 1.60 | 1.35 | .0071 | .0048 | 4.85 | 4.70 | .0086 |
| 181 | apple07 | 95% | 1 | apple | penny | 5 | 5 | 4.58 | 4.49 | 1.40 | 1.00 | .0034 | .0044 | 5.00 | 4.83 | .0080 |
| 182 | apron | 95% | 2 | apron | lager | 5 | 5 | 3.48 | 3.56 | 1.05 | 1.00 | .0062 | .0075 | 4.87 | 4.64 | .0001 |
| 183 | cigarette | 95% | 2 | cigarette | porcelain | 9 | 9 | 4.11 | 4.10 | 2.80 | 2.90 | .0065 | .0071 | 4.88 | 4.63 | .0091 |
| 184 | skunk | 95% | 1 | skunk | quail | 5 | 5 | 3.36 | 3.48 | 1.55 | 1.45 | .0016 | .0024 | 4.88 | 4.65 | .0054 |
| 185 | barnowl | 95% | 2 | owl | jug | 3 | 3 | 4.07 | 4.06 | 1.00 | 1.00 | .0026 | .0016 | 4.93 | 4.96 | .0095 |
| 186 | lipstick02a | 95% | 1 | lipstick | cardigan | 8 | 8 | 3.62 | 3.50 | 2.30 | 1.90 | .0047 | .0064 | 4.90 | 4.96 | .0096 |
| 187 | brick | 95% | 1 | brick | robot | 5 | 5 | 4.18 | 4.09 | 1.00 | 1.60 | .0036 | .0040 | 4.83 | 4.65 | .0017 |
| 188 | leaf02a | 97% | 2 | leaf | pork | 4 | 4 | 4.29 | 4.39 | 1.00 | 1.00 | .0059 | .0048 | 5.00 | 4.79 | 0 |
| 189 | carrot01 | 97% | 2 | carrot | tissue | 6 | 6 | 4.08 | 3.97 | 1.40 | 1.75 | .0059 | .0051 | 5.00 | 4.93 | .0053 |
| 190 | kite | 98% | 2 | kite | cart | 4 | 4 | 3.89 | 3.77 | 1.00 | 1.00 | .0084 | .0064 | 5.00 | 4.89 | .0004 |
| 191 | locker | 98% | 1 | locker | manual | 6 | 6 | 3.60 | 3.70 | 1.00 | 1.75 | .0064 | .0069 | 4.67 | 4.45 | .0048 |
| 192 | pumpkin | 98% | 2 | pumpkin | trolley | 7 | 7 | 3.79 | 3.82 | 1.70 | 1.70 | .0052 | .0055 | 4.90 | 4.73 | .0025 |
| 193 | zebra | 98% | 2 | zebra | snail | 5 | 5 | 3.69 | 3.69 | 1.80 | 1.45 | .0016 | .0026 | 4.86 | 4.93 | .0062 |

| | Image ID | %Agree | Set | Word | | Length | | Zipf | | OLD20 | | BG | | CNC | | Cosine PPMI |
|---|---|---|---|---|---|---|---|---|---|---|---|---|---|---|---|---|
| | | | | C | I | C | I | C | I | C | I | C | I | C | I | |
| 194 | kangaroo | 98% | 1 | kangaroo | lemonade | 8 | 8 | 3.62 | 3.57 | 2.75 | 2.70 | .0077 | .0055 | 4.86 | 4.83 | .0058 |
| 195 | squirrel | 100% | 1 | squirrel | passport | 8 | 8 | 3.94 | 4.01 | 2.10 | 2.25 | .0045 | .0045 | 4.89 | 5.00 | .0051 |
| 196 | mushroom01 | 100% | 2 | mushroom | carriage | 8 | 8 | 3.87 | 3.98 | 2.60 | 1.90 | .0040 | .0044 | 4.83 | 4.86 | .0003 |
| 197 | pear01 | 100% | 1 | pear | lung | 4 | 4 | 3.83 | 3.81 | 1.00 | 1.00 | .0078 | .0055 | 4.93 | 4.82 | .0050 |
| 198 | snowman | 100% | 1 | snowman | pancake | 7 | 7 | 3.52 | 3.48 | 1.90 | 2.05 | .0060 | .0059 | 4.64 | 4.86 | .0054 |
| 199 | onion | 100% | 1 | onion | torch | 5 | 5 | 4.28 | 4.21 | 1.70 | 1.30 | .0086 | .0073 | 4.86 | 4.76 | .0013 |
| 200 | toothbrush03b | 100% | 2 | toothbrush | cheesecake | 10 | 10 | 3.48 | 3.54 | 3.80 | 3.45 | .0084 | .0085 | 5.00 | 4.97 | .0040 |

#### B Behavioural Validation Results

To validate the stimulus generation method for the picture-word stimuli, we ran a behavioural experiment using a stimulus set generated from a very similar pipeline to that described in the manuscript. The only differences in the pipeline were that (a) Zipf frequency was controlled within $\pm.2$, (b) Levenshtein distance was not maximised, (c) OLD20 was not controlled for, and (d) the split into stimulus Sets 1 and 2 was optimised from only 20,000 iterations. The stimuli generated for the validation experiment varied in predictability from 12 to 100%. The procedure was also identical to that described in the Procedure section of the manuscript, except that participants could respond as soon as the word was presented, rather than 1 second after presentation, and the word did not change colour. Participants comprised 35 monolingual native English speakers (15 female, 19 male, 1 non-binary) who were not diagnosed with any reading disorder. Age varied from 18 to 26 years (M=21.4, *SD*=2.05), and all participants reported being right-handed with normal or corrected-to-normal vision. Trials were excluded if response times (RTs) were less than 250 ms or more than 2000 ms. The logic for the validation experiment was as follows: assuming the stimulus pipeline produces suitably controlled stimuli, increased predictability should facilitate task performance for congruent trials and have either no effect or a minimal effect on performance for incongruent trials.

We modelled the RT data with a shifted log-normal distribution. This allowed us to describe changes in the means ($\mu$) and standard deviations ($\sigma$) of log-transformed RTs, while also modelling changes in shift ($\delta$). To model the validation experiment data, we fit a Bayesian mixed-effects model estimating the same fixed and maximal random effects structure for each parameter ($\mu$, $\sigma$, $\delta$) of the shifted log-normal distribution. This was achieved using the *brms* package for R (Bürkner, 2017), a high-level interface for STAN (STAN Development Team, 2023). This model estimated the plausibility of population values for each parameter of the shifted log-normal distribution as a function of the maximal hierarchical structure justified by the experiment's design. The parameter of $\mu$

was modelled with an identity link function, while $\sigma$ and $\delta$ were modelled with log link functions. The same predictors and random effects structure were used for each parameter as described for the EEG experiment, though with a key difference being that predictability was normalised between 12% and 100% rather between than 7% and 100%, due to different minima in the experiments' stimuli. The full formula, in *brms* syntax, was specified as:

```
rt ~ 1 + congruency * predictability +
        (1 + congruency * predictability | subject_id) +
        (1 + congruency | image_id) +
        (1 | word_id),
sigma ~ 1 + congruency * predictability +
        (1 + congruency * predictability | subject_id) +
        (1 + congruency | image_id) +
        (1 | word_id),
ndt ~ 1 + congruency * predictability +
        (1 + congruency * predictability | subject_id) +
        (1 + congruency | image_id) +
        (1 | word_id)
```

Prior distributions were specified to be broad enough as to be uninformative but constrained to cover plausible values for response time distributions for a cognitive task (Figure 2). Fixed effects' slopes' prior distributions were drawn from $N(0, 2.5)$ and fixed effects' intercepts' prior distributions from $N(0, 7.5)$. The prior distributions for the standard deviations of random effects were specified as student's $t$ distributions centred on zero, with 3 degrees of freedom and a scale parameter of 2. The model was fit with 5 Markov chains, each with 25,000 iterations (17,500 warm-up and 7,500 sampling). The *adapt_delta* parameter was set to .99. The densities of the posterior distributions, relative to those of the priors, are shown in Figure 2.

The results from the shifted log-normal model showed the expected effects, with predictability leading to faster responses for congruent trials, but having weak effects on incongruent trials (Figure 3). It also demonstrated that when predictability is low, response times show similar central tendency for congruent and incongruent trials though a larger spread in the distribution for congruent trials. When predictability is high, on the

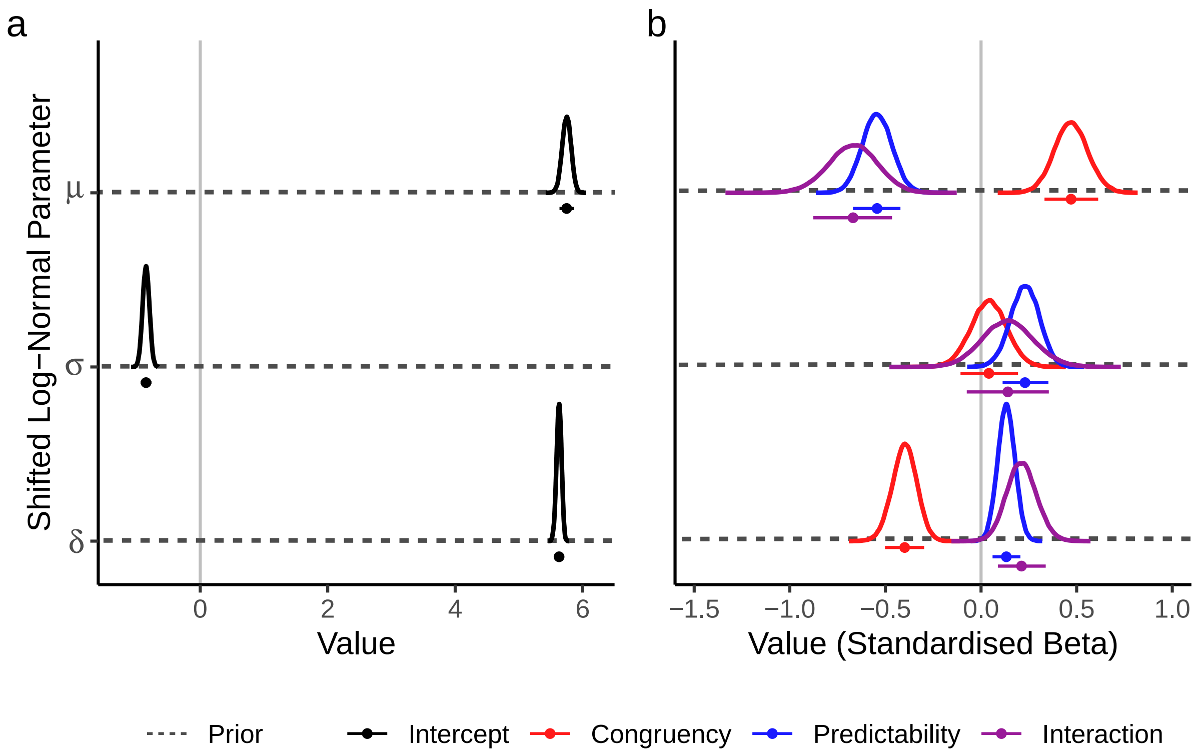

**Figure 2**

*Prior (dashed lines) and posterior distributions (solid lines) for all fixed effects estimated in the Bayesian shifted log-normal model, for the three distribution parameters. Panels show distributions for (**a**) intercept parameters, and (**b**) slope parameters. For both panels, points below distributions' densities depict median posterior estimates, while the whiskers show the extents of 89% highest density intervals (HDIs).*

other hand, the difference is mostly due to changes in shift, whereas other features of the distribution are very similar.

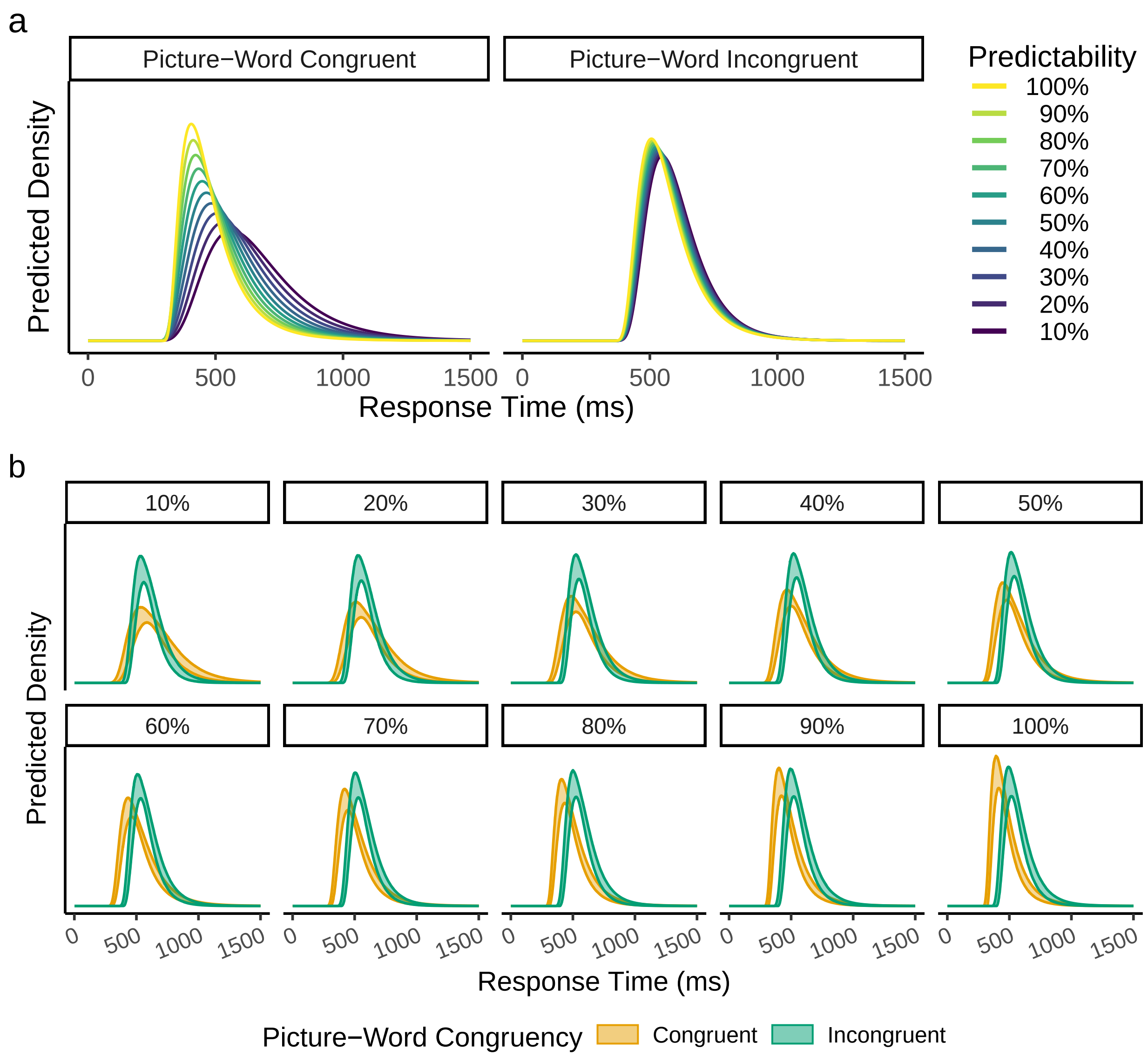

**Figure 3**

*Fixed effect predictions of RT distributions in the behavioural validation experiment for the picture-word stimuli. Predictions of RT distributions, derived from the shifted log-normal model, are shown for congruent and incongruent trials for values of percentage of name agreement, from 10 to 100% in steps of 10. The two panels show the same results but highlight (**a**) the effect of predictability for picture-congruent and picture-incongruent words, and (**b**) the effect of picture-word congruency at different values of predictability, showing the degree of certainty in the predictions with the 89% HDIs of the predictions from all posterior samples. Density is scaled consistently across panels.*

#### C Localiser Task Stimuli

To generate the localiser stimuli, a large list of suitable words ($N$=27,332) was identified by subsetting the word prevalence norms of Brysbaert et al. (2019) to only contain words known by at least 90% of participants and which were not selected for the main experiment. A representative sample ($N$=100) of this list was generated by maximising distributional overlap (Pastore & Calcagnì, 2019), between the sample and the full list of candidates, on 13 variables where observations were available: (1) word prevalence (Brysbaert et al., 2019); (2) length (number of characters); (3) word frequency in Zipf in SUBTLEX-UK (van Heuven et al., 2014); (4) part of speech according to SUBTLEX-UK; (5) character bigram probability calculated from SUBTLEX-UK; (6) OLD20 (Yarkoni et al., 2008) calculated from the LexOPS dataset (Taylor et al., 2020); (7) concreteness (Brysbaert et al., 2014); (8) age of acquisition (Kuperman et al., 2012); (9, 10) average lexical decision response time (RT) and accuracy according to the British Lexicon Project (Keuleers et al., 2012); and (11, 12, 13) the emotion ratings of valence, arousal, and dominance (Warriner et al., 2013). Similarity in the categorical variable of part of speech was maximised with dummy-coded variables (0 or 1 for absence or presence of a category, respectively). Distributional similarity across all variables was maximised by selecting from 500,000 random samples the sample with the highest total distributional overlap with the full list of possible words. Distributions of the selected sample of words are summarised in **Figure 4**. The full list of stimuli for the localiser task is presented in **Table 2**.

The false-font strings consisted of characters from the Brussels Artificial Character Set (BACS; Vidal et al., 2017) in BACS2serif font. In this way, we had an item-wise false-font match to each word, where every Courier New character in the word stimuli is replaced with a BACS character matched in the number of strokes, junctions, terminations, and serifs. The phase-shuffled stimuli were generated by using a Fourier transformation to extract the phase and amplitude from the word images. Phase values were randomly shuffled (i.e., permuted), such that the overall distribution of phase could be preserved,

**Figure 4**

*Distributions of key variables illustrate the similarity between the selected localiser stimuli words (sample) and the list of words from which they were drawn (population).*

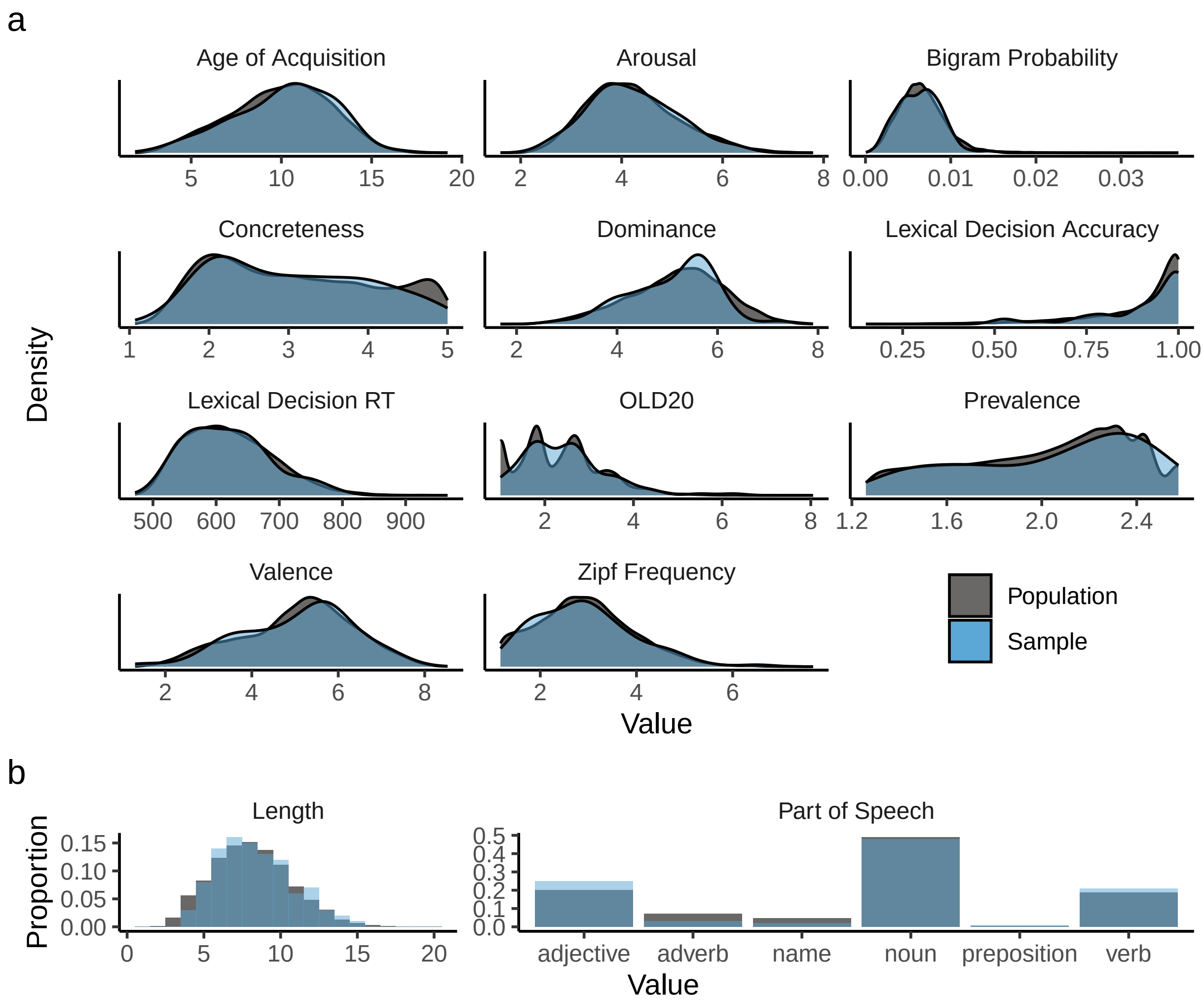

Panel **a** shows distributional similarity of continuous variables. Panel **b** shows similarity in length (all integer values) as a histogram showing proportions, and the similarity in the counts of each part of speech category as a bar plot of proportions. Only the part of speech categories which were present in the sample are shown. No members of less common part of speech categories, such as determiner or number, were selected in the sample.

while amplitude values were unchanged. An inverse Fourier transformation was then used to generate a new image with the original amplitude values, but with phase randomly shuffled. To prevent phase shuffling from producing noticeably large changes in contrast, the phase shuffling was done on a version of the word image with 50% of the original contrast. After the inverse Fourier transformation, the contrast of the generated phase-shuffled image was readjusted to equal that of the original word image. To avoid repeating the same stimuli across participants more than necessary, unique phase-shuffled images were generated for each trial, for each participant.

Versions of the localiser task's stimuli were also created in green, to signal the participant to respond. For words and nonwords, this was done by simply changing the font colour to green. To preserve image intensity, the colour of phase-shuffled images was changed by altering pixels in the following way. For pixels in which the value in the green channel was less than 50% of the maximum intensity (i.e., the intensity of all channels in the grey background), values in red and blue channels were altered to equal the value in the green channel for that pixel. For all other pixels, the values in red and blue channels were set to 50% of the maximum intensity.

**Table 2**

*All word stimuli for the localiser task, and associated values on variables that were matched distribution-wise. False-font strings and phase-shuffled images are not presented here; false-font strings were just the words in BACS2serif font, while a unique phase-shuffled image was generated for each trial. The columns are as follows:* Word *= words presented in the task;* Length *= number of characters;* Zipf *= Zipf frequency in SUBTLEX-UK;* PREV *= word prevalence values in Brysbaert et al. (2019);* OLD20 *= OLD20 values in the LexOPS dataset;* BG *= mean character bigram probabilities in SUBTLEX-UK;* PoS *= dominant part of speech in SUBTLEX-UK;* CNC *= mean concreteness ratings in Brysbaert et al. (2014);* AoA *= mean age of acquisition ratings in Kuperman et al. (2012);* VAL, AROU, *and* DOM *= mean valence, arousal, and dominance ratings, respectively, from Warriner et al. (2013);* LDT RT *and* LDT Acc *= average response times (in ms) and accuracies in lexical decision, from the BLP. Rows are numbered for ease of reference.*

| | Word | Length | Zipf | PREV | OLD20 | BG | PoS | CNC | AoA | VAL | AROU | DOM | LDT RT | LDT Acc |
|---|---|---|---|---|---|---|---|---|---|---|---|---|---|---|
| 1 | tracker | 7 | 3.12 | 2.58 | 1.45 | .0062 | noun | 3.89 | 9.61 | 4.87 | 4.59 | 5.00 | 583.21 | .98 |
| 2 | tablespoonful | 13 | 1.97 | 1.40 | 5.30 | .0045 | adjective | 4.24 | 7.58 | - | - | - | - | - |
| 3 | curricular | 10 | 2.50 | 1.61 | 2.85 | .0042 | adjective | 2.77 | 10.10 | - | - | - | - | - |
| 4 | sheathed | 8 | 1.74 | 1.45 | 1.95 | .0195 | verb | 3.04 | - | - | - | - | 699.78 | .68 |
| 5 | wasabi | 6 | 2.74 | 1.74 | 2.00 | .0041 | noun | 4.67 | 13.95 | - | - | - | - | - |
| 6 | persecute | 9 | 2.43 | 1.80 | 2.65 | .0068 | verb | 2.53 | 10.06 | 3.11 | 5.11 | 4.09 | - | - |
| 7 | enlarge | 7 | 2.70 | 2.16 | 2.35 | .0053 | verb | 3.17 | 8.26 | 5.33 | 3.87 | 5.89 | 568.70 | .95 |
| 8 | harvester | 9 | 3.15 | 2.12 | 2.60 | .0107 | noun | 4.21 | 9.53 | - | - | - | - | - |
| 9 | campaign | 8 | 4.90 | 2.44 | 2.20 | .0027 | noun | 3.00 | 12.55 | 4.55 | 3.50 | 5.14 | 561.37 | .98 |
| 10 | menacingly | 10 | 2.20 | 1.79 | 3.25 | .0078 | adverb | 1.93 | - | - | - | - | - | - |
| 11 | footwork | 8 | 3.40 | 2.13 | 2.15 | .0044 | noun | 3.32 | 10.63 | 5.74 | 3.96 | 5.58 | 680.59 | .88 |
| 12 | respective | 10 | 3.20 | 2.10 | 2.65 | .0067 | adjective | 1.79 | 10.78 | 5.90 | 3.76 | 6.42 | - | - |
| 13 | layperson | 9 | 1.65 | 1.35 | 2.85 | .0068 | noun | 3.44 | 13.74 | - | - | - | - | - |

| | Word | Length | Zipf | PREV | OLD20 | BG | PoS | CNC | AoA | VAL | AROU | DOM | LDT RT | LDT Acc |
|---|---|---|---|---|---|---|---|---|---|---|---|---|---|---|
| 14 | microcomputer | 13 | 1.30 | 1.82 | 4.45 | .0055 | noun | 4.55 | 13.89 | - | - | - | - | - |
| 15 | flatterer | 9 | 2.32 | 1.32 | 1.85 | .0104 | noun | 2.89 | 12.44 | - | - | - | - | - |
| 16 | chilled | 7 | 3.63 | 2.35 | 1.75 | .0074 | verb | 3.22 | - | - | - | - | 566.50 | 1.00 |
| 17 | blackheads | 10 | 1.93 | 2.07 | 2.35 | .0065 | noun | 4.79 | - | - | - | - | 742.83 | .97 |
| 18 | fortunate | 9 | 4.06 | 2.24 | 2.50 | .0056 | adjective | 2.04 | 10.17 | 7.33 | 3.81 | 5.83 | 635.46 | .95 |
| 19 | screeching | 10 | 2.81 | 2.24 | 2.55 | .0092 | verb | 3.71 | - | - | - | - | 621.72 | .93 |
| 20 | chimp | 5 | 3.42 | 2.23 | 1.35 | .0048 | noun | 4.96 | 7.17 | 6.00 | 3.80 | 4.95 | 605.63 | .88 |
| 21 | payroll | 7 | 3.10 | 2.43 | 2.40 | .0042 | noun | 3.70 | 12.79 | 6.19 | 3.82 | 5.11 | 632.25 | .97 |
| 22 | seer | 4 | 2.48 | 1.26 | 1.00 | .0110 | noun | - | 10.56 | 5.35 | 3.77 | 5.41 | 752.86 | .53 |
| 23 | coexist | 7 | 2.00 | 1.99 | 2.45 | .0053 | verb | 2.25 | 11.56 | 5.95 | 3.48 | 5.92 | - | - |
| 24 | smelly | 6 | 3.87 | 2.43 | 1.45 | .0057 | adjective | 3.07 | 4.32 | 2.68 | 5.43 | 4.00 | 533.24 | 1.00 |
| 25 | discouraging | 12 | 2.69 | 2.33 | 3.25 | .0084 | verb | 1.83 | 9.11 | 2.89 | 4.17 | 4.22 | - | - |
| 26 | exotic | 6 | 4.10 | 2.43 | 1.85 | .0039 | adjective | 2.11 | 10.42 | 7.55 | 6.90 | 5.65 | - | - |
| 27 | snow | 4 | 4.79 | 2.33 | 1.00 | .0040 | noun | 4.85 | 4.11 | 6.78 | 4.57 | 5.62 | 506.10 | 1.00 |
| 28 | takeoff | 7 | 2.84 | 1.92 | 2.45 | .0035 | noun | 3.41 | 7.35 | 5.50 | 3.77 | 5.11 | - | - |
| 29 | milkman | 7 | 3.08 | 1.98 | 1.90 | .0054 | noun | 4.61 | 6.37 | 5.75 | 2.73 | 5.54 | 626.19 | 1.00 |
| 30 | intelligent | 11 | 4.09 | 2.58 | 3.15 | .0094 | adjective | 2.46 | 8.28 | 7.60 | 5.67 | 6.77 | - | - |
| 31 | creak | 5 | 2.69 | 1.40 | 1.30 | .0078 | verb | 3.61 | 8.10 | 4.68 | 4.40 | 4.61 | 599.59 | .85 |
| 32 | punchy | 6 | 3.03 | 1.51 | 1.55 | .0024 | adjective | 2.21 | 13.18 | 4.78 | 4.32 | 3.96 | 657.00 | .76 |
| 33 | glutinous | 9 | 2.09 | 1.53 | 2.70 | .0089 | adjective | 2.62 | 14.32 | - | - | - | - | - |
| 34 | monsieur | 8 | 3.70 | 1.35 | 2.75 | .0046 | noun | 3.54 | 10.12 | 5.50 | 3.30 | 5.89 | - | - |
| 35 | sympathetic | 11 | 3.70 | 2.58 | 3.50 | .0105 | adjective | 1.77 | 9.39 | 6.67 | 3.29 | 6.30 | - | - |

| | Word | Length | Zipf | PREV | OLD20 | BG | PoS | CNC | AoA | VAL | AROU | DOM | LDT RT | LDT Acc |
|---|---|---|---|---|---|---|---|---|---|---|---|---|---|---|
| 36 | neurotoxin | 10 | 1.95 | 1.72 | 3.10 | .0071 | noun | 3.12 | 13.58 | - | - | - | - | - |
| 37 | singular | 8 | 3.00 | 2.27 | 2.45 | .0086 | adjective | 2.21 | 9.80 | 4.89 | 3.12 | 5.24 | - | - |
| 38 | snip | 4 | 3.64 | 2.00 | 1.00 | .0012 | noun | 3.68 | 7.24 | 4.32 | 4.74 | 4.95 | 569.42 | .95 |
| 39 | bewildered | 10 | 3.14 | 2.43 | 3.30 | .0080 | verb | 1.80 | 11.63 | 4.32 | 4.57 | 4.42 | - | - |
| 40 | devote | 6 | 3.16 | 2.03 | 1.55 | .0045 | verb | 2.00 | 9.58 | 5.53 | 4.05 | 7.05 | 600.51 | .97 |
| 41 | handily | 7 | 2.30 | 1.62 | 1.90 | .0101 | adverb | 2.08 | - | - | - | - | - | - |
| 42 | orally | 6 | 2.41 | 2.23 | 1.90 | .0076 | adverb | 3.00 | - | - | - | - | - | - |
| 43 | prerecorded | 11 | 1.60 | 2.10 | 3.45 | .0096 | verb | 2.58 | 10.22 | - | - | - | - | - |
| 44 | yodel | 5 | 3.11 | 1.49 | 1.55 | .0054 | name | 4.20 | 8.16 | 6.10 | 3.33 | 5.90 | 703.75 | .50 |
| 45 | impertinently | 13 | 1.17 | 1.38 | 3.90 | .0082 | adverb | - | - | - | - | - | - | - |
| 46 | vacation | 8 | 3.36 | 2.58 | 1.85 | .0063 | noun | 3.14 | 5.22 | 8.53 | 5.22 | 7.11 | - | - |
| 47 | extravagance | 12 | 2.86 | 2.20 | 3.85 | .0038 | noun | 1.73 | 10.74 | 5.74 | 5.40 | 5.79 | - | - |
| 48 | thud | 4 | 3.01 | 2.26 | 1.00 | .0139 | noun | 3.20 | 8.06 | 4.24 | 5.05 | 4.52 | 582.36 | .83 |
| 49 | forewarn | 8 | 1.74 | 1.90 | 2.10 | .0076 | verb | 2.20 | 11.16 | - | - | - | 703.91 | .66 |
| 50 | fatherhood | 10 | 2.73 | 2.44 | 3.20 | .0130 | noun | 2.76 | 8.50 | 6.77 | 4.57 | 5.61 | - | - |
| 51 | correlate | 9 | 2.20 | 2.04 | 2.60 | .0083 | verb | 1.63 | 13.35 | - | - | - | - | - |
| 52 | watercraft | 10 | 1.54 | 1.61 | 2.90 | .0056 | noun | - | - | - | - | - | - | - |
| 53 | sunk | 4 | 3.73 | 2.43 | 1.00 | .0026 | verb | 3.46 | - | - | - | - | 611.78 | .93 |
| 54 | flawlessness | 12 | 1.39 | 1.58 | 3.30 | .0042 | noun | 2.16 | - | - | - | - | - | - |
| 55 | tranquilizer | 12 | 1.47 | 2.02 | 2.30 | .0054 | noun | 4.55 | 11.58 | 4.86 | 3.12 | 4.85 | - | - |
| 56 | pituitary | 9 | 2.47 | 1.52 | 3.70 | .0065 | adjective | 3.33 | 13.06 | 4.79 | 4.40 | 4.91 | - | - |
| 57 | courtside | 9 | 1.81 | 2.00 | 2.85 | .0059 | noun | 3.65 | 12.32 | 6.00 | 4.24 | 6.00 | - | - |

| | Word | Length | Zipf | PREV | OLD20 | BG | PoS | CNC | AoA | VAL | AROU | DOM | LDT RT | LDT Acc |
|---|---|---|---|---|---|---|---|---|---|---|---|---|---|---|
| 58 | wicked | 6 | 4.16 | 2.33 | 1.15 | .0047 | adjective | 2.11 | 8.33 | 2.63 | 5.86 | 3.61 | 579.31 | .93 |
| 59 | regard | 6 | 4.19 | 2.24 | 1.55 | .0067 | noun | 1.79 | 10.20 | 5.70 | 3.39 | 6.38 | 545.31 | .98 |
| 60 | infidelity | 10 | 2.71 | 2.33 | 3.55 | .0072 | noun | 2.07 | 13.89 | 2.10 | 5.70 | 3.86 | - | - |
| 61 | bumping | 7 | 3.29 | 2.34 | 1.55 | .0074 | verb | 4.00 | - | - | - | - | 660.94 | .97 |
| 62 | cannibal | 8 | 2.60 | 2.31 | 2.45 | .0058 | adjective | 3.82 | 9.11 | 2.90 | 6.10 | 3.20 | - | - |
| 63 | texting | 7 | 3.51 | 2.58 | 1.80 | .0093 | verb | 4.23 | - | - | - | - | - | - |
| 64 | apache | 6 | 3.15 | 1.75 | 1.75 | .0091 | name | 3.88 | 10.50 | 5.20 | 3.70 | 4.95 | 747.23 | .68 |
| 65 | generational | 12 | 2.98 | 1.88 | 2.90 | .0084 | adjective | 1.96 | 12.68 | - | - | - | - | - |
| 66 | squint | 6 | 2.79 | 2.33 | 1.75 | .0075 | noun | 4.30 | 8.05 | 4.40 | 3.71 | 4.62 | 586.76 | 1.00 |
| 67 | torture | 7 | 4.00 | 2.43 | 1.80 | .0089 | verb | 3.59 | 10.70 | 1.40 | 5.09 | 2.76 | 530.51 | 1.00 |
| 68 | shattering | 10 | 3.10 | 2.32 | 1.75 | .0115 | verb | 3.43 | 8.00 | 3.67 | 5.00 | 4.63 | - | - |
| 69 | freckled | 8 | 1.30 | 2.43 | 1.90 | .0061 | adjective | 3.86 | 6.58 | - | - | - | 645.19 | .98 |
| 70 | perversion | 10 | 2.35 | 2.07 | 2.70 | .0087 | noun | 2.04 | 13.11 | 3.55 | 5.48 | 3.85 | - | - |
| 71 | shag | 4 | 3.37 | 2.00 | 1.00 | .0073 | noun | 3.15 | 10.53 | 5.38 | 4.95 | 4.86 | 546.18 | .98 |
| 72 | stifle | 6 | 2.68 | 1.97 | 1.70 | .0058 | verb | 2.59 | 10.26 | - | - | - | 659.20 | .82 |
| 73 | syllable | 8 | 2.89 | 2.25 | 2.00 | .0038 | adjective | 3.26 | 8.10 | 4.95 | 2.50 | 5.70 | - | - |
| 74 | ionic | 5 | 2.50 | 1.79 | 1.40 | .0063 | adjective | 2.14 | 14.19 | - | - | - | - | - |
| 75 | explicable | 10 | 1.65 | 2.20 | 2.65 | .0037 | adjective | 1.58 | 12.25 | - | - | - | - | - |
| 76 | dashboard | 9 | 3.06 | 2.33 | 2.65 | .0038 | noun | 4.61 | 9.21 | 5.25 | 3.15 | 5.32 | 651.98 | 1.00 |
| 77 | concessionary | 13 | 2.78 | 1.37 | 3.25 | .0064 | adjective | 2.15 | 14.43 | - | - | - | - | - |
| 78 | retort | 6 | 2.40 | 2.03 | 1.80 | .0103 | noun | 2.75 | 11.50 | - | - | - | 628.15 | .87 |
| 79 | extent | 6 | 4.40 | 2.34 | 1.70 | .0063 | noun | 1.44 | 10.72 | 5.57 | 3.68 | 5.00 | 573.03 | .97 |

| | Word | Length | Zipf | PREV | OLD20 | BG | PoS | CNC | AoA | VAL | AROU | DOM | LDT RT | LDT Acc |
|---|---|---|---|---|---|---|---|---|---|---|---|---|---|---|
| 80 | mutual | 6 | 3.72 | 2.14 | 1.85 | .0038 | adjective | 2.21 | 8.90 | 6.48 | 3.50 | 6.45 | 598.86 | .95 |
| 81 | problematic | 11 | 3.43 | 2.32 | 3.15 | .0050 | adjective | 2.11 | 11.63 | 2.58 | 4.80 | 4.65 | - | - |
| 82 | shiftless | 9 | 1.30 | 1.62 | 2.40 | .0047 | adjective | 2.27 | 12.12 | - | - | - | 693.12 | .70 |
| 83 | pleasantness | 12 | 1.47 | 1.59 | 3.55 | .0072 | noun | 2.00 | 8.44 | - | - | - | - | - |
| 84 | nonpayment | 10 | 1.17 | 1.71 | 3.60 | .0064 | noun | 2.83 | 10.00 | - | - | - | - | - |
| 85 | context | 7 | 4.28 | 2.24 | 1.85 | .0068 | noun | 2.17 | 10.00 | 5.00 | 3.18 | 5.60 | 597.95 | .98 |
| 86 | shifting | 8 | 3.73 | 2.34 | 1.65 | .0088 | verb | 2.86 | - | - | - | - | 605.50 | 1.00 |
| 87 | creamer | 7 | 2.65 | 1.92 | 1.45 | .0101 | noun | 4.66 | 8.72 | 5.47 | 2.81 | 6.09 | 738.38 | .88 |
| 88 | felicity | 8 | 3.44 | 1.49 | 2.10 | .0052 | name | 1.56 | - | - | - | - | - | - |
| 89 | deferred | 8 | 2.97 | 2.05 | 1.75 | .0080 | verb | 2.00 | - | - | - | - | 666.58 | .95 |
| 90 | gyroscope | 9 | 2.19 | 1.67 | 2.75 | .0028 | noun | 4.25 | 12.69 | - | - | - | - | - |
| 91 | recalculate | 11 | 1.81 | 2.15 | 2.95 | .0064 | verb | 2.93 | 11.53 | - | - | - | - | - |
| 92 | frosty | 6 | 3.51 | 2.35 | 1.80 | .0046 | adjective | 3.90 | 6.33 | 6.15 | 4.61 | 5.00 | 607.38 | .98 |
| 93 | cohesiveness | 12 | 1.60 | 1.85 | 3.85 | .0088 | noun | 2.62 | - | - | - | - | - | - |
| 94 | meld | 4 | 2.19 | 1.34 | 1.00 | .0060 | verb | 2.86 | 11.63 | - | - | - | 601.62 | .34 |
| 95 | awfulness | 9 | 2.37 | 1.67 | 2.80 | .0031 | noun | 2.20 | 9.67 | - | - | - | - | - |
| 96 | rolled | 6 | 4.16 | 2.25 | 1.45 | .0069 | verb | 3.64 | - | - | - | - | 546.38 | .97 |
| 97 | orange | 6 | 4.64 | 2.26 | 1.40 | .0101 | noun | 4.66 | 3.26 | 6.81 | 4.04 | 5.58 | 519.53 | .98 |
| 98 | easily | 6 | 4.69 | 2.43 | 1.75 | .0061 | adverb | 1.80 | - | - | - | - | - | - |
| 99 | reestablish | 11 | 1.70 | 1.67 | 3.40 | .0077 | verb | 2.54 | 10.33 | 6.14 | 4.00 | 6.18 | - | - |
| 100 | lacquer | 7 | 3.06 | 1.56 | 1.85 | .0050 | noun | 4.28 | 13.19 | 4.95 | 3.30 | 5.00 | 699.11 | .75 |

#### D Statistical Power Analysis

We conducted simulations to identify the number of participants required to reach at least 80% power (an arbitrary but commonly used target for statistical power), if we were to carry out the same experiment a large number of times. To match our hypothesis, the planned analysis for this experiment focused on the Congruency-Predictability interaction. A fixed effect coefficient for the interaction in the expected direction would be evidence for a Congruency-dependent effect of Predictability on the N1 that is consistent with a simple predictive coding account. The expected fixed effects coefficients were calculated assuming an interaction between Predictability and image-word Congruency consisting of a .75 $\mu$V reduction in N1 amplitude for the most relative to the least predictable congruent trials, with no difference for incongruent trials. Importantly, while we simulated a pattern of effects in which predictability reduced N1 amplitude for picture-congruent words, but not -incongruent words, the interaction term would capture any pattern of results consistent with our predictive coding hypothesis.

To determine the .75 $\mu$V effect size, first we decided to simulate the difference as a proportion of the maximum N1 amplitude, because different EEG systems and setups can result in vastly different voltage measurements. Next, to identify a realistic proportional difference at the maximum level of predictability (100% name agreement) between picture-congruent and picture-incongruent words, we considered the design by Kim and Gilley (2013), which is as close to this design as we could find. In their study, 53 participants were presented with highly predictable target words which were either prediction-congruent or prediction-incongruent. Kim and Gilley observed left-lateralised occipitotemporal electrodes' N1 peaks that were less negative when the word was prediction-congruent (-2.6 $\mu$V) than when the word was prediction-incongruent (-3.9 $\mu$V), equal to a proportional difference of .33. A less comparable, though still possibly informative, study from Kim and Lai (2012) presented 20 participants with 180 high Cloze probability sentences (with 550 ms SOAs such that overlap of ERPs was minimised). The last word in each sentence was either a highly predictable word, an orthographically similar pseudoword, an orthographically dissimilar pseudoword, or a consonant string nonword. Here, the N1 (170-205 ms) for a left occipitotemporal electrode was shown to be more negative for nonwords and orthographically dissimilar pseudowords (both around -4 $\mu$V) than for the predicted word and an orthographically similar pseudoword (both around -3 $\mu$V). This is equal to a proportional difference of .25. We decided that other potentially comparable studies, published at the time of the power analysis, were too different in their experimental design, either because they used manipulations other than biasing predictions for specific word forms (Chen et al., 2013, 2015; Segalowitz & Zheng, 2009; Strijkers et al., 2015; Walsh et al., 2020; Wang & Maurer, 2017) or they presented the target items midway through sentences using an SOA of 300 ms or less resulting in overlapping ERPs (Dambacher et al., 2012; Kretzschmar et al., 2015; Sereno et al., 2019).

Given the lack of relevant data, we decided a proportional difference of .15 was a realistic effect size for the difference between picture-congruent and picture-incongruent trials at the maximum level of predictability. In previous participants recorded on the same EEG system for a separate experiment, we observed a mean peak N1 amplitude of around -5 $\mu$V. Assuming a proportional difference of .15, we therefore expected a .75 $\mu$V reduction in N1 amplitudes at the highest level of predictability, relative to the lowest level of predictability, in the picture-congruent condition. The values we predicted for the extremities of each independent variable are presented in Table 3.

In each iteration of the simulation, we simulated 200 (100 per congruency condition) trials for each of $N$ subjects with subject-, picture-, and word-specific random intercepts and slopes. The predictability values were taken directly from the generated stimuli. The simulation can be understood through reference to the formula that describes the linear mixed effects model:

**Table 3**

*The coding method and predicted N1 amplitudes for the extremities of each predictor variable. As congruency is deviation-coded and there are an equal number of congruent and incongruent trials, the values for $Cong_{spw}$ are presented as between -.5 and .5, though the actual values are likely to differ slightly after observations fitting exclusion criteria are removed (in both the simulation and actual analysis). $Pred_{spw}$ values are calculated as proportion of agreement normalized between 0 and 1.*

| Congruency | $Cong_{spw}$ | Percentage of modal name agreement (%) | $Pred_{spw}$ | Predicted N1 amplitude ($\mu$V) |
|---|---|---|---|---|
| Incongruent | -.5 | 7 | 0 | -5.00 |
| Incongruent | -.5 | 100 | 1 | -5.00 |
| Congruent | .5 | 7 | 0 | -5.00 |
| Congruent | .5 | 100 | 1 | -4.25 |

$$y_{spw} = \beta_0 + S_{0s} + P_{0p} + W_{0w} + (\beta_1 + S_{1s} + P_{1p})Cong_{spw} + (\beta_2 + S_{2s})Pred_{spw}$$

$$+(\beta_{12} + S_{12s})Cong_{spw}Pred_{spw} + e_{spw}$$

Table 4 explains each term in this model and presents the values simulated for the power analysis. The simulated values for the fixed effects were calculated based on the predictions and coding scheme, and are also presented in Table 4. The simulated values of subject random intercepts were based on mixed effects models for N1 amplitudes in prior research from the lab (*citation removed for double-blinding*), where subject random effects showed much greater variability between subjects than items. The variance for the distribution residuals was also based on estimates from mixed effects models in these analyses. Due to the coding method of the coefficients, the $\beta$ terms in the table and equation above can be interpreted as follows:

$\beta_0$ reflects the average amplitude at the lowest level of predictability,

$\beta_1$ reflects the difference between congruent and incongruent trials at the lowest level of

**Table 4**

*The meaning of each term in the design's linear mixed effects model, and the value simulated for the power analysis. Where simulated variables were drawn from distributions, $\sim N(\mu, \sigma)$ indicates that the respective variable's values were drawn from a normal distribution with mean $\mu$ and standard deviation $\sigma$.*

| Term | Meaning | Simulated Value ($\mu$V) |
|---|---|---|
| $y_{spw}$ | Trial-level N1 amplitudes for subject $s$, picture $p$, and word $w$ | |
| $\beta_0$ | Grand intercept | $= -5$ |
| $S_{0s}$ | Subject random intercept for subject $s$ | $\sim N(0, 2.5)$ |
| $P_{0p}$ | Picture (image) random intercept for picture $p$ | $\sim N(0, 2.5)$ |
| $W_{0w}$ | Word random intercept for word $w$ | $\sim N(0, 2.5)$ |
| $\beta_1$ | Fixed effect of congruency | $= 0$ |
| $S_{1s}$ | Subject random slope for congruency for subject $s$ | $\sim N(0, .75)$ |
| $P_{1p}$ | Picture (image) random slope for congruency for picture $p$ | $\sim N(0, .5)$ |
| $Cong_{spw}$ | Trial-level congruency values (deviation-coded) | |
| $\beta_2$ | Fixed effect of predictability | $= .375$ |
| $S_{2s}$ | Subject random slope for predictability for subject $s$ | $\sim N(0, 1)$ |
| $Pred_{spw}$ | Trial-level predictability values | |
| $\beta_{12}$ | Fixed effect of congruency-predictability interaction | $= .75$ |
| $S_{12s}$ | Subject random slope for congruency-predictability interaction for subject $s$ | $\sim N(0, 1)$ |
| $e_{spw}$ | Residual random noise | $\sim N(0, 3)$ |

predictability,

$\beta_2$ reflects the overall effect of predictability across congruent and incongruent trials, and

$\beta_{12}$ reflects the difference between congruent and incongruent trials at the highest level of predictability.

In each simulation, simulated participants were pseudo-randomly assigned to stimulus sets 1 and 2 in equal number, or with randomly allocated counts of $\frac{N}{2} - 0.5$ and $\frac{N}{2} + 0.5$ if the number of simulated participants were odd. $N$ varied from 10 to 100 in steps of 5, with 500 iterations run at each value. Before models were fit to simulated data in each iteration, data exclusion was simulated as a random 10% loss of trials. The first 5% simulated data loss observed in the stimuli validation due to trials being responded to incorrectly or with response times less than 250 ms or greater than 1500 ms. No lower bound for response time exclusions was applied in the EEG experiment, as the word was visible for 1 second before responses are permitted. As a conservative estimate, however, we expected a similar percentage of data loss to that seen in the validation of the picture word stimuli. The remaining 5% of data loss was simulated because, given the participant exclusion criteria, this is the maximum allowable loss of data due to a combination of technical problems with the EEG system. This conservative estimate can be considered a worst-case scenario in terms of EEG data loss. The possibility of participants being excluded was not simulated, as we opted to simply continue collecting data until we reached the desired number of participants, and excluded participants' data would not be analysed. Covarying random effects were simulated using the R package *faux* (DeBruine, 2020). Linear mixed effects models were fit using the same functions, formula, and optimiser as those used for the analysis of the actual data. In the case of non-convergence, models were re-fit without random correlations before significance testing, as this is the action we would take when modelling the actual data. Likelihood ratio Chi-square model comparisons were conducted between the full model and a version of the model lacking the interaction term, and the resulting $p$ values were recorded from each iteration.

Given that the hypothesis was directional, simulated significance tests were performed using one-tailed comparisons with an alpha level of .05. Running only 500 simulations is likely to give noisy estimates of power when simulating data which can vary in many parameters. Since fitting a much larger number of models would be unfeasible due to the time taken to fit each mixed effects model, the underlying relationship between the number of participants and the design's statistical power was estimated by fitting log-linear binomial generalised linear models (GLMs) to all iterations for one-tailed and two-tailed comparisons. Figure 5 depicts the resulting power curves. The power analysis suggested that a sample size of 68 participants (divisible by four, so as to assign an equal number of participants to each combination of counterbalanced response and stimulus groups) would be sufficient to reach at least 80% power for detecting the effect of interest in the predicted direction with a one-tailed comparison. Specifically, the model predicted that at this number of participants, assuming the predicted effect exists, we could expect 81.72% power (99% confidence interval = [80.46%, 82.91%]).

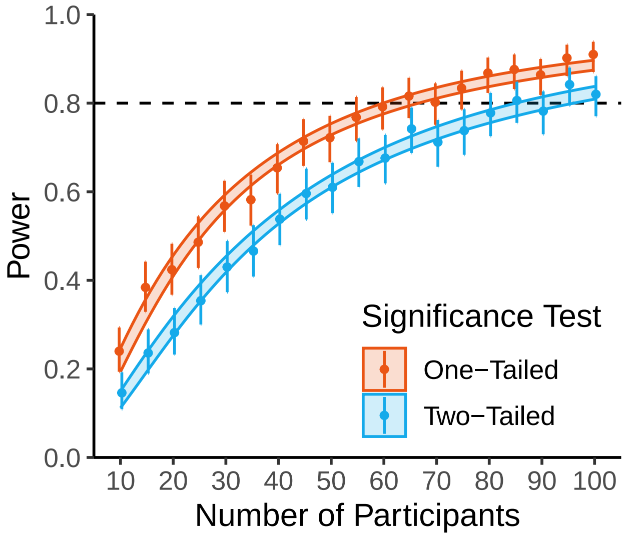

**Figure 5**

*Power curves calculated from the simulations. For comparison, both one-tailed and two-tailed power are presented, though the* p *value used in the actual planned analysis is one-tailed. Points (shifted horizontally for visibility) present the observed proportions of simulations which resulted in statistically significant p values. Vertical error bars present 99% binomial confidence intervals of these individual proportions. The coloured lines showing a logarithmic relationship depict the upper and lower bounds of 99% confidence intervals of predicted probabilities from log-linear binomial GLMs fit to the data. The dashed horizontal line highlights the 80% power target.*

We note that due to a lack of relevant data from similar designs, variance-covariance matrices for the power analysis were simulated with all random effects correlations set to zero. To check this did not result in heavily biased estimates, the power analysis was also run with all random effects correlations set to values of .2, .4, .6, and .8. Each of these analyses estimated a strikingly similar relationship between the number of participants and statistical power (Figure 6).

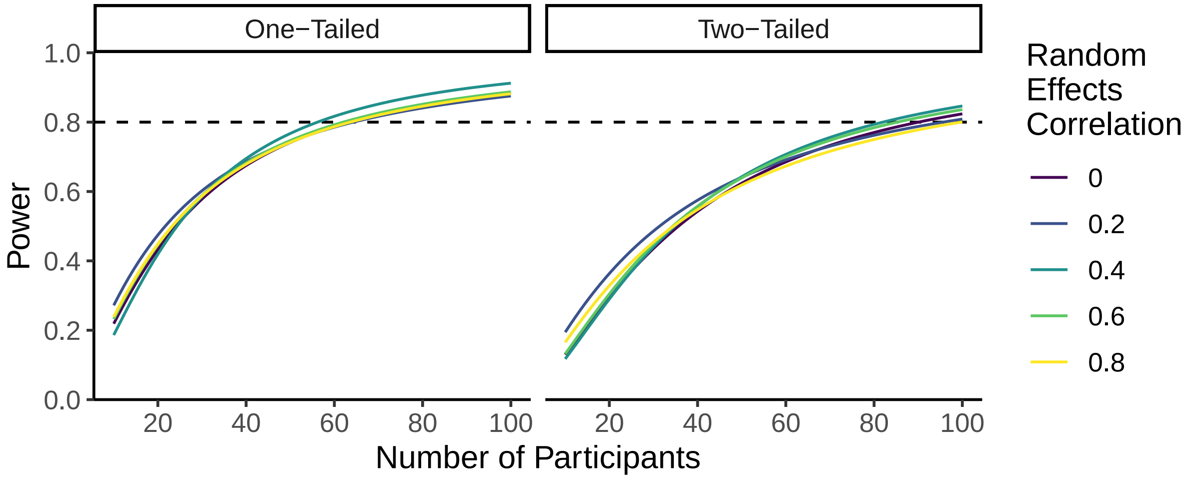

**Figure 6**

*Power curves when all random effect correlations are set to 0, .2, .4, .6, and .8. Each line depicts the predicted relationship between number of participants and power from a single loglinear binomial GLM. As in the original power analysis, results were simulated with N of 10 to 100 in steps of 5, though here with only 100 simulations at each step rather than 500. The overall relationship between the number of participants and the statistical power for finding the predicted interaction remains mostly unchanged across different random effects correlations. As in Figure 5, both one-tailed and two-tailed power are presented, though the* p *value used in the experiment is one-tailed. The dashed horizontal line highlights the 80% power target.*

#### E Instructions given to Participants

Instructions for the localiser and picture-word tasks, shown below, were presented multiple times: at the start of each task, after practice trials, and before the start of each block. The words AFFIRMATIVE and NEGATIVE below were replaced with the text "Left Control" or "Right Control" respectively, depending on which response group the participant was assigned to. In the practice trials, an additional line of text read, "For the practice trials, you will be given feedback on your accuracy for each trial.". For all other trials, this line instead read, "Unlike the practice trials, you will not be given feedback on your accuracy for each trial.".

The instructions for the localiser task were as follows:

*In each trial, the following things will happen:*
*1) You will be shown a picture of a word, nonword, or noise image.*
*2) The image will turn green.*
*3) When the image turns green:*
*Press the AFFIRMATIVE key if the image is of a real word.*
*OR*
*Press the NEGATIVE key if it is not of a real word.*

*Once the image changes colour, try to respond as quickly and accurately as possible.*

*When you have read these instructions, press the space key to begin...*

The instructions for the picture-word task were as follows:

*In each trial, the following things will happen:*
*1) You will be shown a picture of an object for 2 seconds.*
*2) There will be a short delay.*
*3) You will be shown a word.*
*4) The word will turn green.*

*5) When the word turns green:*

*Press the AFFIRMATIVE key if the word describes the object you saw.*

*OR*

*Press the NEGATIVE key if it does not.*

*Once the word changes colour, try to respond as quickly and accurately as possible.*

*When you have read these instructions, press the space key to begin...*

### F Change to the High-Pass Filter Cut-Off

We originally pre-registered a high-pass filter cut-off of .5 Hz. After pre-registration, we changed this to .1 Hz to address possible artefactual distortions in timings of effects (Rousselet, 2012; Tanner et al., 2015; VanRullen, 2011). Here, we report what our results would have been had we not made this alteration. All other elements of the analysis pipeline match those reported in the manuscript. This analysis reproduced the main finding of an interaction term in the opposite direction to that we expected under a simple predictive coding hypothesis. This suggests that our change to the high-pass filter cut-off did not alter our main results or conclusions.

#### Maximal Electrode Analysis

The fixed-effect relationship estimated after preprocessing with a high-pass cut-off of .5 Hz is presented in Figure 7, showing a pattern of results similar to that observed with a cut-off of .1 Hz. The model intercept was estimated to be $\beta$=-2.83 (*SE*=.46). The fixed effect of Congruency was estimated as $\beta$=-.4 (*SE*=.3), and the main effect of Predictability was estimated as $\beta$=.3 (*SE*=.26). Importantly, the effect of interest, the interaction between Congruency and Predictability, was in the opposite direction from that hypothesised, estimated as $\beta$=-1.24 (*SE*=.46), with a larger effect than that reported in the analysis using a .1 Hz filter (i.e., $\beta$=-.79 µV, *SE*=.52).

We also re-fit the Bayesian linear mixed-effects model, as described in the manuscript, to the maximal electrode data extracted after filtering with a .5 Hz cut-off. This revealed a similar posterior distribution to that reported in the manuscript, but with even less of the posterior distribution consistent with the simple predictive coding hypothesis (Figure 8). The median posterior estimate for the Congruency-Predictability interaction was $\beta$=-1.23 µV (89% highest density interval = [-1.94, –.48]). We calculated, given this posterior distribution, that the Congruency-Predictability interaction is 239.38 times more likely to be less than 0, than it is to be greater than zero (that is, $BF_{01}$).

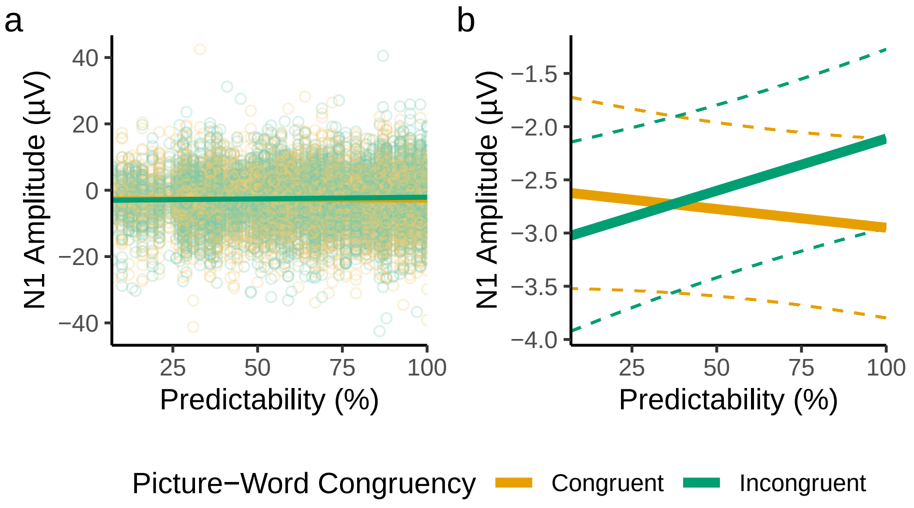

**Figure 7**

*Maximal electrode results with a .5 Hz cut-off. (**a**) Model-derived fixed-effect predictions, visualised over results from all trials (individual points). (**b**) Fixed-effect predictions visualised alone for visibility. Dashed lines depict the bounds of 95% bootstrapped prediction intervals (5,000 bootstrap samples). For feasibility, bootstrapped predictions were generated from a version of the model that lacked random slopes.*

#### Full ERP Analysis

We also re-fit the models estimating the full time-course of effects in the region of interest. Importantly, as shown in Figure 9, the Congruency-Predictability interaction term remained negative throughout the N1 period.

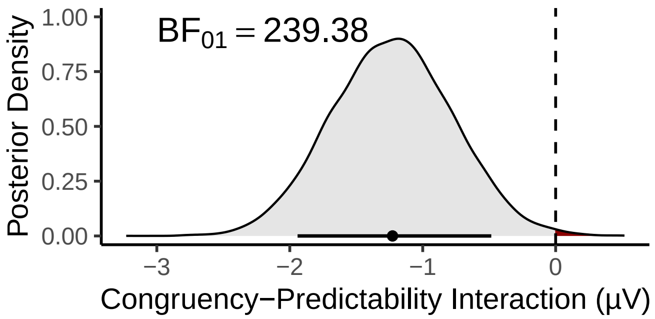

**Figure 8**

*Posterior density for the Congruency-Predictability interaction after filtering with a .5 Hz high-pass cut-off. The point below the density plot depicts the median estimate; the horizontal line shows the 89% HDI of the posterior distribution.*

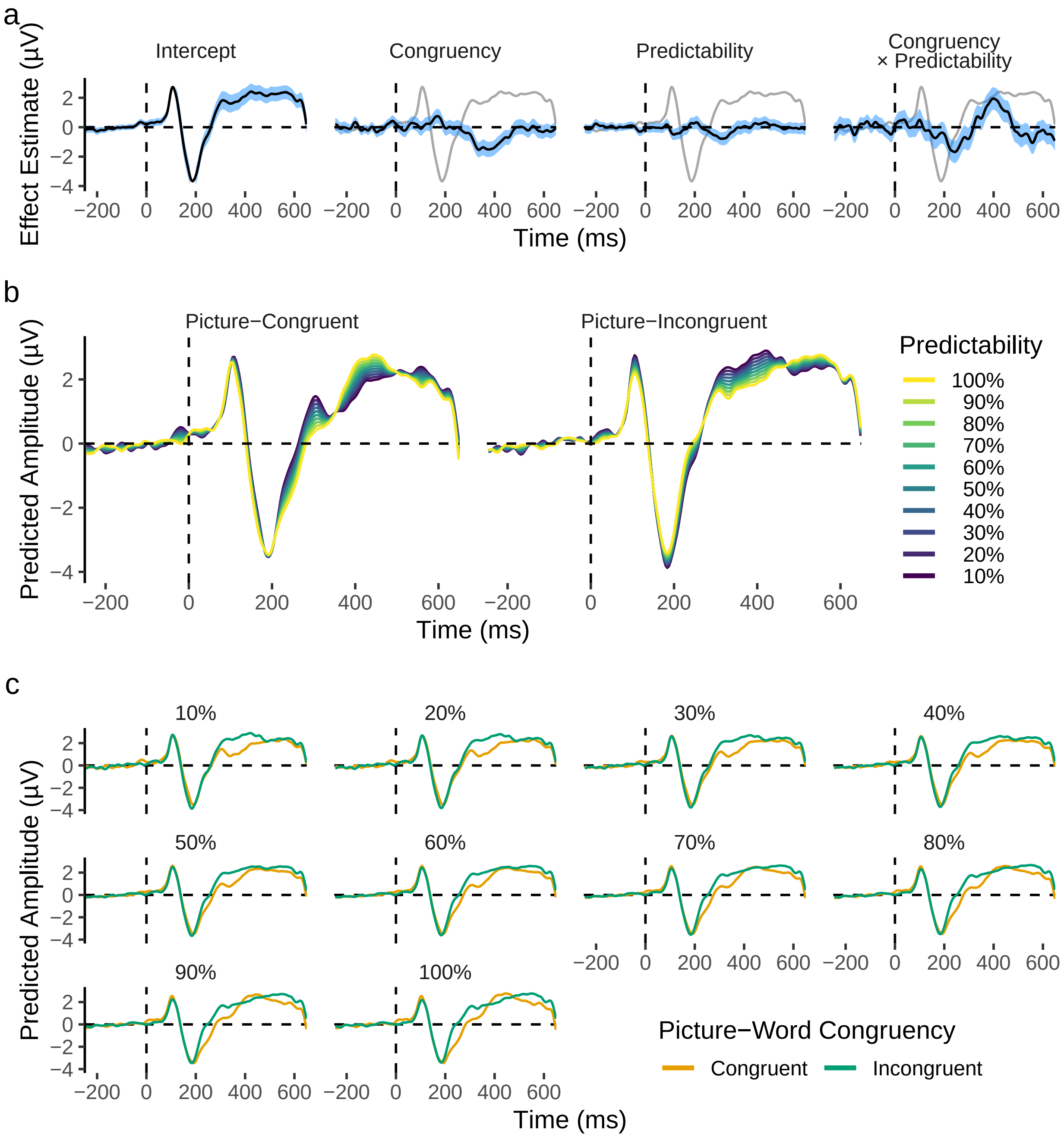

**Figure 9**

*Time-course of fixed effects from the sample-level analysis of the left-lateralised occipitotemporal region of interest, when filtering with a .5 Hz high-pass cut-off. (**a**) Time-course of fixed-effect estimates. (**b**) Fixed effect predictions showing how predictability affects amplitudes. (**c**) Fixed effect predictions contrasting congruent and incongruent ERPs at each level of predictability.*

#### G Behavioural Results from the Picture-Word Task

We conducted exploratory behavioural analyses of data from the picture-word EEG task.

##### Response Times Results

We analysed response times (RTs) to examine whether the pattern of effects was similar to that observed for the behavioural validation experiment (Supplementary Materials B). We fit a Bayesian distributional shifted log-normal model, estimating the same model formula as that described for the behavioural validation experiment for all shifted log-normal parameters ($\mu$, $\sigma$, and $\delta$):

```
rt ~ 1 + congruency * predictability +
        (1 + congruency * predictability | subject_id) +
        (1 + congruency | image_id) +
        (1 | word_id),
sigma ~ 1 + congruency * predictability +
        (1 + congruency * predictability | subject_id) +
        (1 + congruency | image_id) +
        (1 | word_id),
ndt ~ 1 + congruency * predictability +
        (1 + congruency * predictability | subject_id) +
        (1 + congruency | image_id) +
        (1 | word_id)
```

The parameter of $\mu$ was modelled with an identity link function, while $\sigma$ and $\delta$ were modelled with log link functions. We specified prior distributions based on the posterior distributions from the behavioural validation experiment. Priors for the behavioural analysis of the EEG experiment were not exact replicas of the validation experiment's posteriors, but were rather specified with greater uncertainty than that observed in the validation experiment's posteriors. We decided to specify this uncertainty because of key differences in the task demands: participants in the validation experiment could respond to stimuli without lower limit, whereas in the EEG experiment, responses were only permitted 500 ms after stimulus presentation. As a result of the additional time participants had to consider their responses, and because RTs were measured from the time point at which the stimulus changed colour, we reasoned that (1) responses would be faster overall in the EEG experiment (reflected in a reduced prior for the $\delta$ parameter intercept), and (2) effects observed in the validation experiment would be likely smaller in the EEG experiment. Specifically, fixed and random effect prior distributions for the $\mu$ and $\sigma$ parameters, and random effect priors for $\delta$, were specified such that they were centred on the median estimate from the stimulus validation analysis, but with variance of the random effects ten times that observed in the stimulus validation posterior distributions. The fixed effect prior distributions for the $\delta$ parameter were specified to be more uninformative than this, as we expected this parameter to change the most. The prior distribution for the $\delta$ intercept was drawn from $\sim N(0, 7.5)$, while the fixed effect slopes' priors also had *SD*s of 7.5, but were centred on the posterior estimates from the stimulus validation analysis. Priors for all correlations of effects were kept as the *brms* default of a flat distribution between -1 and 1. The model was fit with 5 Markov chains, each with 10,000 (7,500 warm-up and 2,500 sampling) iterations. The *adapt_delta* parameter was set to .99, and the *max_treedepth* parameter was set to 10. Summaries of the fixed effect posterior distributions, relative to those of the priors, are shown in Figure 10. Similar results are shown for all random effects in Figure 11.

Results revealed that, although the effects were smaller than in the validation experiment, the main finding was replicated, with low predictability eliciting later RTs for picture-congruent words, to a greater extent than it does for picture-incongruent words (Figure 12a). RTs from the EEG experiment also replicated the difference in spread between picture-congruent and -incongruent RTs at low levels of predictability, with the congruency conditions showing more similar spread in RTs as predictability increases (Figure 12b). Again, this effect was smaller for RTs in the EEG experiment than it was for RTs in the validation experiment. Conversely, the difference in shift observed between picture-congruent and -incongruent words at high predictability in the validation experiment was not observed in the EEG experiment.

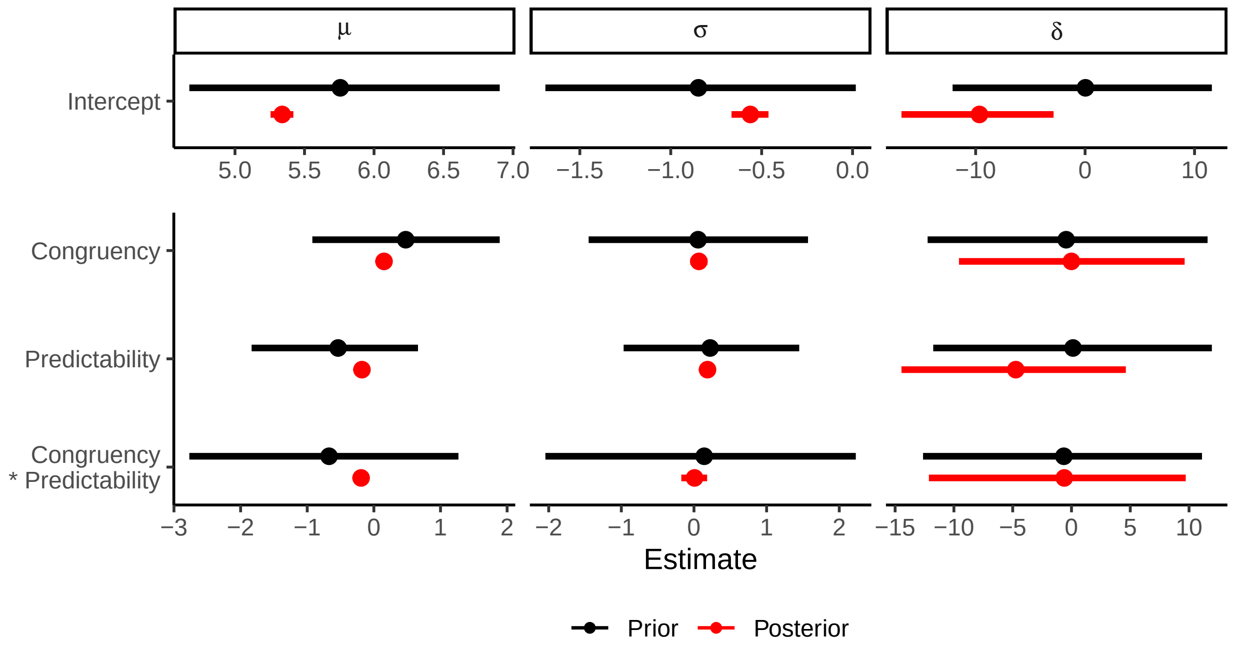

**Figure 10**

*Prior and posterior distributions for all fixed effects estimated in the Bayesian shifted log-normal model of the RT data from the EEG experiment's picture-word task, for the* $\mu$*,* $\sigma$*, and* $\delta$ *parameters. Points depict median estimates, while whiskers depict 89% HDIs, for prior* (black) *and posterior* (red) *distributions.*

#### Accuracies Results

We similarly analysed accuracies in the picture-word task. We fit a logit-link binomial Bayesian generalised linear mixed effects model (GLMM) to accuracy data, using the same maximal mixed effects formula as that described for the planned analysis of EEG data. All fixed effect prior distributions were specified to be flat, with the exception of the model intercept. As we expected overall accuracy to be very high, we specified the prior distribution for the fixed effect intercept as $\sim N(4,1)$, where logit 4 would be equivalent to an average accuracy of .982. Priors for the *SD*s of random effects distributions were drawn from Student's $t$ distributions with 3 degrees of freedom, $\mu$ of 0, and $\sigma$ of 2.5. Prior distributions for all correlations were flat (between -1 and 1). The model was fit via *brms*, with 5 chains each sampling for 10,000 iterations (5,000 warmup). The *adapt_delta* parameter was set to .9, and the maximum tree depth (*max_tree_depth*) was set to 10.

Results revealed a main effect of predictability with higher accuracy at higher levels of predictability (Figure 13). An interaction with congruency was also observed, where predictability had a larger effect for picture-congruent than for picture-incongruent words, while accuracy remained more consistent across predictability for picture-incongruent words.

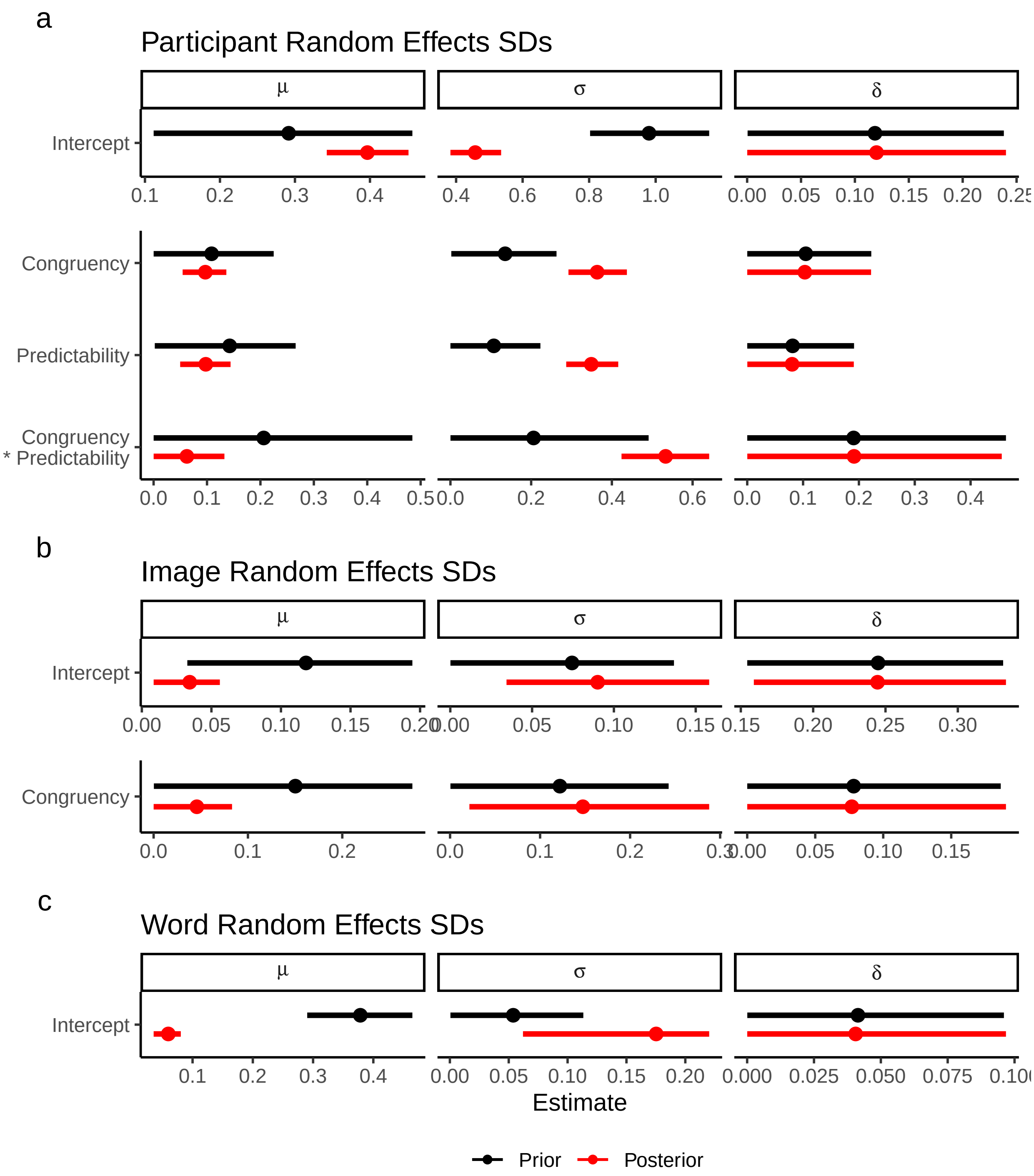

**Figure 11**

*Prior and posterior distributions for all random effects estimated in the Bayesian shifted log-normal model fit to describe RT data from the EEG experiment's picture-word task, for the $\mu$, $\sigma$, and $\delta$ parameters. Results are separately for (**a**) participant, (**b**) image, and (**c**) word random effects. Points depict median estimates, while whiskers depict 89% HDIs, for prior (*black*) and posterior (*red*) distributions.*

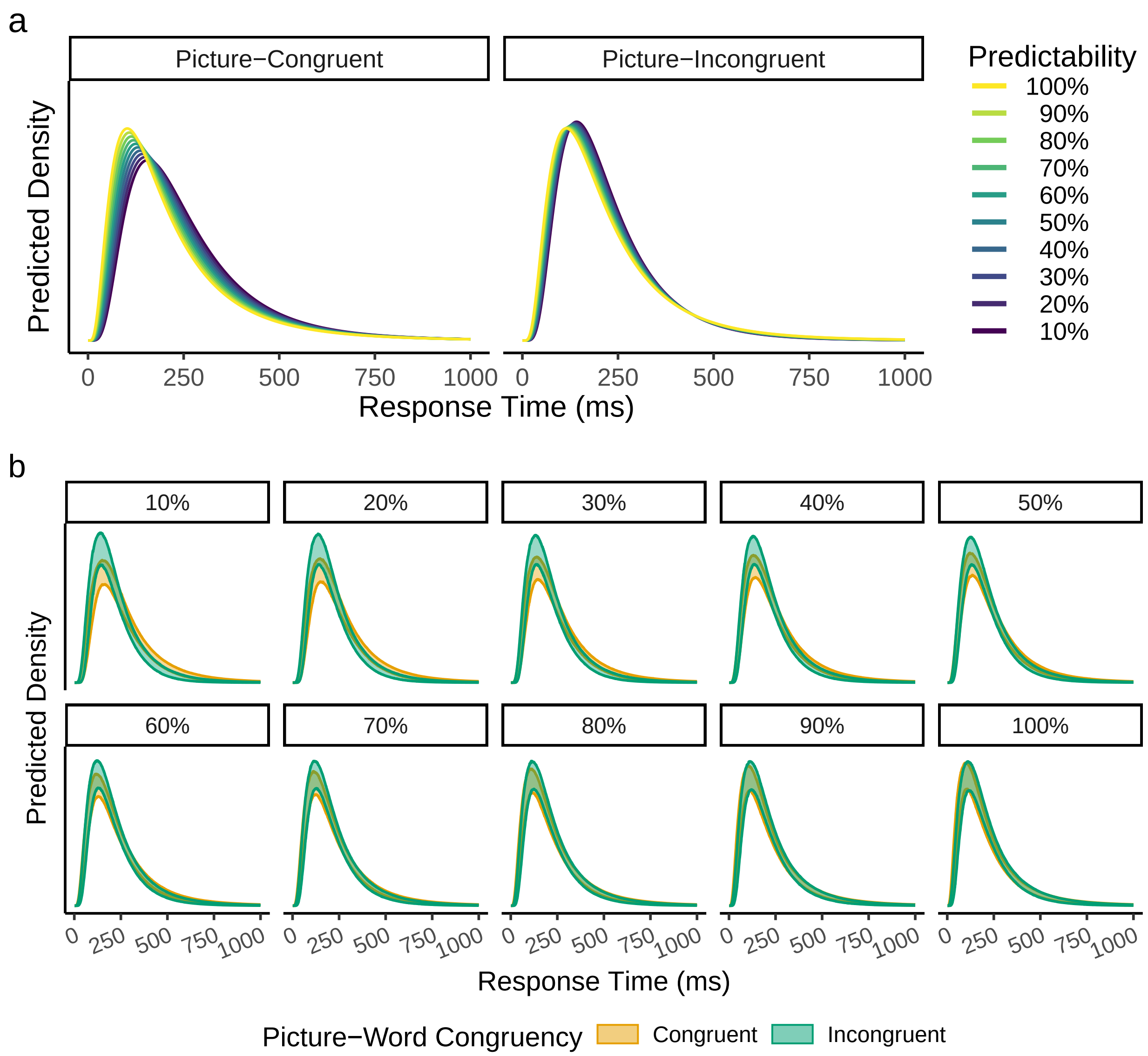

**Figure 12**

*Fixed effect predictions of RT distributions in the EEG experiment. Figure layout is identical to that described for the validation experiment RTs, except that the axis limits for RTs are here limited to $\leq$1,000 ms. Unlike the validation experiment, where RTs reflect latency from stimulus presentation, RTs here reflect latency from a colour change in the stimulus, that occurred 500 ms after stimulus presentation.*

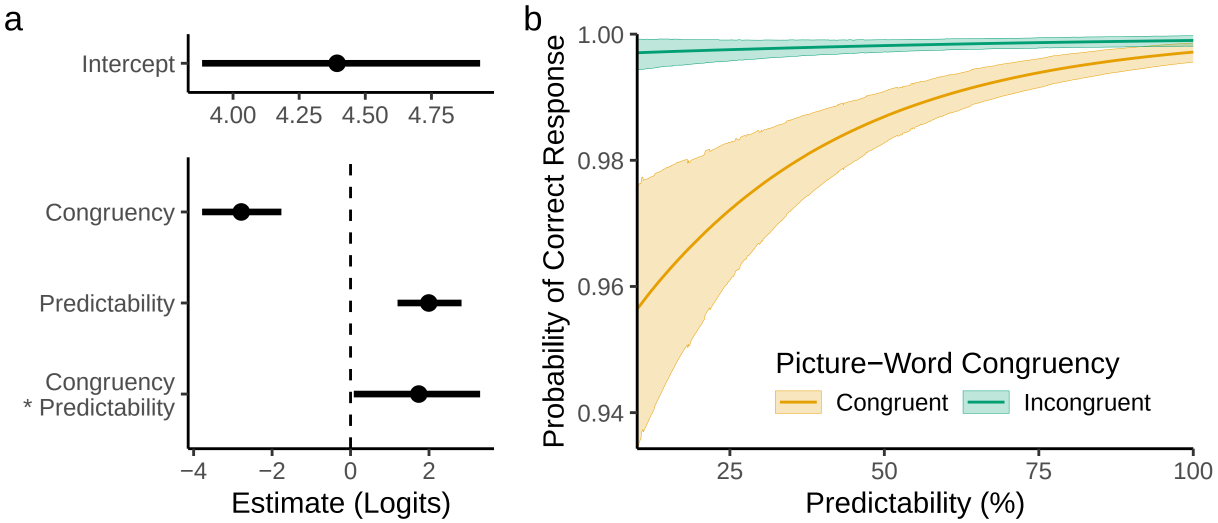

**Figure 13**

*Fixed effect results for the analysis of accuracies in the picture-word task during the EEG experiment. (**a**) Fixed effect logit estimates, where points depict median estimates and whiskers depict 89% HDIs. (**b**) Model-predicted accuracies, for all levels of predictability in each congruency condition, where the central lines depict median estimates, while the shaded areas depict 89% HDIs.*

#### H Results from the Localiser Task

We analysed results from localiser task, examining the full time-course of stimulus effects on ERP amplitudes, and patterns of RTs and accuracies.

##### ERP Results

We analysed the full time-course of stimulus effects in the localiser task, for right- and left-hemispheric occipitotemporal regions of interest. Specifically, separate linear mixed effects models to each time point (256 Hz) via *lme4*, estimating models with the following formula:

```
amplitude ~ 1 + (false_font + noise) * hemisphere +
  (1 | participant_id) +
  (1 | participant_id:electrode_id) +
  (1 | match_set) +
  (1 | item_id)
```

Here, *false_font* and *noise* were deviation-coded variables, comparing the two nonword conditions to the null condition of words (i.e., BACS-font nonwords, and phase-shuffled words, respectively). In this way, the fixed effect slopes represented the difference between words and each non-lexical stimulus type. The deviation-coded variable, *hemisphere*, distinguished observations in the left (*hemisphere*=-.5) and right (*hemisphere*=.5) hemisphere. The *match_set* variable uniquely identified each triplet of matched items. As in the sample-level analysis of the picture-word task, random intercepts were also estimated for each combination of participant and electrode (*participant_id:electrode_id*), and random slopes were excluded for feasibility.

Results revealed that differences between words and phase-shuffled words emerged clearly in the P1 component, with more positive-going amplitudes observed for phase-shuffled words (Figure 14). Differences between words and false-font nonwords, meanwhile, remained small until later, in the N1. In both hemispheres, N1 components were more negative-going for false-font stimuli than for phase-shuffled words. Both positive-going and negative-going

ERP components elicited by words were overall more positive in amplitude for the right hemispheric occipitotemporal electrodes, that is, the P1 was more positive-going, and the N1 less negative-going, in the right hemisphere. The N1 elicited by word stimuli was left-lateralised. An interesting stimulus-hemisphere interaction was observed, wherein ERPs elicited by words showed N1 peak amplitudes most similar to false-font stimuli in the left hemisphere, but most similar to phase-shuffled words in the right-hemisphere. Similar differences in timing were observed in the N1 peak for stimuli across both hemispheres, with phase-shuffled words peaking first, followed by false-font stimuli, and then words. Stimulus effects at occipitotemporal electrodes after the N1 were more consistent across hemispheres, with phase-shuffled words showing the most positive amplitudes, followed by false-font stimuli, which in turn elicited more positive amplitudes than words did, although the difference between words and phase-shuffled words was larger, post-N1, in the right hemisphere. The post-N1 difference between words and false-font nonwords, meanwhile, did not interact with hemisphere except for a brief period around 250 ms.

**Behavioural Results**

We also analysed stimulus effects on lexical decision RTs and accuracies. Specifically, we fit a logit-link binomial model to trial-level accuracies, and a distributional shifted log-normal model to RTs, with maximal random effects structures.

Accuracies were modelled via a logit-link binomial model (Figure 15a) with an informative prior for the model's logit intercept of $\sim N(5,1)$ (centred on average accuracy of .993), reflecting the expectation that accuracy overall would be very high. Weakly informative priors were defined for fixed effect slopes ($\sim N(0,5)$) and for the *SD*s of random effect distributions ($\sim t(5,0,1)$). Prior distributions for correlations within the model were specified to be flat. The model was fit with 5 chains, each with 10,000 iterations (7,500 warmup, 2,500 sampling). The *adapt_delta* parameter was set to .99, and the *max_tree_depth* was set to 10. In *brms* syntax, the model estimated coefficients from the following formula:

```
correct ~ 1 + false_font + noise +
  (1 + false_font + noise | participant_id) +
  (1 + false_font + noise | match_set) +
  (1 | item_id)
```

RT data were modelled with a shifted log-normal model (Figure 15b). The parameter of $\mu$ was modelled with an identity link function, while $\sigma$ and $\delta$ were modelled with log link functions. The maximal random effects structure was estimated for the distributional parameters $\mu$ and $\sigma$, whereas the $\delta$ parameter was modelled with a global intercept only. This decision was based on persistent divergent transitions in the Hamiltonian Monte Carlo sampler used to explore the model's parameter space. These divergences were caused by the extremely low shift (non-decision time) in the RT data from the EEG experiment, which approached 0 ($-\infty$ on a log scale). This problem is also the likely cause of the high uncertainty for the $\delta$ intercept, and also for the parameter coefficients in the analysis of the RT data from the picture-word task. Priors for fixed-effect intercepts were specified to be centred on posterior averages from the picture-word study RT analysis, though with additional uncertainty specified in the distributions to reflect the expectation that RT distributions would differ somewhat from the picture-word task. Specifically, the intercept for $\mu$ was specified as $\sim N(5.3, 1)$, $\sigma$ as $\sim N(-.56, 1)$, and $\delta$ as $\sim N(-9, 5)$. Priors for fixed effect slopes were specified as $\sim N(0, 1)$. Prior distributions for the *SD*s of random effects were drawn from Student's $t$ distributions centred on 0, with 5 degrees of freedom and a $\sigma$ parameter of 1. Prior distributions for all correlations were flat. As with the model of accuracies, the RT model was fit with 5 chains, each with 10,000 iterations (7,500 warmup, 2,500 sampling). The *adapt_delta* parameter was set to .9, and the *max_tree_depth* was set to 10. In *brms* syntax, the model estimated coefficients from the following formula:

```
rt ~ 1 + false_font + noise +
  (1 + false_font + noise | participant_id) +
  (1 + false_font + noise | match_set) +
  (1 | item_id),
sigma ~ 1 + false_font + noise +
```

```
  (1 + false_font + noise | participant_id) +
  (1 + false_font + noise | match_set) +
  (1 | item_id),
ndt ~ 1
```

Results revealed that responses were fastest and most accurate for phase-shuffled words (Figure 16). Responses were slowest and least accurate for word stimuli. RT distributions were similar for false-font and phase-shuffled words, though accuracies for false-font stimuli were closer to those observed for words. Behavioural results overall suggest that participants found it easy to reject phase-shuffled words in lexical decision, but found it relatively more difficult to reject false-font stimuli.

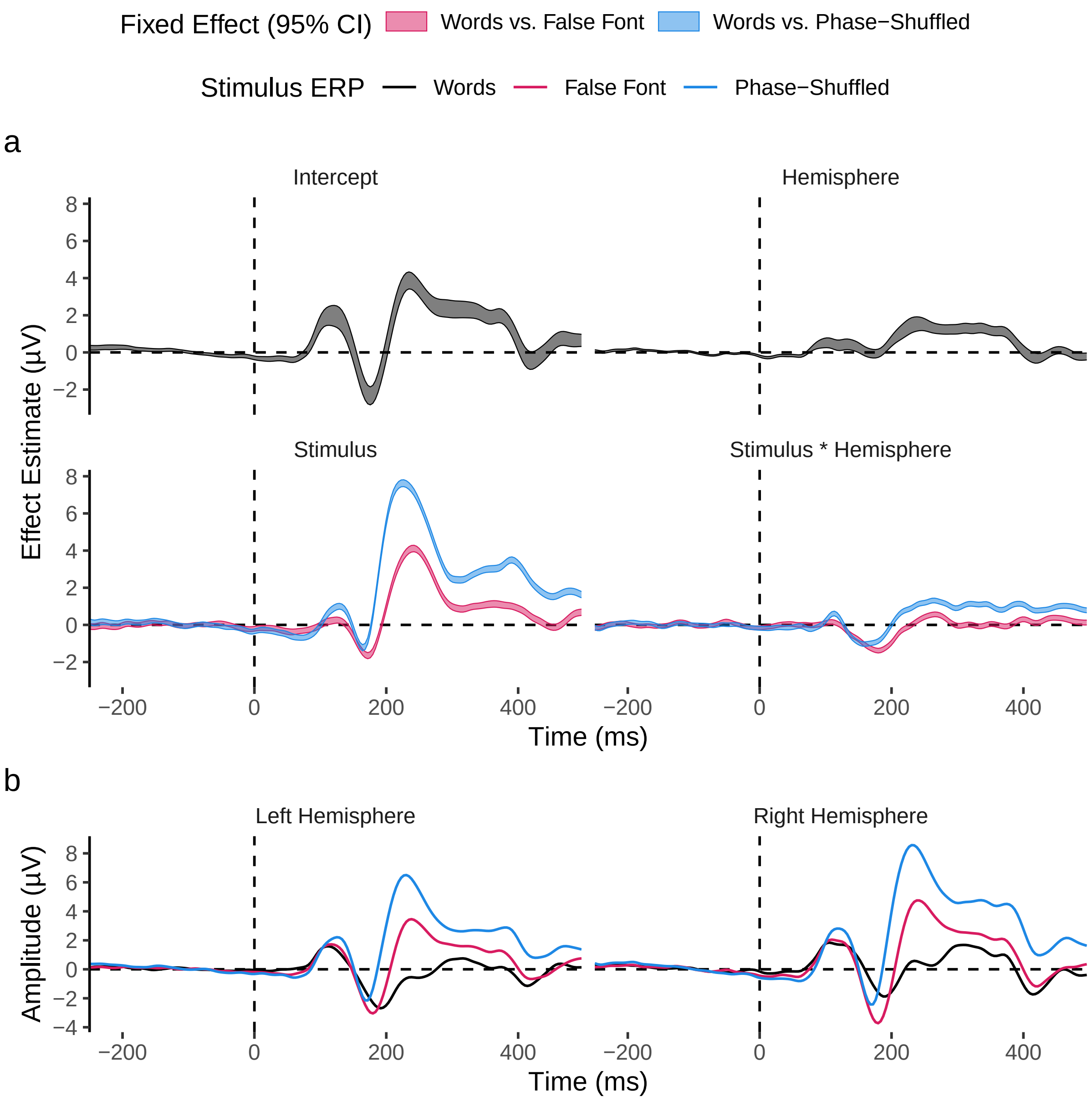

**Figure 14**

*Fixed effect results for ERPs in the localiser task. (a) Fixed effects estimates for each time point, with the shaded areas depicting 95% confidence intervals. (b) Model-derived predictions for ERPs of left- and right-hemispheric occipitotemporal electrodes.*

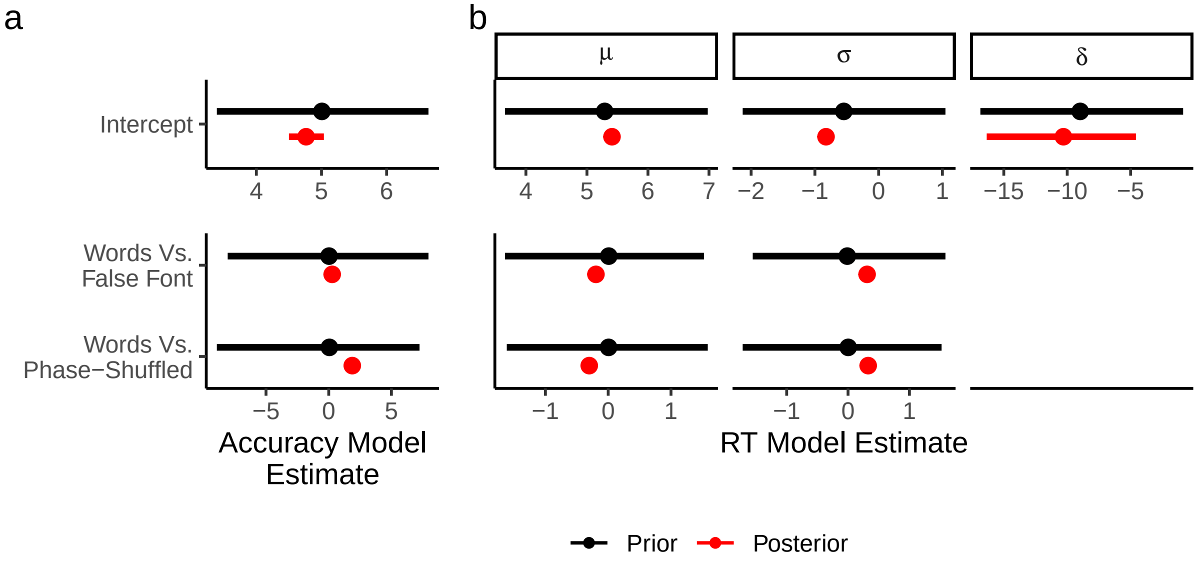

**Figure 15**

*Prior and posterior distributions for all fixed effects estimated by the (**a**) logit-link Binomial model to describe accuracies and (**b**) Bayesian shifted log-normal model fit to describe RT data from the localiser task. Estimates in (**a**) are in logit units. Estimates in (**b**) are depicted for each shifted log-normal parameter separately. In both panels, points depict median estimates, while whiskers depict 89% HDIs, for prior (*black*) and posterior (*red*) distributions.*

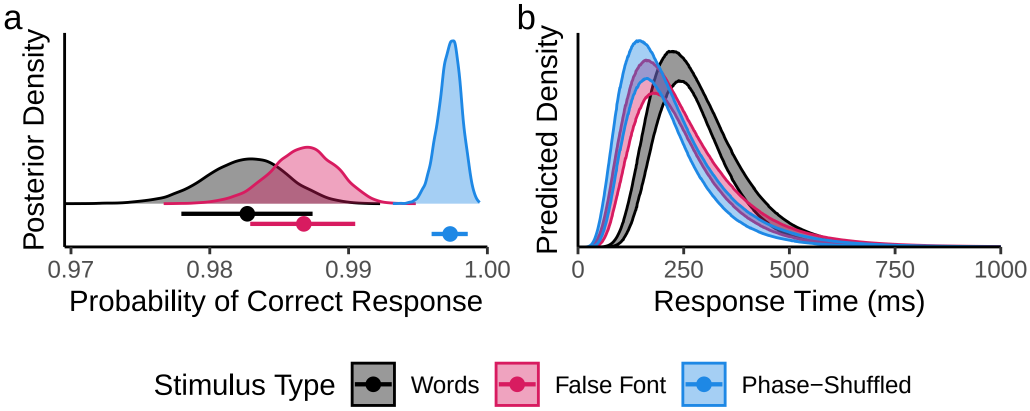

**Figure 16**

*Fixed effect predictions for behavioural outcomes in the localiser task. (**a**) Posterior distributions for accuracies in the localiser task, where points below densities depict median posterior estimates, while whiskers depict 89% HDIs of posterior samples. (**b**) Predicted RT distributions, where the shaded regions depict 89% HDIs of posterior samples (density values on the* y*-axis begin at 0).*

#### I Exploratory Alterations to the Maximal Electrode Method

In our planned analysis of the Picture-Word task, we modelled amplitudes recorded at electrodes which also showed, for a given participant, the maximum sensitivity to the word-false-font difference in the Localiser task. We conducted exploratory analyses to examine the impact on our results of different methodologies for extracting trial-level amplitudes from the Picture-Word EEG recordings. For both methodologies, all other parts of the analysis pipeline match those used in the planned analysis. Both methodologies revealed similar results to our planned analysis, with an interaction term in the opposite direction to that expected by our simple predictive coding hypothesis.

##### Using the Word-Noise Difference

First, rather than identifying maximal electrodes as those that showed maximal sensitivity to the word-false-font difference, we instead identified electrodes that showed maximal sensitivity to the difference between words and phase-shuffled (noise) images. As in the planned analysis, we also used the per-participant peak in this difference to identify the time point at which amplitudes should be extracted. This revealed a similar pattern of effects to that obtained in our planned analysis, with higher predictability eliciting smaller N1s for picture-incongruent words, and larger N1s for picture-congruent words (Figure 17).

To summarise the fixed effects, the model intercept was estimated to be $\beta$=-4.1 µV ($SE$=.55). The interaction was estimated to be $\beta$=-1.76 µV ($SE$=.55). The main effect of congruency was estimated to be $\beta$=.75 µV ($SE$=.37), and the main effect of predictability was $\beta$=-.02 µV ($SE$=.35).

In a Bayesian model identical to that described in the manuscript, but with amplitudes from the alternative maximal electrodes as the outcome variable, the Posterior distribution for the Predictability-Congruency was estimated to be $\beta$=-1.73 µV (89% highest density interval = [-2.6, -.88]; 1562 times more likely to be less than 0 than it is to be greater than zero **(Figure 18)**.

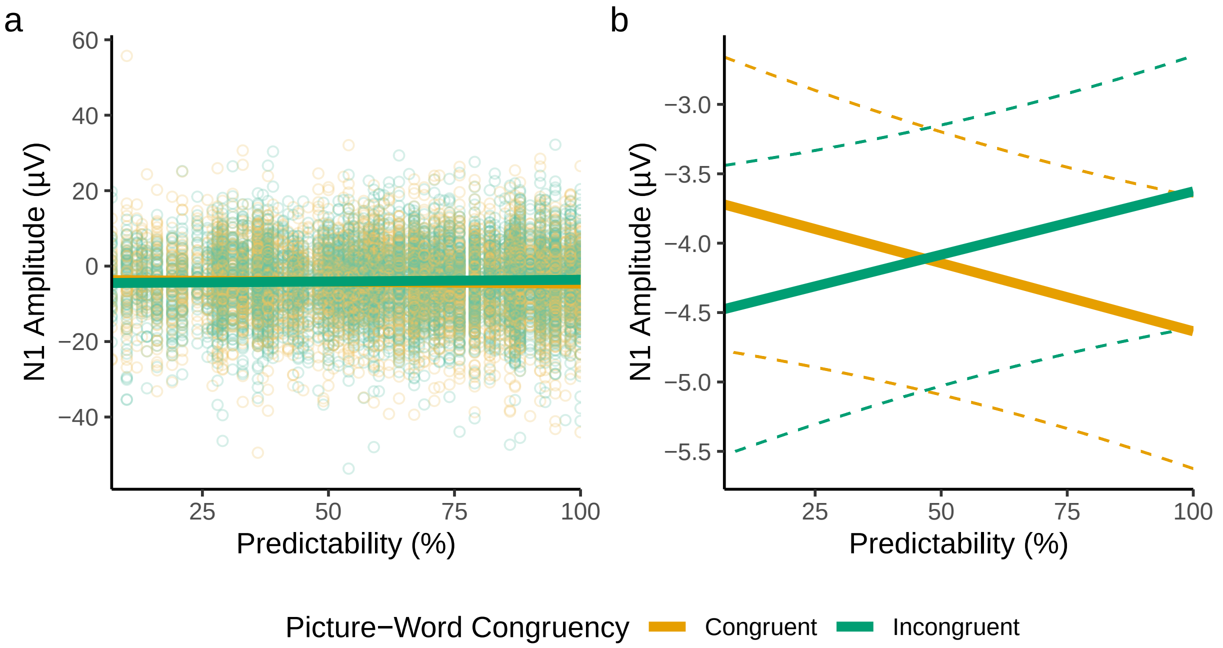

**Figure 17**

*Results using the word-noise difference to identify maximal electrodes. (**a**) Model-derived fixed-effect predictions, visualised over results from all trials (individual points). (**b**) Fixed-effect predictions visualised alone for visibility. Dashed lines depict the bounds of 95% bootstrapped prediction intervals (5,000 bootstrap samples). For feasibility, bootstrapped predictions were generated from a version of the model without random slopes.*

##### Using Region-of-Interest Averages

Second, we calculated trial-level amplitudes as the average amplitude across all electrodes in the left-hemispheric occipitotemporal region of interest, across all time points in a 120-200 ms N1 window. Again, this analysis revealed very similar results to our planned analysis **(Figure 19)**.

Here, the model intercept was estimated to be $\beta$=-2.95 µV ($SE$=.27). The interaction was estimated to be $\beta$=-1.03 µV ($SE$=.3). The main effect of congruency was estimated to be $\beta$=-.12 µV ($SE$=.21), and the main effect of predictability was $\beta$=ß.05 µV ($SE$=.19).

In a Bayesian model identical to that described in the manuscript, but with the ROI average as the outcome variable, the Posterior distribution for the Predictability-Congruency was estimated to be $\beta$=-1.03 µV (89% highest density interval

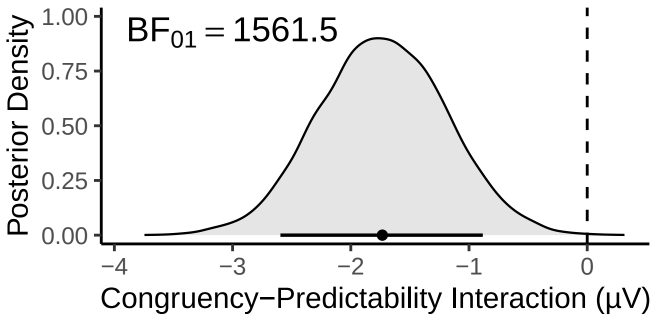

**Figure 18**

*The region of the posterior distribution consistent with the predictive coding hypothesis (where β>0) is highlighted in red. The point and horizontal line below the density plot depict respectively the median estimate and 89% highest density interval of the posterior distribution.*

= [-1.52, -.058]; 2082 times more likely to be less than 0 than it is to be greater than zero **(Figure 20)**.

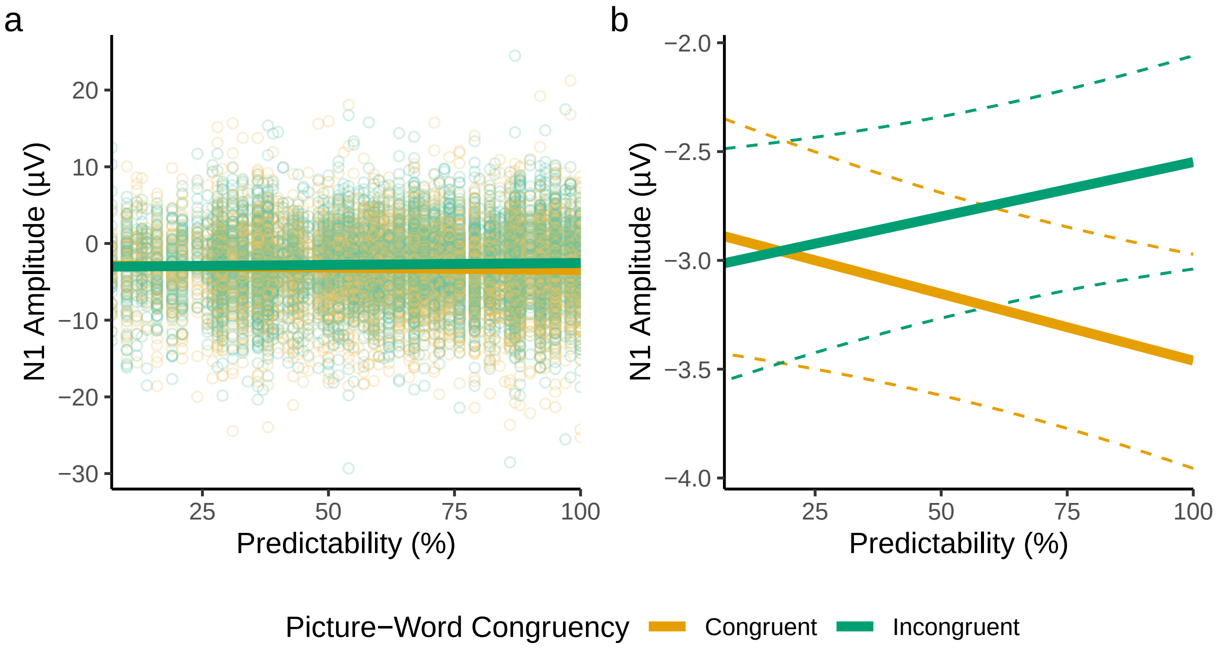

**Figure 19**

*Results from trial-level models of region-of-interest averages in the N1 window. (**a**) and (**b**) are as described in Figure 17.*

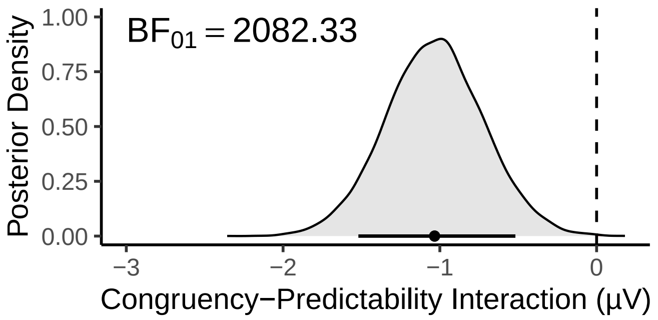

**Figure 20**

*The region of the posterior distribution consistent with the predictive coding hypothesis (where $\beta$>0) is highlighted in red. The point and horizontal line below the density plot depict respectively the median estimate and 89% highest density interval of the posterior distribution.*

#### J Model Estimates for the N400

The exploratory scalp-wide analysis of ERPs revealed a Congruency-Predictability interaction during the period of the N400 component. To more clearly interpret this effect, we generated topographic plots of estimated N400 amplitudes at different levels of predictability, for picture-congruent and -incongruent words separately (**Figure 21**).

**Figure 21**

*Topographic plots of model estimates at 400 ms.*

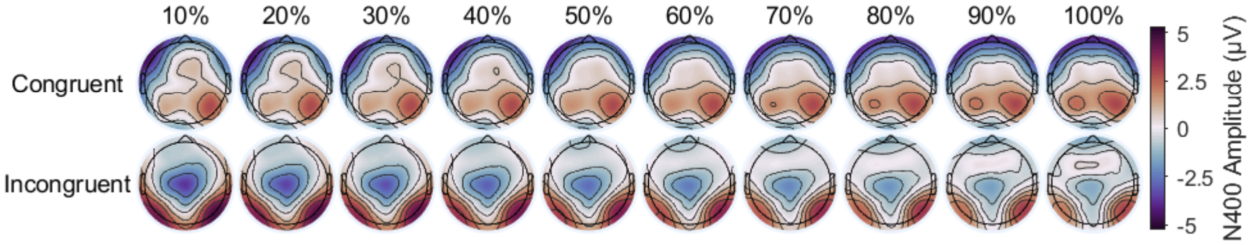

Estimates are shown for picture-congruent and -incongruent words separately, at levels of predictability from 10 to 100%.

Results showed that picture-incongruent words elicited a clear N400 difference from picture-congruent words, and that this difference was largest at low levels of Predictability, and smallest at high levels of Predictability.

#### K Checking ERPs Time-Locked to Pictures

We conducted an exploratory analysis of the ERPs observed when the evoked signals are time-locked to the onset of pictures that preceded words. The reasoning behind this analysis was twofold. First, we considered that this may reveal an effect of Predictability (Cheng et al., 2010), since name agreement is a feature of the picture stimuli in isolation, as well as of picture-word pairs. Such an effect would have affected the baseline period of the word ERP, and may have influenced estimates of Predictability. Second, we wanted to check that there were no effects of Congruency prior to word presentation. Because we presented words *after* pictures, we reasoned that there should be no effect of Congruency or the Congruency-Predictability interaction in the ERPs elicited by pictures.

To examine picture-elicited ERPs, we fit linear mixed-effects models like those described in the section, *Exploratory Scalp-Wide Analysis*, of the manuscript, but with ERPs time-locked to picture presentation rather than word presentation.

Results (**Figure 23**) revealed an ERP with high positivity over posterior regions. As sometimes seen in ERPs elicited by images (e.g., Cheng et al., 2010; Kaufmann et al., 2009; Rousselet et al., 2004; Weinberg & Hajcak, 2010), the N1 deflection was so small, relative to the P1, that posterior amplitudes remained positive throughout. Consistent with findings reported by Cheng et al. (2010), we found evidence of predictability (i.e., name agreement) effects on the picture ERP. We note, however, that Cheng et al. (2010) found evidence for more positive amplitudes across most of the scalp and time-course. In contrast, we found evidence for more negative amplitudes at occipitotemporal sites, and, conversely, more positive amplitudes in central and frontal locations. By the time of stimulus presentation, the effect of predictability showed a very noisy pattern of differences between midline and peripheral electrodes, similar in magnitude to effects of Congruency and the Congruency-Predictability interaction estimated for the same time point. As expected, we observed no clear effects of Congruency, or of the Congruency-Predictability interaction, on the picture ERP.

**Figure 22**

*Time-course of scalp-wide fixed-effects estimates for picture ERPs.*

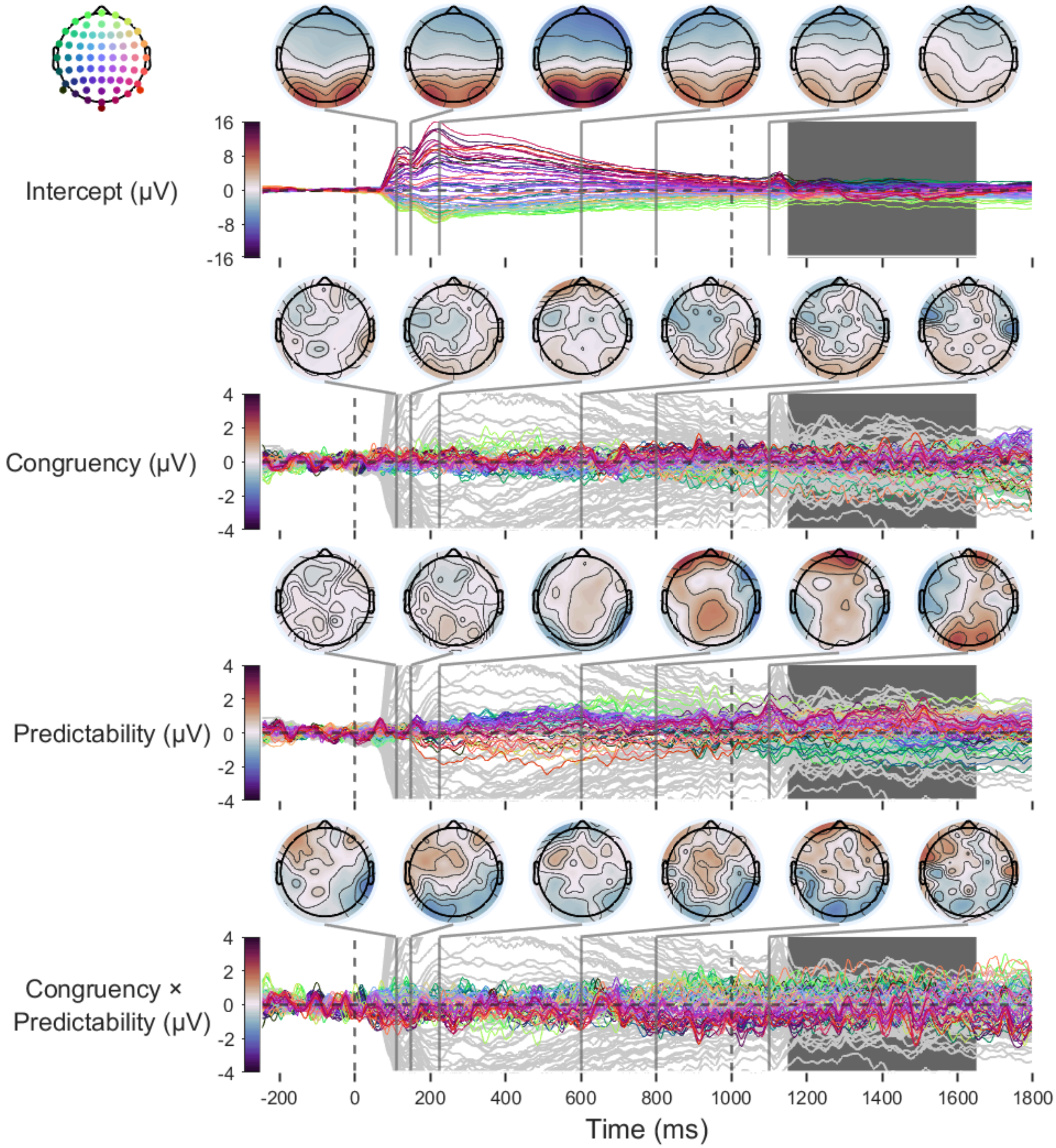

The first dashed vertical line (0 ms) indicates stimulus (picture) onset. The second dashed vertical line (1000 ms) indicates the time-point at which the picture was replaced with a fixation point. The large dark grey period from 1150 to 1650 ms reflects the jittered period during which words were presented. Topographic plots of fixed effects are highlighted at key time-points. Model intercepts (reflecting amplitudes at the lowest level of Predictability) are depicted as grey lines on each panel to provide a reference for timing and magnitude of effects. Note that axis limits for the fixed-effect slopes are reduced by four times for visibility of any small effects.

Although the estimates are noisy, we consider it possible that a sustained effect of Predictability influenced the ERPs elicited by words. Indeed, such an effect may even index preactivation processes that underlie prediction effects on the word N1. Nevertheless, the Congruency-Predictability interaction we used to test our hypotheses can only have emerged after word onset.

#### L Checking ERPs Time-Locked to Responses

We considered that ERPs elicited by words may have also included signals associated with participants preparing to initiate their response. To examine this we calculated ERPs time-locked to participants' button presses, via linear mixed-effects models like those described in the section, *Exploratory Scalp-Wide Analysis*, of the manuscript. However, as we were interested in ERPs prior to the event, we did not include any baseline correction. Results showed that, beginning around 400 ms prior to response, picture-incongruent words elicited more negative-going centroparietal amplitudes. Given that the colour change, after which participants could respond, occurred at 500 ms, this effect is likely to reflect the large and long-lasting N400 effect of Congruency observed in the word ERPs.

The effect of Predictability, and the Congruency-Predictability interaction, did not show clear effects in the ERP time-locked to responses. Effects of these variables on the word ERP are likely not reflected in the response ERP because, relative to the N400 effect, effects of Predictability and the Congruency-Predictability interaction were smaller and more transient. As a result, these effects are averaged out by variability in response times, which introduces jitter to effects elicited by words.

**Figure 23**

*Time-course of scalp-wide fixed-effects estimates for Response ERPs.*

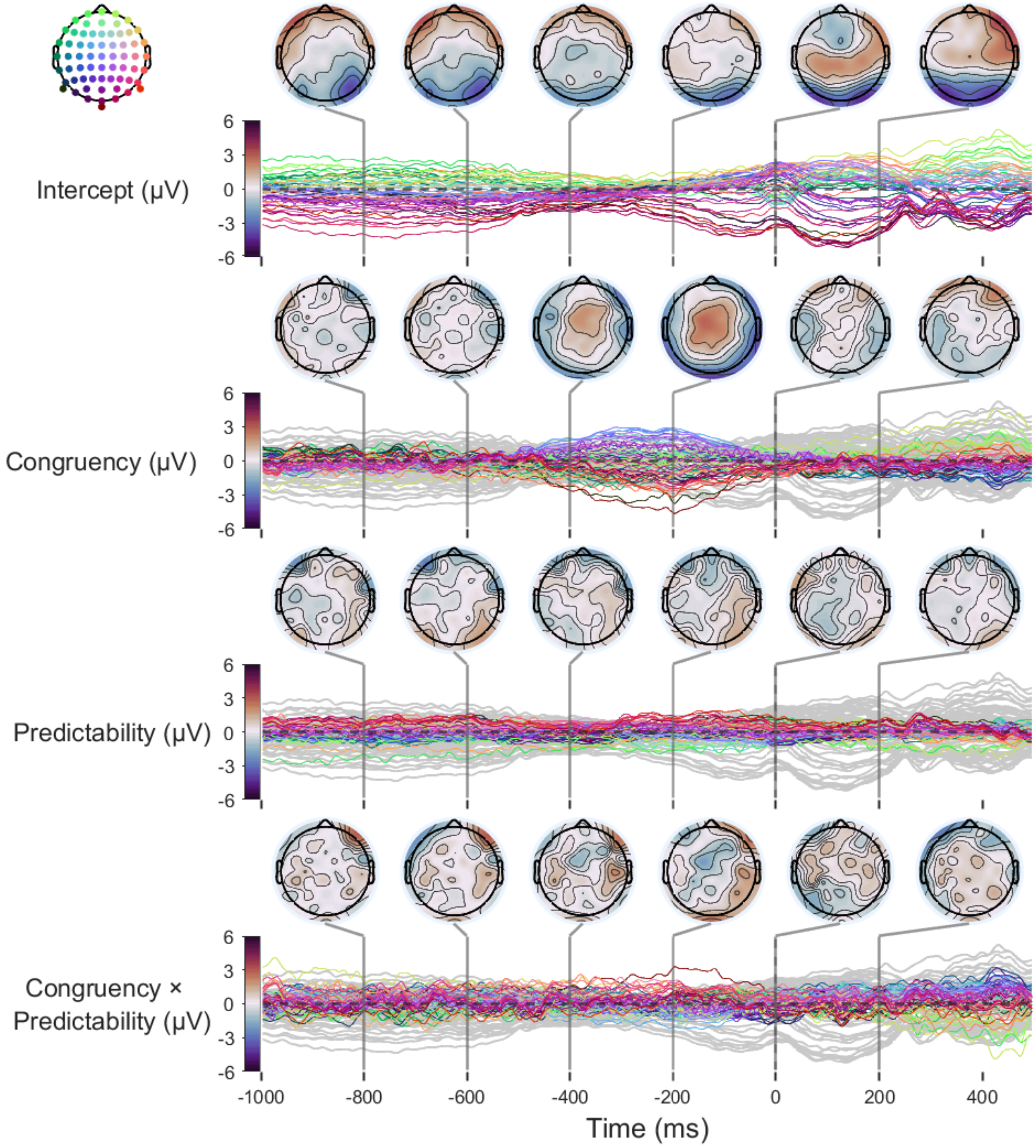

Model intercepts (reflecting amplitudes at the lowest level of Predictability) are depicted as grey lines on each panel to provide a reference for timing and magnitude of effects. Note that axis limits for the fixed-effect slopes are reduced by four times for visibility of any small effects.
